# An improved *hgcAB* primer set and direct high-throughput sequencing expand Hg-methylator diversity in nature

**DOI:** 10.1101/2020.03.10.983866

**Authors:** Caitlin M. Gionfriddo, Ann M. Wymore, Daniel S. Jones, Regina L. Wilpiszeski, Mackenzie M. Lynes, Geoff A. Christensen, Ally Soren, Cynthia C. Gilmour, Mircea Podar, Dwayne A. Elias

## Abstract

The gene pair *hgcAB* is essential for microbial mercury methylation. Our understanding of its abundance and diversity in nature is rapidly evolving. In this study we developed a new broad-range primer set for *hgcAB*, plus an expanded *hgcAB* reference library, and used these to characterize Hg-methylating communities from diverse environments. We applied this new Hg-methylator database to assign taxonomy to *hgcA* sequences from clone, amplicon, and metagenomic datasets. We evaluated potential biases introduced in primer design, sequence length, and classification, and suggest best practices for studying Hg-methylator diversity. Our study confirms the emerging picture of an expanded diversity of HgcAB-encoding microbes in many types of ecosystems, with abundant putative mercury methylators *Nitrospirae* and *Chloroflexi* in several new environments including salt marsh and peat soils. Other common microbes encoding HgcAB included *Phycisphaerae, Aminicenantes, Spirochaetes*, and *Elusimicrobia*. Gene abundance data also corroborate the important role of two “classic” groups of methylators (*Deltaproteobacteria* and *Methanomicrobia*) in many environments, but generally show a scarcity of *hgcAB*+ *Firmicutes*. The new primer set was developed to specifically target *hgcAB* sequences found in nature, reducing degeneracy and providing increased sensitivity while maintaining broad diversity capture. We evaluated mock communities to confirm primer improvements, including culture spikes to environmental samples with variable DNA extraction and PCR amplification efficiencies. For select sites, this new workflow was combined with direct high-throughput *hgcAB* sequencing. The *hgcAB* diversity generated by direct amplicon sequencing confirmed the potential for novel Hg-methylators previously identified using metagenomic screens. A new phylogenetic analysis using sequences from freshwater, saline, and terrestrial environments showed *Deltaproteobacteria* HgcA sequences generally clustered among themselves, while metagenome-resolved HgcA sequences in other phyla tended to cluster by environment, suggesting horizontal gene transfer into many clades. HgcA from marine metagenomes often formed distinct subtrees from those sequenced from freshwater ecosystems. Overall the majority of HgcA sequences branch from a cluster of HgcAB fused proteins related to *Thermococci, Atribacteria* (candidate division OP9), *Aminicenantes* (OP8), and *Chloroflexi*. The improved primer set and library, combined with direct amplicon sequencing, provide a significantly improved assessment of the abundance and diversity of *hgcAB*+ microbes in nature.

**Contribution to the Field Statement:** The gene pair *hgcAB* is essential for microbial production of the neurotoxin methylmercury. In recent years these genes have been used as biomarkers to determine the potential of a microbiome to generate methylmercury via PCR amplification using degenerate primers from several research groups. However, improved techniques for capturing *hgcAB* diversity are necessary for identifying the major environmental producers of the neurotoxin as well as the expanding diversity of novel putative methylators, and the genes’ evolutionary history. The work described herein advances *hgcAB* detection in environmental samples through an updated primer set coupled with a direct high-throughput sequencing method that enables broader diversity capture. We provide an expanded *hgcAB* sequence reference library that allows for more sensitive and robust estimations of Hg-methylator diversity and potential for MeHg generation in the environment. The *hgcAB* diversity generated by high-throughput sequencing confirms the potential for novel Hg-methylators previously only identified using metagenomic screens. This study provides a significantly improved assessment of the abundance and diversity of *hgcAB*+ microbes in nature. By expanding our understanding of the microbial metabolic clades associated with mercury methylation, this work improves our ability to predict environmental conditions that drive production and accumulation of the neurotoxin in aquatic ecosystems.

## 1 Introduction

Microorganisms influence the environmental form of mercury (Hg), including the production and degradation of the bio-accumulative neurotoxin methylmercury (MeHg) (Hsu-Kim et al., 2013). We still know relatively little about the distribution of Hg-methylating organisms in the environment, although the discovery of the *hgcAB* gene pair conferring Hg-methylation capability has greatly improved their identification (Parks et al., 2013; Bravo and Cosio, 2019). Mercury methylation is a species (and possibly strain)-specific trait that is not well predicted by 16S rRNA taxonomic classification (Ranchou-Peyruse et al., 2009; Gilmour et al., 2013; Yu et al., 2013; Gilmour et al., 2018; Christensen et al., 2019). The presence or abundance of groups of microbes that include Hg methylators is not predictive of the abundance of *hgcAB*^+^ microbes. Thus, identification of *hgcAB* sequences is required to evaluate the distribution of Hg-methylators in nature. The known diversity of Hg methylators is expanding as increasing numbers of *hgcAB* sequences are identified from shotgun metagenomics, metagenome-assembled genomes (MAGS), and *hgcAB* amplicon-specific sequencing.

In addition to the Hg-methylators confirmed in culture, sulfate- and iron-reducing *Deltaproteobacteria*, acetogenic and syntrophic *Firmicutes*, and methanogenic *Archaea* (Compeau and Bartha, 1985; Fleming et al., 2006; Kerin et al., 2006; Ranchou-Peyruse et al., 2009; Gilmour et al., 2013; Yu et al., 2013; Gilmour et al., 2018; Goñi-Urriza et al., 2019)-genomic analyses have identified *hgcAB* (and thus inferred the ability to produce MeHg) in many other lineages including *Chloroflexi, Chrysiogenetes, Aminicenantes* (Candidate Phylum OP8), *Atribacteria* (Candidate Phylum OP9), *Nitrospira, Nitrospina, Elusimicrobia, Thermotogae* and *Spirochaetes* (Podar et al., 2015a; Gionfriddo et al., 2016; Liu et al., 2018a; Bowman et al., 2019; Christensen et al., 2019; Jones et al., 2019). The occurrence of *hgcAB* in Bacterial and Archaeal genomes is rare but widely dispersed across phyla (Parks et al., 2013; Podar et al., 2015a; Jones et al., 2019). Hg-methylating microorganisms often constitute less than 5% of the bulk microbial community (Christensen et al., 2019), yet in these same environments MeHg can comprise up to 10% of the total Hg within a system (Christensen et al., 2019). Since MeHg readily biomagnifies up aquatic food webs, it poses a significant risk to ecosystem and public health even at picomolar concentrations in the water column and sediments (Chen et al., 2014; Lee and Fisher, 2017).

PCR primers and reference libraries for *hgcAB* have improved with the increasing number of *hgcAB* sequences identified. Early studies relied on sequences from a small number of cultured methylators (Bae et al., 2014; Liu et al., 2014; Schaefer et al., 2014) while more recent work has captured a wider diversity (Christensen et al., 2016; Bravo and Cosio, 2019; Christensen et al., 2019; Jones et al., 2019). Improved techniques for capturing *hgcAB* presence and diversity are necessary for identifying both the major players in environmental MeHg production as well as the evolutionary history of the gene pair.

Here we report an improved PCR primer set, reference library and amplification protocol for *hgcAB* detection from environments significant to Hg biogeochemical cycling. The new primer set is a modification of the Christensen broad-range *hgcAB* primer set, which was designed using a database of 84 known and predicted Hg-methylator genomes (Christensen et al., 2016). While the highly divergent primers amplified *hgcAB* from many types of pure cultures, amplification efficiencies from natural environments were often low (Christensen et al., 2019). PCR primer degeneracy is an important tool for capturing genetic diversity while minimizing biases (von Wintzingerode et al., 1997; Sipos et al., 2007), but high levels of degeneracy can reduce amplification of individual primer matches.

To improve amplification across clades, we modified the reverse primer to explicitly target only *hgcAB* sequences found in nature. This change significantly reduced primer degeneracy while maintaining the ability to capture broad *hgcAB* sequence diversity. Use of this primer set limits non-specific amplification, resulting in increased primer efficiency. To improve detection from complex environments, we also optimized sample volumes and concentrations to mitigate PCR inhibition, using several sample matrices. The updated primer and PCR method were validated using mock communities and culture spikes into sediments and soils. Additionally, an expanded *hgcAB* reference library was developed based on a larger number of sequences from confirmed *hgcAB*^+^ cultures plus sequences from publicly available MAGS for a total of 283 sequences.

Finally, the new primer, the protocol, and the reference library were used to evaluate the community structure of HgcAB-encoding microbes from several novel environments, including salt marsh soils, freshwater sediments and peat soils. Hg methylation genes were evaluated using both clone libraries and high-throughput amplicon-specific sequencing. The use of high-throughput amplicon sequencing eliminates the need to create clone libraries and very significantly increases the efficacy of identifying *hgcAB*^+^ organisms. A new phylogenetic analysis of HgcA was constructed using the deep sequencing data from fresh and estuarine sediments of two of the study sites.

## 2 Materials & Methods

### 2.1 Expanded reference database of *hgcAB* sequences

An expanded library was constructed based on confirmed Hg-methylators plus sequences from single cell genomes and MAGs from diverse environments (Table S1). The new *hgcA* library contains 283 *hgcA* sequences, including 124 *Deltaproteobacteria*, 33 *Firmicutes*, 15 *Methanomicrobia*, and 111 sequences from other clades (Gionfriddo et al., 2019). Of those sequences, 239 had an available *hgcB* sequence. This custom database of known and predicted Hg-methylators with NCBI taxonomy was compiled on the 16^th^ of October 2018 using Taxtastic (v0.8.5, https://github.com/fhcrc/taxtastic) (Sayers et al., 2009). The compiled Hg-methylator reference database is maintained by the ORNL Mercury SFA and is publicly available through the DOE Data Explorer (Gionfriddo et al., 2019). This database expands the *hgcAB* sequence library substantially over most libraries used in *hgcAB* classification to date (Podar et al., 2015b; Bravo et al., 2018b; Liu et al., 2018b).

The ORNL compiled Hg-methylator database can be used to assign taxonomic classification to HgcA sequences from clone, amplicon, or meta-omic datasets (Gionfriddo et al., 2019). The database contains reference *hgcA* and *hgcB* sequences, and two reference packages for identifying and classifying longer (654 nt bp) or shorter (201 nt bp) *hgcA* sequences. Each reference package contains an HgcA hmm-model and HgcA phylogenetic tree. To construct the hmm-models, nucleotide *hgcA* sequences were aligned (MUSCLE), trimmed (201 bp or 654 bp), translated into amino acids using Geneious version 10.2, and compiled into an hmm-model using ‘hmmbuild’ (Edgar, 2004b; Johnson et al., 2010; Kearse et al., 2012). A maximum likelihood (ML) HgcA distribution was estimated using the GAMMA model of rate heterogeneity and ML estimate of alpha-parameter in RAxML with 150 bootstrap replicates. The tree was rooted by carbon monoxide dehydrogenases from non-Hg-methylators *Candidatus Omnitrophica bacterium* CG1-02-41-171 and *Thermosulfurimonas dismutans* (Stamatakis, 2014). These paralogous sequences with distinct taxonomy from Hg-methylators were included in the database to identify non-HgcA encoding sequences that may have passed previous filtering.

### 2.2 Primer design

Our goal in creating a revised broad-diversity primer set for *hgcAB* was to increase the specificity and efficiency of the previously published primer set (Christensen et al., 2016) while retaining the ability to capture the full diversity of the community. The Christensen 2016 primer set (ORNL-HgcAB-uni-F & ORNL-HgcAB-uni-R) was designed based on the *hgcAB* sequences from 84 strains that were available at that time, including *Deltaproteobacteria, Firmicutes, Archaea*, and a limited number of other clades. The primers target the highly conserved cap-helix encoding region of *hgcA* and the highly conserved ‘C(M/I)ECGA’ ferredoxin-type encoding domain of *hgcB* (Parks et al., 2013), amplifying a ∼950 nt bp product. To capture the diversity of the reference strains, the Christensen primers (especially the reverse primer) are highly (96-fold) degenerate. When used in combination with a diverse reference library for OTU identification, these primers capture a highly diverse community of *hgcAB*+ organisms, including many groups outside of the three used to construct the primers (Christensen et al., 2019; Jones et al., 2019).

Our strategy was to reduce the overall degeneracy of the reverse (*hgcB*) primer by eliminating oligonucleotide sequences from the primer mix that are not found in nature. Lower degeneracy should result in more efficient amplification, and thus recover more *hgcAB* sequences from the environmental samples. Only the reverse primer sequence, not position, was updated.

The 96 reverse primer sequences that make up ORNL-HgcAB-uni-R (Table S2) were screened against the expanded *hgcAB* reference library. Only 39 of the possible 96 sequences in ORNL-HgcAB-uni-R degenerate primer matched sequences in the expanded reference library. In the reverse primer-binding region, seven of twenty bases show 100% pairwise identity, twelve are fairly conserved (30%-100% sequence identity), and one is highly divergent (<30% pairwise identity) across the available *hgcB* reference sequences (Figure 1). To choose the final primer, degeneracy was reduced at the 5’ end and increased at the 3’ end to combat primer mismatch and binding efficiencies. Mismatches within five base pairs (nt bp) of the 3’ end can have larger effects on primer efficiency than 5’ located mismatches (Stadhouders et al., 2010; Campos and Quesada, 2017). Therefore, increasing the possible primer sequences at the 3’ end increases the likelihood of exact matches, theoretically improving binding specificity (Stadhouders et al., 2010). Changing, from the 5’ end, the third base B = C/T/G to G, the ninth base from R=G/A to ‘G’, and increasing the 15^th^ base degeneracy from ‘C’ to S = C/G decreased the degeneracy from 96 to 32 and also resulted in eliminating rare *hgcB* reference sequences (Figure 1). To summarize, the reverse primer sequence was changed from ‘CA**B** GCN CC**R** CAY TC**C** ATR CA’ to ‘CA**G** GCN CC**G** CAY TC**S** ATR CA’ (differences shown in bold). The new primer is designated ORNL-HgcAB-uni-32R, signifying the 32-fold degeneracy.

**Figure 1.**
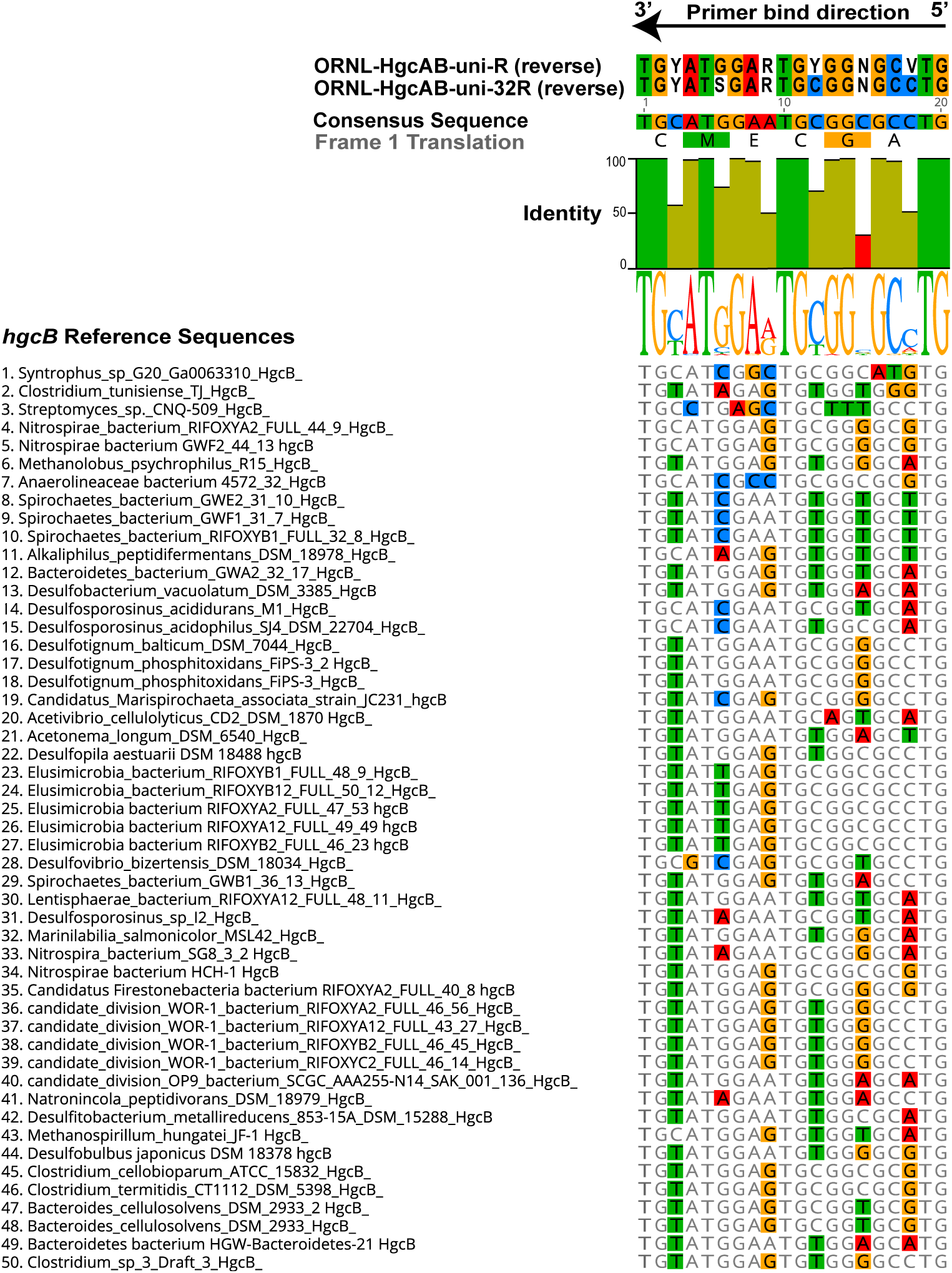
Alignment showing the highly conserved 20 nt bp region of *hgcB* where the reverse primer binds. Alignment is based on 239 reference *hgcB* sequences, however only the top 50 sequences most divergent from the consensus sequence are shown. Bases that disagree with consensus sequence are highlighted. Reverse complements of each primer (ORNL-HgcAB-uni-R, ORNL-HgcAB-uni-32R) are included for reference. The bar graph at the top of the image shows mean pairwise identity for each base across all sequences – green is 100%, green-brown is between 30 and 100%, and red is <30% conserved in 239 sequences. Y = C/T, R = G/A, V = C/G/A, S = C/G, N = A/T/C/G.

*In-silico* primer match efficiency and specificity was examined using the Geneious version 10.2 ‘test primer’ function, allowing zero to two mismatches (http://www.geneious.com) (Kearse et al., 2012). Mismatches within five nucleotides of the 3’ end were not allowed. Efficiency is defined as whether the primer sequence matched HgcB-encoding sequences from known and predicted methylators. Reverse primers were tested for specificity against a reference database of publicly available nucleotide sequences encoding 4Fe-4S ferredoxins (i.e. non-HgcB encoding paralogs) from JGI (https://img.jgi.doe.gov/). Specificity is defined as whether the primer sequence would theoretically bind to non-HgcB encoding sequences. The reference set contained a total of 14,161 nucleotide sequences from environmental metagenomes.

For initial validation, ORNL-HgcAB-uni-32R was tested on genomic DNA (gDNA) from 31 pure culture strains, (26 *hgcAB*+ and 5 *hgcAB-* controls) (Christensen et al., 2016). Premixed and individual oligonucleotides were ordered from Integrated DNA Technologies and the latter were mixed in-house at equimolar concentrations. The annealing temperatures for the sequences in the 96-fold degenerate primer (ORNL-HgcAB-uni-R) ranged from 56.1–68.2 °C, compared to 60.2– 68.5 °C for ORNL-HgcAB-uni-32R. It is used with the published forward primer (ORNL-HgcAB-uni-F, 5’ AAYGTCTGGTGYGCNGCVGG). Annealing temperatures for the forward primer ranged from 60.9–70.7 °C.

### 2.3 Sample sites and DNA extraction

Environmental samples were obtained from Oak Ridge National Laboratory (ORNL) collaborative sites including peatland sediments from Spruce and Peatland Responses Under Changing Environments (SPRUCE) study plots in the USDA Forest Service Marcell Experimental Forest (MEF) (Grand Rapids, MN), freshwater sediments from Sandy Creek (Durham, NC), mesohaline tidal salt marsh sediments from Rhode River (Edgewater, MD), periphyton and sediment from East Fork Poplar Creek (EFPC) (Oak Ridge, TN), and rice paddy soil from Yanwuping (Guizhou Province, China). Sample collection information, including location and date of collection are summarized in Table S3. SPRUCE sediments were collected from treed hummock (2012-10-T-Hum; 2012-6-T-Hum) and hollow (2012-10-T-Hol, 2012-6-T-Hol) sites of the S1-Bog from surface to 200-300 cm (Iversen et al., 2012). The EFPC sediment and periphyton samples were collected from the New Horizon (NH) study site (35.9662°N, 84.35817°W). Total Hg (HgT) concentrations ranged ∼20–50 mg kg^−1^ (Southworth et al., 2010). EFPC water ranged from 110–211 ng Hg L^−1^ (unfiltered) and 15–89 ng L^−1^ (filterable), respectively (Olsen et al., 2016). Total and filterable methylmercury (MeHg) ranged from 0.15– 0.43 ng L^−1^ and 0.10–0.29 ng L^−1^, respectively (Olsen et al., 2016). Mesohaline tidal salt marsh sediments were collected from a *Phragmites*-dominated area of the Global Change Research Wetland on the Rhode River (Edgewater, Maryland, USA), where soil Hg concentrations averaged 0.17 ± 0.007 mg kg^−1^ Hg (Mitchell and Gilmour, 2008). Sandy Creek sediments were collected from a freshwater retention pond (Durham, NC, USA) that receives legacy Hg, with sediments averaging 0.05 ± 0.006 mg kg^−1^ Hg (Ndu et al., 2018). Rice paddy soils were collected during the rice growing season from Yanwuping in Guizhou province, China. Rice paddy fields were 2–14.5 km downstream from a historical Hg mining site, with HgT levels ranging from 0.83–990 mg kg^−1^ and MeHg from 0.0038–0.021 mg kg^−1^ (Vishnivetskaya et al., 2018).

Sediment, soil, and periphyton samples were stored at −80°C prior to DNA extraction. Genomic DNA (gDNA) was isolated from sediments using the MoBio Powersoil kit (now Qiagen DNeasy) with the addition of a 30-minute incubation at 65°C following the bead beating step. A standard phenol chloroform extraction was used to isolate gDNA from periphyton biomass. All gDNA was quantified using Qubit™ (Thermo Fisher Scientific) and quality was assessed with NanoDrop™ One (Thermo Fisher Scientific).

### 2.4 Mock communities

Pure cultures of known Hg-methylators were spiked into fresh homogenized Rhode River tidal salt marsh soil sample #1064 (∼0.2 g) to form mock community samples. Two mock communities were constructed. For Combo 1, sediment was spiked with 1.00E9 cells *Geobacter (G.) sulfurreducens* (∼3800 ng DNA), 4.93E8 cells *Desulfosporosinus (Ds.) acidophilus* (2700 ng DNA), and 1.01E9 cells *Methanocorpusculum (Mc.) bavaricum* (1900 ng DNA). Combo 2 was spiked with 6.34E8 cells *Desulfovibrio (Dv.) desulfuricans* ND132 (2700 ng DNA), 1.13E8 cells *Desulfitobacterium (Db.) dehalogens* (540 ng DNA), and 4.10E8 cells *Methanomassiliicoccus (Mm.) luminyensis* (1200 ng DNA). DNA was recovered using a modified MoBio Powersoil kit. To estimate DNA recovery, extraction efficiencies were evaluated from individual cultures alone and spiked into the marsh soil. Average extraction efficiencies for cultures spiked into sediment were similar to efficiencies from cultures alone, ∼30%, but with significant variability among cultures. For the mock communities, DNA recoveries for spiked cultures were 10 ± 4.8% and 37 ± 6% respectively from Combo 1 and 2.

### 2.5 PCR amplification of 16S rRNA and hgcAB

For environmental samples, both 16S rRNA and *hgcAB* genes were amplified. For 16S rRNA amplification, the 25-µL reaction mixture contained 0.2 µM of 8F (5’ AGA GTT TGA TCC TGG CTC AG) and 1492R (5’ GGT TAC CTT GTT ACG ACT T) (Turner et al., 1999), Apex TaqRed polymerase (Genesee Scientific, San Diego, CA), and a dilution series of gDNA template (10 pg–10 ng) to determine the optimal gDNA concentration and ensure amplification. Amplification was initiated with pre-incubation (2 min at 98°C), 30 cycles of denaturation (15s at 98°C), annealing (30s at 54°C), and extension (60s at 72°C), then a final incubation (3 min at 72°C).

For *hgcAB* amplification from both pure culture and environmental samples, the use of a polymerase without a 3’ to 5’ exonuclease proofreading activity is important. In our experience, a proofreading polymerase that uses 3’ to 5’ exonuclease, such as a Pfu polymerase, leads to multiple visible gel bands of non-specific PCR product. For *hgcAB* amplification, 20 and 50-µL reaction mixtures used 0.5–1 µM of ORNL-HgcAB-uni-F and ORNL-HgcAB-uni-32R, Apex TaqRed polymerase (Genesee Scientific, San Diego, CA), and 100 pg to 10 ng of gDNA. A touch-down PCR protocol was used with pre-incubation (2 min at 98°C), then 5 cycles of denaturation (30s at 98°C), annealing (30s at 68°C; −1 °C/cycle), and extension (30s at 72°C), then 30 cycles of denaturation (30s at 98°C), annealing (30s at 63°C), and extension (60s at 72°C), then a final incubation step (2 min at 72°C).

### 2.6 Clone libraries

Clone libraries of *hgcAB* were constructed from all of the environmental samples listed in Table S3. PCR conditions were 0.5 ng/ µL (final concentration) gDNA template and 0.5 µM primer in 20 µL reaction volume. Clone libraries were created from both the ORNL-HgcAB-uni-F/ ORNL-HgcAB-uni-R and the ORNL-HgcAB-uni-F/ ORNL-HgcAB-uni-32R primer pairs for comparison.

The number of clones created for each sampling area ranged from 12 to >100 (Table S3). The environmental clone libraries contained 509 *hgcAB* sequences amplified with the ORNL-HgcAB-uni-F/ ORNL-HgcAB-uni-32R primers and 119 with ORNL-HgcAB-uni-F/ ORNL-HgcAB-uni-R primers. To create the libraries, *hgcAB* PCR products (∼950 bp) from each sample were excised from an agarose gel, cleaned with Wizard SV Gel and PCR Clean-Up system (Promega, Madison, WI) and cloned into One Shot TOP10 cells with TOPO-TA cloning kit (Invitrogen, Waltham, MA). Cloned plasmids were screened for the ∼1150 bp insert using 100 pM of M13F (5’ GTAAAACGACGGCCAG) and M13R (5’ CAGGAAACAGCTATGAC) primers with Apex TaqRed polymerase. The insert size was confirmed (on 1% agarose gel), cleaned (Wizard SV Gel and PCR Clean-Up) and sent for Sanger sequencing by Eurofins Genomics (Louisville, KY).

For classification of the clone sequences, the following pipeline was used. All *hgcAB* sequences from the clones were confirmed by blastx against the NCBI non-redundant database (Altschul et al., 1990) and non-specific products were removed from further analysis. The resulting sequences were aligned, trimmed, and primers annotated using Geneious version 10.2 (http://www.geneious.com) (Kearse et al., 2012). The trimmed *hgcAB* sequences were aligned using MUSCLE (Edgar, 2004a) to a reference database of 296 *hgcA* and 200 *hgcB* sequences from known and predicted Hg-methylators (Table S1). While both genes were present in cloned sequences, only the *hgcA* portion of the sequence was used for downstream analyses. This is due to both the discrepancy in the number of available reference sequences between *hgcA* and *hgcB*, as well the difficulty in aligning sequences across the portion of DNA between the protein-coding regions of *hgcA* and *hgcB*.

We performed taxonomic classification of clone sequences using our custom in-house short-read HgcA reference package (‘ORNL_HgcA_201.refpkg’) and longer-read HgcA reference package (‘ORNL_HgcA_654_full.refpkg’) (Gionfriddo et al., 2019). In order to compare to the shorter amplicon sequences, clone *hgcA* sequences were trimmed to 201 nt bp and classified following the same protocol as amplicon sequences. In short, clone *hgcA* sequences were clustered using ‘cd-hit-est’ on a 90% sequence identity cut-off over a 201 nt bp region of *hgcA*. Taxonomic classifications were assigned to OTUs based on phylogenetic placement of translated HgcA sequences on HgcA reference tree (in ‘ORNL_HgcA_201.refpkg’) using ‘pplacer’ and ‘guppy classify’ with posterior probability classification cut-off of 90% (Matsen et al., 2010). To compare how our classification pipeline performed on short (201 nt bp) versus longer (654 nt bp) *hgcA* sequences, we classified the same clone sequences, now trimmed to 654 nt bp, using ‘ORNL_HgcA_654_full.refpkg’. The longer-read reference package contains reference HgcA sequences encoded by 654 nt bp, corresponding to the 268–955 nt bp region of *D. desulfuricans* ND132 *hgcA* (Genbank CP003220.1, 1150554:1151240). The environmental clone *hgcA* sequences from this study are publicly available under the NCBI GenBank accession numbers MT122211 - MT122744.

### 2.7 High-throughput sequencing of *hgcAB* amplicons

For environmental samples from the SERC salt marsh and the NH site in EFPC, and for select mock communities (Table S4), *hgcAB* sequence abundance and diversity were also evaluated by high-throughput amplicon-specific sequencing, using Illumina MiSeq 2×300 bp chemistry at the University of Minnesota Genomics Facility (UMGC). A two-step PCR amplified *hgcAB* from environmental and mock community gDNA by adapting published protocols (Gohl et al., 2016; Jones et al., 2017). The modified *hgcAB*-specific primers included 5’ adaptor sequences (adaptor TCG TCG GCA GCG TCA GAT GTG TAT AAG AGA CAG with ORNL-HgcAB-uni-F; adaptor GTC TCG TGG GCT CGG AGA TGT GTA TAA GAG ACA G with ORNL-HgcAB-uni-R and ORNL-HgcAB-uni-32R). Amplicon libraries were created by amplifying *hgcAB* with the modified primers for 30 or 35 cycles (Table S4) following the PCR protocol for *hgcAB* cloning described above as in Jones et al. (2017). Illumina sequencing adaptors and dual indices were added at UMGC with a second amplification step (10 cycles, after diluting PCR products 1:100) (Gohl et al., 2016). Libraries were pooled and sequenced on an Illumina MiSeq v3 PE300. Read length was not sufficient to produce an overlap between forward and reverse reads, therefore two separate libraries were produced, the forward reads encompassing approximately the first 300 bp of *hgcA* starting at the cap-helix and reverse reads encompassing a 300 bp portion of *hgcB* and *hgcA*. The presence of both genes (*hgcA* and *hgcB*) is needed for PCR amplification. The environmental and mock community *hgcAB* amplicon raw sequence files are publicly available at the NCBI SRA accession (PRJNA608965).

Raw sequences were filtered and trimmed using Trimmomatic, and only forward *hgcA* reads were used (Bolger et al., 2014). We adapted OTU clustering methods applied to functional genes from Bravo et al. (2018a) and Pelikan et al. (2016). For this study, *hgcA* sequences were trimmed (201 nt bp), dereplicated, and singletons removed with USEARCH (Edgar, 2010). Chimeras were removed using de-novo and reference filtering with VSEARCH (Rognes et al., 2016). For NH *hgcAB*, 88.2–91.7% of amplicons from the old primer set and 80.8–90.5% of amplicons from the new primer set passed quality and chimeric filtering. The non-chimeric dereplicated sequences were clustered at 90% identity using cd-hit-est (Huang et al., 2010), and trimmed forward reads were mapped to the centroid database using ‘-usearch_global’ to produce OTU counts.

The sequencing error rate was calculated for both primer-sets using the quality filtered and trimmed combo 1 and combo 2 mock community datasets using the ‘seq.error’ function in mothur (Schloss et al., 2009). The sequencing error for combo 1 mock community amplified with the new primer set was 1.94%, while with the old published primer set was 2.20%. The sequencing error for combo 2 mock community amplified with the new primer set was 2.05%, while with the old published primer set was 2.27%.

### 2.9 Taxonomic classification of amplicon HgcA sequences

The Taxtastic reference package ‘ORNL_HgcA_201.refpkg’ was used to assign taxonomy to the short (201 nt bp) centroid sequences. The nucleotide sequences are first translated and then aligned to reference sequences using ‘transeq’ and ‘hmmalign’, respectively (Johnson et al., 2010; Li et al., 2015). Only forward sequences where the first base was the start codon were kept after translation. Prior to alignment, sequences with stop codons were removed, and reads with <1e-7 inclusion value with the HgcA cap-helix region were filtered using ‘hmmsearch’ (Johnson et al., 2010). Pplacer placed the filtered, translated, and aligned centroid sequences on the ML reference tree. Taxonomic classifications were assigned to each centroid sequence based on the lowest common ancestor (LCA) of the subtree where the sequence was placed using ‘guppy classify’ with a classification cutoff of 90% (Matsen et al., 2010). To avoid misclassifications of distantly related sequences, only placements with a branch length <1 were classified. OTU assignments, phylogenetic placements of centroid sequences, and taxonomic classifications were analyzed in R using ‘BoSSA’ (v3.4, https://cran.r-project.org/web/packages/BoSSA/) and ‘phyloseq’ (McMurdie and Holmes, 2013).

In the NH *hgcA* dataset, 86.0–96.1% of filtered non-chimeric amplicons from the old primers and 69.0–92.7% of amplicons from the new primers could be classified. The top ten OTUs from both primer-sets that did not pass filtering criteria contained sequences with high similarity to *hgcA* and *hgcB* outside the 201 nt bp *hgcA* region as well as non-*hgcAB* genes that encode 4Fe-4S ferredoxins with reverse primer-binding sites when allowing up to 3 mismatches (NADH:ubiquinone oxidoreductase, xanthine dehydrogenase) and genes that encode proteins that do not appear to contain primer-binding sites even when allowing for mismatches (ATPases, diguanylate cyclase, threonine/serine dehydratase, O-methyltransferase, proteins with unknown function).

For comparison, the 50 most abundant OTUs from NH amplicon libraries were classified based on LCA (cut-off 50%) taxonomy from protein BLAST against NCBI non-redundant protein database using MEGANv6 (Altschul et al., 1990; Huson et al., 2016) (Figure S1). The closest BLAST matches were often ‘unclassified’ sequences (Figure S1). Therefore, only *hgcAB* sequences from genomic data with accompanying phylogeny were included in the reference database. As a direct comparison, publicly available rice paddy metagenomic reads were downloaded from the NCBI sequence archive (Accession: PRJNA450451) (Liu et al., 2018b). The *hgcA* genes were identified in sequence reads and aligned to reference sequences using the hmm model from the ‘ORNL_HgcA_201.refpkg’ using ‘hmmsearch’ and ‘hmmalign’ (inclusion value 1E-7). Taxonomic classifications were assigned to *hgcA* reads using pplacer with ‘ORNL_HgcA_201.refpkg’ with the same parameters as amplicon sequences.

## 3 Results

### 3.1 Evaluating the need for degeneracy in broad-range *hgcAB* primers

To evaluate whether the degeneracy of the new primer set was high enough to capture the natural diversity of *hgcAB*, we queried *hgcAB* libraries constructed from several environmental matrices (Table S3). The *hgcAB* products were searched for primer-binding locations to resolve which primer sequences were likely binding and amplifying *hgcAB* genes from environmental samples. Libraries were constructed using both the original (ORNL-HgcAB-uni-F, ORNL-HgcAB-uni-R) primer set and the new less degenerate primer set (ORNL-HgcAB-uni-F, ORNL-HgcAB-uni-32R) for comparison. We evaluated our clone libraries constructed from a range of environments including boreal peats, freshwater stream sediments, freshwater periphyton mats, and salt marsh and rice paddy soils (Table S3). We also evaluated high-throughput *hgcAB* amplicon libraries for the New Horizon site in EFPC, chosen due to the expected high diversity in Hg-methylators at this site (Table S5).

Forward primer binding sites were found in 84% of the clones and 95% of the amplicons searched. Each of the 48 possible primer oligonucleotide sequences were found in the clone and amplicon *hgcAB* sequence libraries (Figure S2). The relatively even distribution of percent occurrence among the forward oligonucleotide sequences in environmental sequences indicates that the degeneracy (n =48) of the forward primer is necessary for capturing *hgcAB* diversity. The *hgcAB* clone and amplicon libraries sequenced were searched for reverse oligonucleotide sequences from both ORNL-HgcAB-uni-R and ORNL-HgcAB-uni-32R primers (Figure2). In amplicon sequence libraries amplified with ORNL-HgcAB-uni-F and ORNL-HgcAB-uni-R, 39.2% of oligonucleotide sequences found in amplicon sequences were shared between primer sets (Figure 2). In amplicon libraries amplified with ORNL-HgcAB-uni-F and ORNL-HgcAB-uni-32R, the percentage of shared oligonucleotide sequences found in amplicon sequences increased to 89.9% (Figure 2). When allowing up to two mismatches, both reverse primers theoretically bind to all available clone and amplicon sequences with reverse primer binding sites from this study. This indicates that theoretical binding efficiencies of both reverse primers to *hgcAB* genes in environments tested are highly similar. This is supported by the high similarity between taxonomic profiles of *hgcAB* genes amplified from the same sample but with different reverse primers (Figure 3).

**Figure 2.**
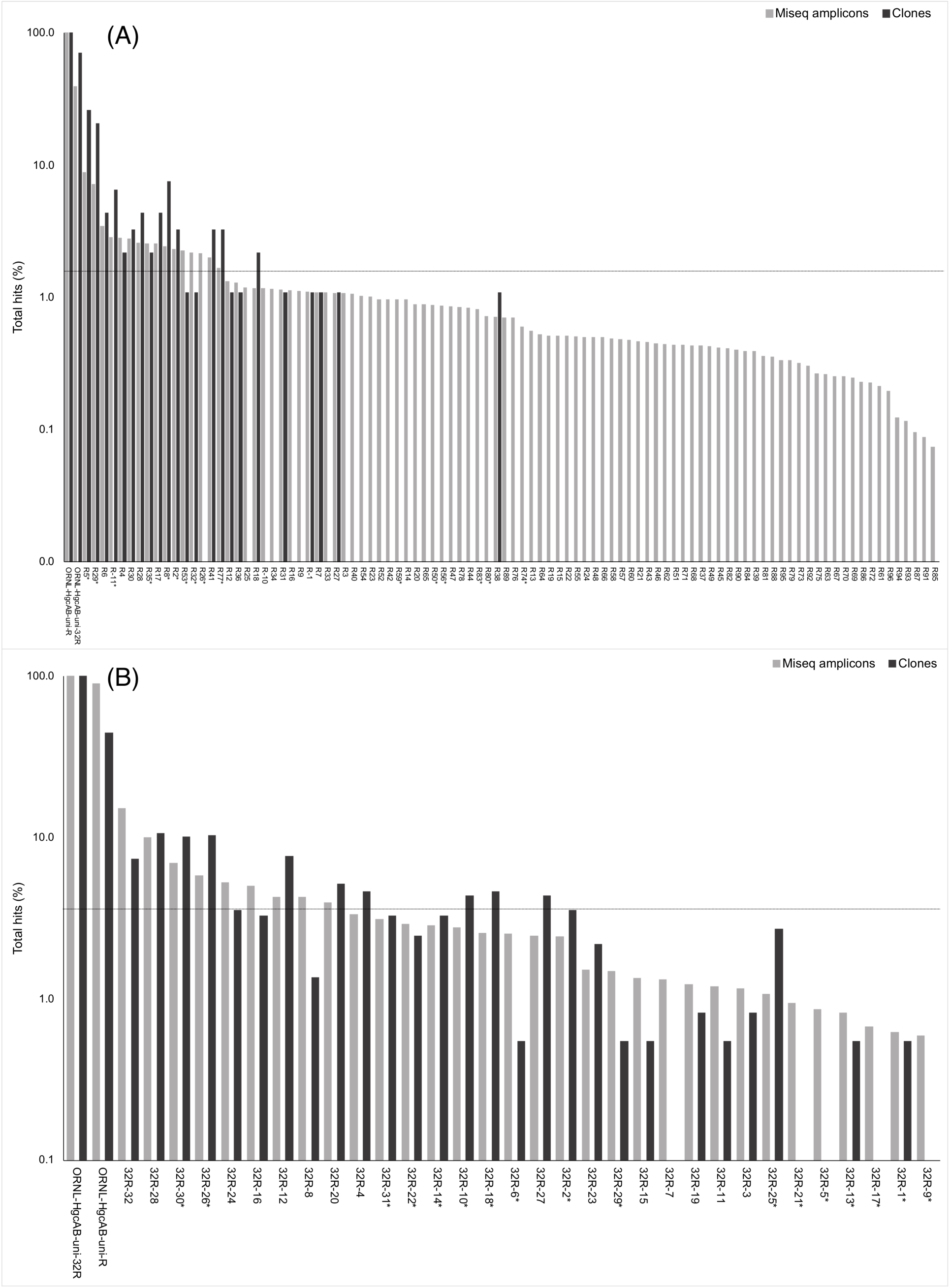
Percent occurrence of each 96-degenerate reverse (top; ORNL-HgcAB-uni-R) and 32-degenerate reverse primer (bottom; ORNL-HgcAB-uni-32R) oligonucleotide sequence in the environmental clone *hgcAB* sequences and Miseq amplicons. Primers are listed in Table S2. Matches to each broad-range reverse primer (ORNL-HgcAB-uni-R, ORNL-HgcAB-uni-32R) are shown for each set of clone and amplicon sequences amplified with the ORNL-HgcAB-uni-R (A) and ORNL-HgcAB-uni-32R (B). Oligonucleotide sequences shared between each reverse primer indicated with asterisk (*). Reverse primer sequences were found in 95 clones (A) and 369 clones (B), and 210, 362 (A) and 233, 934 (B) amplicon reads. Data shown on a log scale. Dashed line indicates equal distribution of reverse primer sequences across amplicon sequences: A) 1.04%, B) 3.13%. Ordered by decreasing abundance in Miseq amplicon libraries.

**Figure 3.**
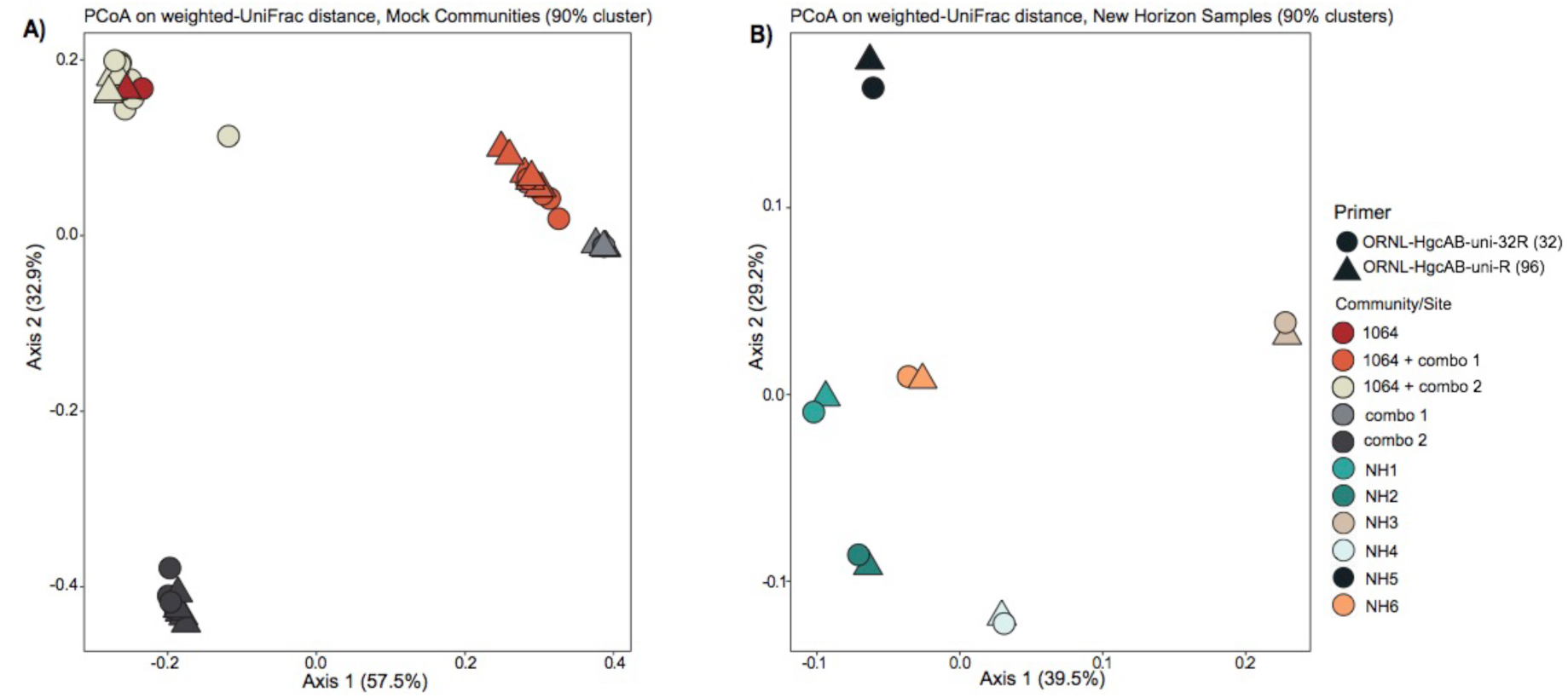
Principle coordinate analyses (PCoA) showing the variance in phylogenetic distances between mock community (A) and New Horizon (NH) sediment (B) *hgcAB* amplicon libraries. Reverse primer (i.e. ORNL-HgcAB-uni-R, ORNL-HgcAB-uni-32R) is indicated by shape. Weighted UniFrac distances calculated from the taxonomic classifications of clustered (90%) OTUs using the program ‘pplacer’ and the R package ‘phyloseq’.

One of the goals for reducing the degeneracy of the reverse primer from 96 to 32 was the removal of redundant oligonucleotide sequences, i.e. sequences that do not actually bind and amplify template DNA. Removal of the redundant sequences from the primer mix should increase amplification efficiencies by increasing the ratio of each of the remaining oligonucleotide sequences to template DNA (Lee et al., 2012). Decreasing degeneracy also reduced the annealing temperature range of the primer-set (Table S2), which can also have an effect on PCR amplification bias (Lee et al., 2012). There are 50 sequences from ORNL-HgcAB-uni-R that do not appear in the reference *hgcB* database (i.e. 52.1% redundant) (Table S2). When searching the 210,362 amplicon sequences amplified by ORNL-HgcAB-uni-R, each of the 96 ORNL-HgcAB-uni-R oligonucleotide sequences are found in amplicons. However, 65 oligonucleotide sequences fall below the 1.04% threshold that represents equal distribution of reverse primer sequences across amplicon sequences; 59 of which were not conserved in the primer redesign (Figure 2). Of the 32 possible ORNL-HgcAB-uni-32R sequences, 14 do not appear in the reference *hgcB* database (i.e. 43.8% redundant) (Table S5). The ORNL-HgcAB-uni-32R oligonucleotide sequences were tallied for each clone (n = 369) and amplicon (n = 233,934) amplified by ORNL-HgcAB-uni-32R to determine the frequency of each oligonucleotide to generate a PCR product (Table S5). When searching the 233,934 *hgcAB* amplicon sequences, all 32 ORNL-HgcAB-uni-32R oligonucleotide sequences match at least 0.59% of amplicons. However, 22 oligonucleotide sequences fall below the 3.13% threshold that represents equal distribution of reverse primer sequences across amplicon sequences (Figure 2). This indicates that primer degeneracy could further be reduced if desired.

We identified six sequences that were underrepresented in both clone and amplicon libraries and rarely found in reference *hgcAB* sequences, ORNL-HgcAB-uni-32R-05, ORNL-HgcAB-uni-32R-09, ORNL-HgcAB-uni-32R-17, ORNL-HgcAB-uni-32R-21, ORNL-HgcAB-uni-32R-15, and ORNL-HgcAB-uni-32R-7 (Table S5). These six oligonucleotide sequences were removed during the preparation of equimolar mixtures of the ORNL-HgcAB-uni-32R primer and labeled ‘equimolar 26’. Equimolar mixtures of the 32 and 26 reverse oligonucleotide sequences were used to amplify *hgcAB* genes from peatland soil, periphyton, and freshwater sediments (Figure S3). Decreasing degeneracy in the reverse primer sequence from 96 to 32, and further to 26, appeared to produce brighter, clearer bands at the target amplicon size (∼950 bp) (Figure S3).

### 3.2 In-silico evaluation of ORNL-HgcAB-uni-32R primer binding efficiency and specificity

We compared the theoretical binding efficiency of ORNL-HgcAB-uni-32R and ORNL-HgcAB-uni-R primers to reference *hgcB* sequences (Table S6). Since mismatches likely occur between primers and template DNA, *in-silico* binding efficiencies were calculated allowing for up to two mismatches. Mismatches within five bases of the 3’ were not allowed. When allowing for mismatches, binding efficiencies of both reverse primers to reference sequences increased to 98% coverage (Table S6; Figure S4). Mismatches primarily occurred at the third or fourth nucleotide base from the 5’ binding position of the reverse primer. This correlates with the fairly conserved 5’ end guanines which are (>30% but ≤100% sequence identity) (Figure 1). Reverse primer degeneracy was decreased by a factor of three by changing the third base of ORNL-HgcAB-uni-R from ‘B’, (C/T/G), to ‘G’. Mismatches at these sites would likely not decrease binding efficiencies due to the proximity to the 5’ end of the primer-bind region (Campos and Quesada, 2017). When analyzing the ability of each primer to bind *in-silico* to *hgcB* sequences from various clades, sequences that appear most often in the database were conserved between the primers (Figure S4). When allowing for mismatches, there was no loss in primer binding efficiency to *hgcB* from various clades, despite decreased degeneracy (Figure S4, Table S6).

The reverse primer targets the highly conserved ‘C(M/I)ECGA’ motif in HgcB. This triple cysteine (CXXCXXC) motif is well-conserved in other ferredoxins so non-specific reverse primer binding may occur. *In-silico* primer specificity was tested against a reference database of 14,161 ferredoxin-encoding gene sequences. Combined, the reverse primers aligned to 786 sequences (5.6% of the database), with up to two mismatches. Of those matches, only 88 sequences were predicted to encode HgcB from possible Hg-methylators. Sequences were identified as HgcB by conserved motifs, sequence similarity to known HgcB using blastp, and presence of an HgcA-encoding region upstream of HgcB (if scaffolded sequence was available). Both reverse primers aligned to the 88 *hgcB* sequences. The additional 698 genes likely encode ferredoxins paralogous to HgcB. Shared sequences of both primers did not align completely with any paralogous sequence but matched 114 sequences when allowing for two mismatches (<1% of total ferredoxin sequences). When allowing mismatches, both reverse primers performed quite similarly *in-silico*. Sequences unique to the less degenerate primer, ORNL-HgcAB-uni-32R matched 424 paralogous ferredoxins (3.0% of total ferredoxin sequences), while sequences unique to ORNL-HgcAB-uni-R matched 344 paralogous sequences (2.4% of total ferredoxin sequences), when allowing up to two mismatches. Overall, the primer sequences appeared unique to *hgcB*, and false positives occurred only when mismatches were allowed. Decreasing the degeneracy of the reverse primer did not decrease primer binding to genes encoding non-HgcB ferredoxins *in-silico*. However, less degeneracy may increase selective amplification of *hgcA* and *hgcB* from environmental samples by limiting false positives from hybridization.

### 3.3 Amplification of *hgcAB* from pure cultures using ORNL-HgcAB-uni-R32

The new less degenerate reverse primer, in combination with ORNL-HgcAB-uni-F, correctly amplified ∼950 bp *hgcAB* product from 25 *hgcAB*+ pure cultures. Species tested included *Deltaproteobacteria, Firmicutes* and *Archaea* (Christensen et al., 2016). The 20 µl PCR mixture contained 1×10^6^ copies of gDNA from pure cultures and 1.0 µM of each primer. As with ORNL-HgcAB-uni-R, no amplification was observed from *Pyrococcus furiosus* Vc1 which has a fused *hgcAB* gene. No product of the correct size was amplified from five *hgcAB*^−^ cultures with either reverse primer. However, multiple bands of smaller size, and thus likely not *hgcAB*, were amplified from most pure cultures. Repeated attempts to purify, ligate, and clone PCR products <950 bp failed. These extra bands are likely mispriming, possibly to ferredoxin-encoding genes, or primer dimers. Overall, amplification of *hgcAB* from known methylators using the new reverse primer was consistent with amplification from the more degenerate reverse primer (Christensen et al., 2016).

### 3.4 Optimizing PCR amplification of *hgcAB* from environmental samples

In addition to improving the reverse primer, other PCR-modifications were tested to improve *hgcAB* amplification efficiencies from environmental samples. Using DNA from sediment, soil and periphyton samples, and *Desulfovibrio desulfuricans* ND132, we tested template concentrations between 10 ng to 100 pg, reaction volumes from 20 μl to 50 μl, and primer concentrations from 1.0 μM to 0.5 μM (see Table S7). For low template concentrations, increasing the reaction volume to 50 μl was necessary in order to ensure enough PCR product could be gel-excised and cleaned prior to high-throughput sequencing. Both the 96-fold degenerate ORNL-HgcAB-uni-R and 32-fold degenerate ORNL-HgcAB-uni-32R were tested. PCR products were compared visually on 1% agarose gels (Figure S3), for brightness of the ∼950 bp band. An important result was that no single PCR protocol was ideal for all samples. Periphyton were particularly difficult to amplify, although we observed improved efficiencies with lower primer to template ratios (Condition B). This was somewhat unexpected, since increased primer to template ratios can reduce PCR inhibitors (Schrader et al., 2012). In contrast, a clear PCR product was produced from freshwater sediment and peatland soil under multiple PCR conditions. We therefore recommend optimizing protocols for each sample matrix, and not assuming a one-size fits all strategy. Optimal template concentration can be determined by 16S rRNA amplification (8F/1492R) (Turner et al., 1999).

### 3.5 Recovery of Mock Communities

The revised reverse primer ORNL-HgcAB-uni-32R recovered *hgcAB*^+^ cultures spiked into salt marsh soils in about the same abundance as the more degenerate ORNL-HgcAB-uni-R (Figure 4). Recovery of mock communities was performed using high-throughput amplicon specific sequencing. Both primer sets recovered *Mm. luminyensis, Mc. bavaricum, Dv. desulfuricans* ND132, *G. sulfurreducens* and *Df. dehalogenans* mock community “Combo 1” was spiked with a mix of cells containing 41% *G. sulfurreducens*, 50% *Ds. acidophilus*, and 9% *Mc. bavaricum. Ds. acidophilus* was substantially under-recovered by both primer sets, although pure *Ds. acidophilus* gDNA was amplified by both. One explanation may be the difficulty in isolating high quality gDNA from endospore-forming bacteria such as *Ds. acidophilus* (Filippidou et al., 2015). The taxonomic composition of mock community “Combo 2” was 59% *Dv. desulfuricans* ND132, 7% *Df. dehalogenans*, and 35% *Mm. luminyensis. Mm. luminyensis* was over-represented relative to the spike, *Dv. desulfuricans* ND132 was somewhat under recovered, while *Df. dehalogenans* recoveries were similar to the spike. Interestingly, *Df. dehalogenans hgcAB* was better recovered using the less degenerate primer. This may be due to higher *Deltaproteobacteria* and *Methanomicrobia* representation in the 96-degenerate primer-mix, as seen in the *in-silico* binding efficiencies (Table S6). The 32-degenerate primer also had a higher theoretical binding capacity to *Deltaproteobacteria* and *Methanomicrobia* compared to other clades, however the differences were not as significant. Discrepancies between expected and actual DNA recovery could be due to extraction efficiencies.

**Figure 4.**
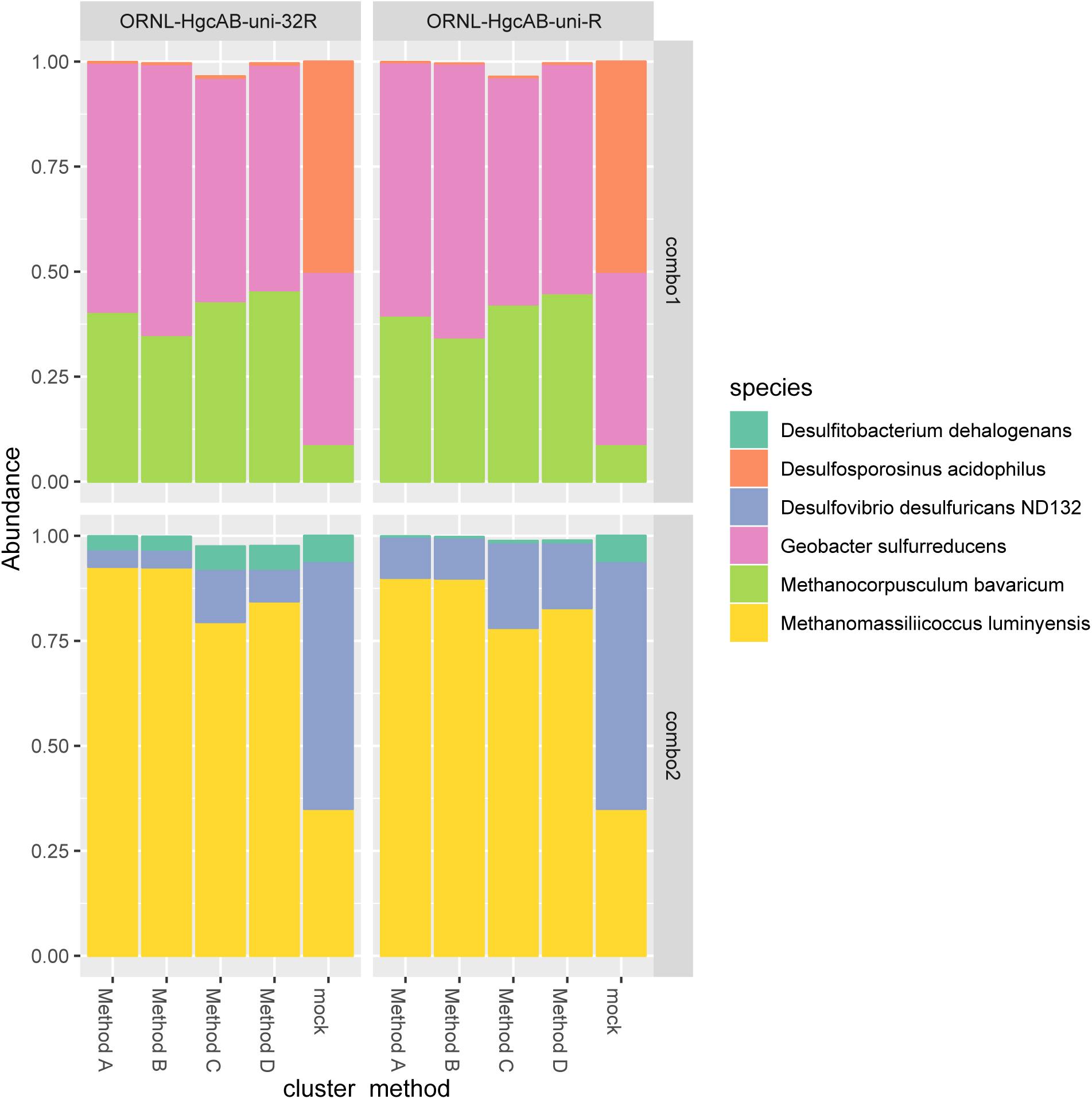
Comparison of expected and actual taxonomic composition of *hgcA* sequences in recovered mock community amplicon libraries. Top, mock community Combo1; bottom, mock community Combo2. Libraries were constructed using the newer less degenerate *hgcAB* reverse primer (left) or the Christensen 2016 reverse primer (right). Comparison of various clustering methods: **Method A**: cluster nucleotide sequences at 90% identity using VSEARCH; **Method B:** cluster peptide sequences at 90% identity, using USEARCH; **Method C:** clustering nucleotide sequences at 90% using cd-hit-est; **Method D:** chimeras removed prior to clustering nucleotide sequences at 90% using cd-hit-est.

### 3.6 Comparing OTU clustering methods of *hgcA* and HgcA sequences

We investigated variance in nucleotide and translated peptide sequences of reference *hgcA* genes to determine best sequence identity cut-offs for clustering and classifying environmental clone and amplicon sequences. Clustering *hgcA* nucleotide sequences and classifying the translated peptide sequences at a sequence identity cut-off of 90% retained genus-level resolution in taxonomic classifications. Sequence divergence of reference *hgcA* genes appear incongruent to phylogeny. In the 201 nt bp reference *hgcA* database, 79.0% of the 283 *hgcA* sequences are unique from one another. Variance in *hgcA* nucleotide sequences increased over the 654 nt bp region, with the percent of unique sequences increasing to 80.7%. Some *hgcA* reference sequences were highly variant (<80% sequence identity) to one another even amongst genus-level representatives (e.g. *Syntrophus, Desulfovibrio, Desulfobulbus, Desulfitobacterium*). Reference *hgcA* genes from strain-level representatives or from MAGs from the same environment often had 100% sequence identity to one another over these two regions. With the exception of two *hgcA* genes from bacterial endosymbionts (i.e. *Spirochaeta* sp. JC202 and *Deltaproteobacterium* sp. OalgD1a), all other reference *hgcA* with 100% sequence identity shared common phylogeny. Over the 201 nt bp region of *hgcA*, 73.0% of reference sequences were unique at a 90% sequence identity cut-off, and 65.5% unique with an 80% sequence identity cut-off. Over the 654 nt bp region, 73.0% of reference *hgcA* sequences were unique at 90% sequence identity cut-off, and 68.9% were unique at an 80% cut-off. We found that clustering nucleotide sequences at 90% identity over both the 201 nt bp and 654 nt bp region kept genus-level resolution of *hgcA* phylogeny. With an 80% cut-off, clustering of *hgcA* from reference genomes of different phyla but the same environment occurred (e.g. *Nitrospira* bacterium SG-35-1 and *Desulfobacterales* bacterium SG-35-2).

Clustering of peptide sequences at 80-90% sequence identity also affected the reliability in classifying translated *hgcA* sequences. Collapsing subtrees of the translated 201 nt bp (67 aa bp) HgcA reference tree at a 90% cut-off reduced the nodes from 283 to 181, combining species-level variants as well as some MAGs and single cell genomes from the same environment and phyla. Collapsing the HgcA reference tree at an 80% cut-off reduced the nodes from 283 to 136 and resulted in a loss of resolution to the order level or higher for some subtrees. For example, the HgcAB fused proteins related to *Desulfarculaceae, Kosmotoga*, and *Methanococcoides* are 80% similar across the translated 201 nt bp region. The subtree containing HgcA from *Aminicenantes* and *Atribacteria* marine genomes also collapsed with an 80% cut-off. Other subtrees that collapsed included HgcA from *Dehalococcoides mccartyi* with *Clostridium cellulosi* and HgcA from *Dehalobacter* species with HgcA from *Bacteroides cellulosolvens, Clostridium cellobioparum* and *Acetivibrio cellulolyticus*, and *Acetonema longum*. Therefore, we recommend both clustering and classifying environmental samples with a 90% nt and aa sequence identity cut-off. High sequence similarity (>90%) between environmental and reference sequences likely indicates they are close relatives at the species level.

We compared several methods for OTU clustering using the mock community amplicon datasets (Figure 4). We clustered non-chimeric dereplicated nucleotide sequences at 90% with VSEARCH, clustered nucleotide sequences not filtered for chimeras at 90% using cd-hit-est, and clustered translated peptide sequences at 90% similarity with USEARCH (Rognes et al., 2016) (Figure 4). OTU clustering method affected relative abundance distribution of mock community *hgcA* amplicon sequences (Figure 4). Clustering of nucleotide sequences at 90% identity using cd-hit-est was closest to the expected composition of the mock community. However, biases are likely to occur with any clustering method. Chimera filtering removed 7.9% of sequences from the mock community *hgcAB* dataset (Figure 4). Non-chimeric sequences clustered at 90% using cd-hit-est were used for downstream analyses.

### 3.7 Direct amplicon sequencing from environmental samples

We validated PCR amplification of *hgcAB* from environmental matrices by Sanger sequencing the cloned PCR product and directly sequencing PCR amplicons. We observed similar phylogeny in *hgcAB* clone and amplicon sequences amplified using both primer-sets. The *hgcAB* genes were amplified from pooled SPRUCE peatland gDNA (SPRUCE1, clone IDs: OP1, NP1; Table S3), freshwater periphyton and sediment from NH (NH3, clone IDs: OP3, NP3; Table S3) using PCR condition B (Table S7; Figure S3). Amplification from periphyton was poor, and only the PCR products from SPRUCE soil and NH sediment that amplified with both primer sets were chosen for cloning (Figure S3). The trimmed and quality filtered *hgcAB* clone libraries for SPRUCE (n=58, n=65) and NH (n=42, n=61) using the 96- and 32-degenerate primers, respectively, showed similar diversity (Figure 5). The NH *hgcAB* genes (NH3) amplified from both primer sets were also directly sequenced, resulting in 65,832–106,964 reads per sample (Table S4). A direct comparison of the two sequencing strategies revealed a highly similar phylogenetic distribution (Figure 5).

**Figure 5.**
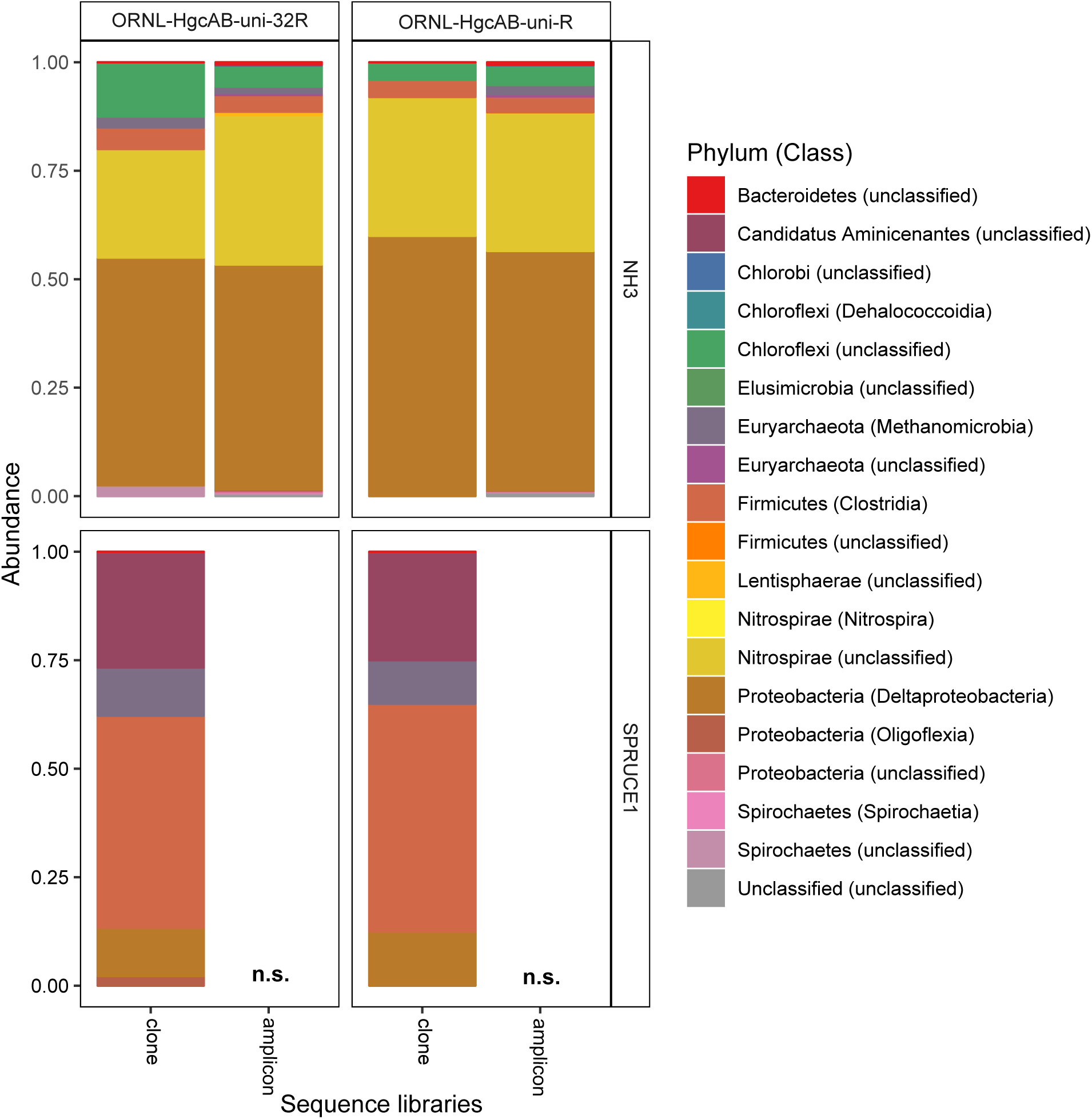
Relative abundance of cloned and amplicon *hgcAB* sequences by phyla and class. *hgcAB* genes were amplified from New Horizon sediment (NH3) and SPRUCE soil (SPRUCE1) using ORNL-HgcAB-uni-F with both reverse primers (ORNL-HgcAB-uni-R, ORNL-HgcAB-uni-32R). Taxonomic classifications based on phylogenetic placement of representative sequences on the reference tree (‘ORNL_HgcA_201.ref.pkg’) using program Taxtastic with classification cutoff of 90%. OTUs have been merged at the class-level, with relative abundance values calculated from proportion of total clone of amplicon sequences in each library. Figure produced using R packages ‘phyloseq’ and ‘ggplot2’. *hgcA* amplicons were not sampled (n.s.) from SPRUCE soils.

Both the mock and NH communities were used to compare performance of both primer-sets in capturing diverse *hgcAB* sequences (Figure 3). Principle coordinate analyses (PCoA) of phylogenetic distances between *hgcA* OTUs indicated that sample-type rather than primer-set accounted for the greatest variance in *hgcA* diversity (Figure 3). Furthermore, taxonomic composition of *hgcA* OTUs amplified from the same mock community or NH sample, but using different reverse primers, were highly similar (Figure 6). This indicated that decreased reverse primer degeneracy did not lower the captured *hgcAB* diversity from the mock communities or environmental samples.

**Figure 6.**
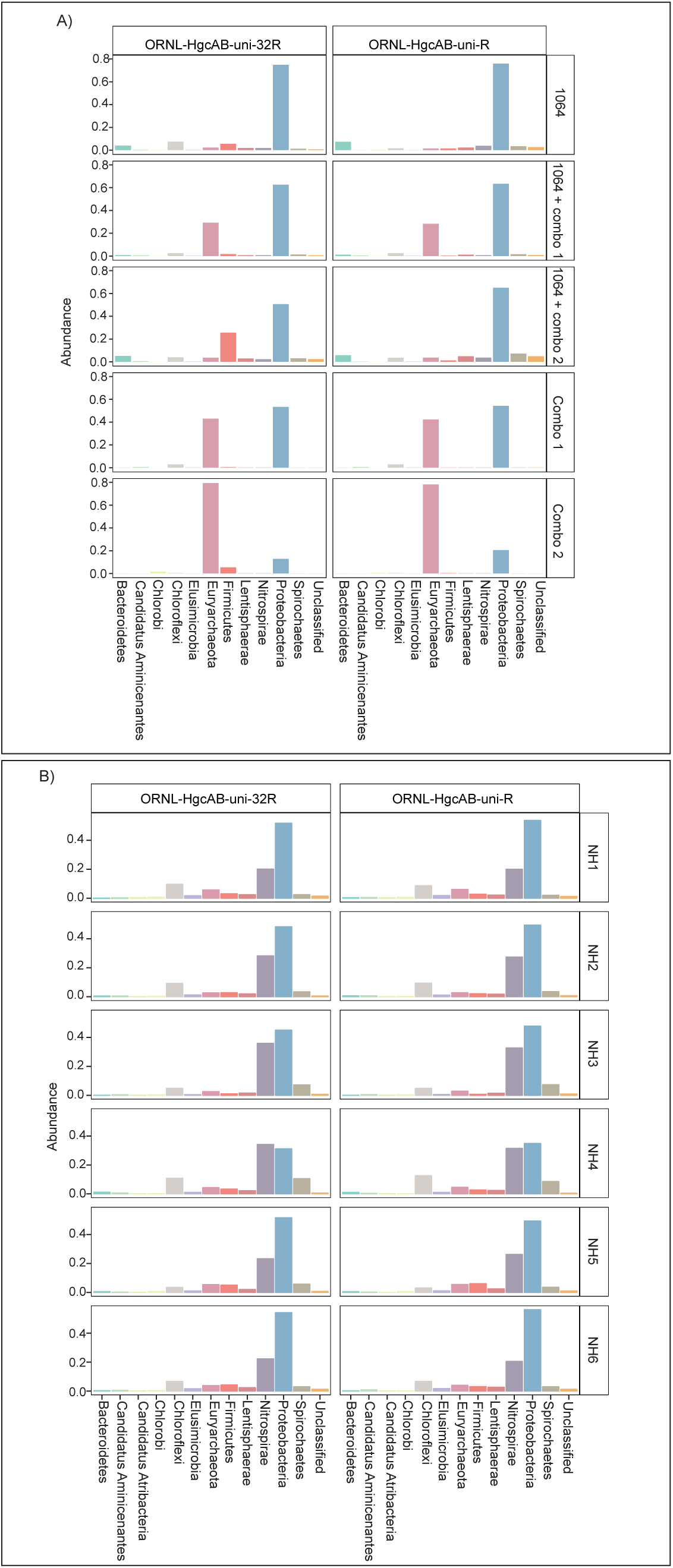
Relative abundance profiles of *hgcA* amplicons from mock (A) and environmental (B) NH sediment samples at the phyla level. OTUs were clustered at 90% sequence similarity, and taxonomic classifications were assigned based on phylogenetic placement of representative sequences on the ‘ORNL_HgcA_201.ref.pkg’ reference tree using the program ‘pplacer’ with a classification cut-off of 90%. OTUs have been merged at the phyla level.

All *hgcAB* sequences from NH3 and SPRUCE1 clone and amplicon libraries were compared by trimming to 201 nt bp and clustered on a 90% sequence identity cut-off, resulting in 3,861 unique operational taxonomic units (OTUs). Taxonomic classifications of translated *hgcA* sequences were similar between two clone libraries produced from the same sample but different primer-sets (Figure 5). The *hgcA* clones from SPRUCE (SPRUCE1; OP1, NP1) were classified as *Methanomicrobia, Deltaproteobacteria, Firmicutes* and *Aminicenantes* (Figure 5). The *hgcA* clones from NH (NH3; NP3, OP3) primarily grouped with *Chloroflexi, Firmicutes, Methanomicrobia, Nitrospirae*, and *Deltaproteobacteria*, particularly *Geobacter hgcA* sequences (Figure 5). However, at the class level, several OTUs were unique to clone libraries produced with the new primer including unclassified *Spirochaetes* and *Lentisphaerae, Oligoflexia*, and *Methanomicrobia* (Figure 5). While rarefaction curves of clone libraries did not approach saturation, deeper MiSeq amplicon sequencing of the NH3 *hgcA* captured these excluded OTUs from each primer set. This indicated that both primer-sets, when evaluated at sufficient sequencing depth, can capture the same *hgcAB* diversity.

### 3.8 Broad-range detection of *hgcAB*+ microorganisms in environmental samples

The new primer set captured a diverse array of sequences similar to *hgcA* from cultured isolates as well as environmental MAGs. A phylogenetic analysis of translated *hgcA* sequences highlights several novel potential Hg-methylators (Figure 7). The *hgcA* high-throughput amplicon libraries from freshwater EFPC sediment and tidal marsh sediment contained sequences closely related to known and potential Hg-methylators from the three major clades, *Deltaproteobacteria, Methanomicrobia*, and *Firmicutes* (Figure 7) as well as novel sequences similar to *Nitrospirae, Elusimicrobia, Spirochaetes, Chloroflexi, Lentisphaerae, Bacteroidetes, Atribacteria*, and candidate phyla WOR-3 and KSB1 bacteria (Figure 7) with many lacking cultured representatives. In SPRUCE peatland clone libraries, both primers captured *hgcA* sequences closely related to *Aminicenantes* (Figure 5); however, this clade was not prevalent in freshwater or tidal marsh amplicon libraries (Figure 7). Interestingly, *hgcA* amplicons and clones closely related to *Bacteroidetes* and *Chloroflexi* were prevalent in both tidal marsh and freshwater sediments, while nitrogen-cycling *Nitrospirae* were prevalent primarily in freshwater sediment (Figures 5,6). Several *Chloroflexi, Bacteroidetes, and Nitrospirae* containing *hgcA* have been sequenced (Baker et al., 2015; Podar et al., 2015a; Anantharaman et al., 2016; Jones et al., 2019) but none have been tested for Hg-methylation. *Nitrospirae* are often found alongside sulfate-reducing bacteria, and can perform sulfate-reduction (Henry et al., 1994; Teske et al., 1994; Zecchin et al., 2018). Further, *Nitrospirae* and *Bacteroidetes* possess the reductive acetyl-CoA (Wood-Ljungdahl) pathway for carbon fixation and acetate production, which may be a connected biochemical pathway for HgcAB function (Baker et al., 2015). This expanded phylogeny of potential Hg-methylators includes metabolic clades associated with nitrogen-cycling, fermentation, and chemoheterotrophy. Members of these metabolic clades have previously been connected to Hg-cycling in environmental metagenome studies (Podar et al., 2015a; Gionfriddo et al., 2016; Jones et al., 2019).

**Figure 7.**
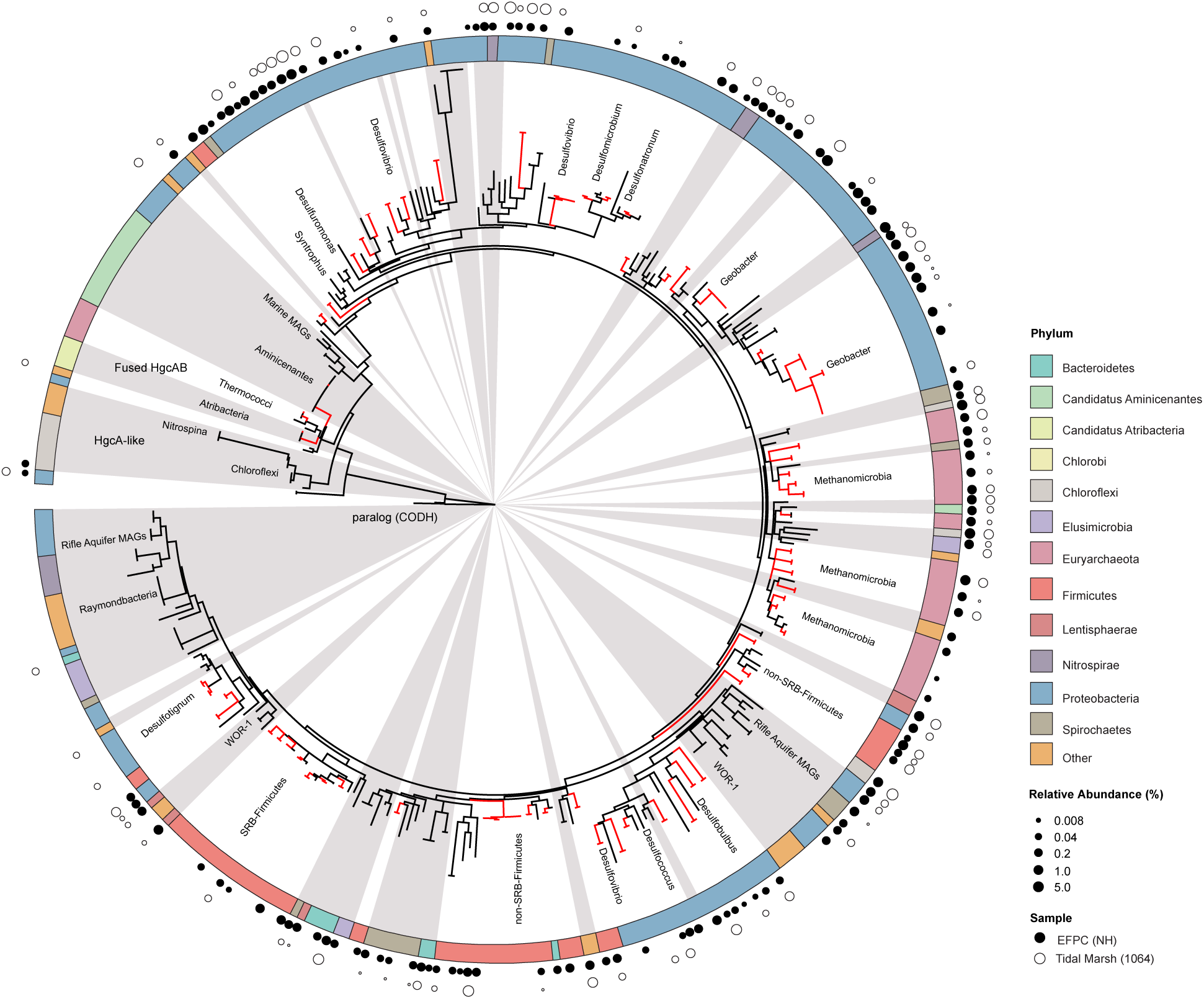
Molecular phylogenetic analysis of HgcA reference sequences by Maximum Likelihood method. Phylogeny was calculated using the GAMMA model of rate heterogeneity and ML estimate of alpha-parameter in RAxML with 150 bootstrap replicates. The tree was rooted by carbon monoxide dehydrogenases from non-Hg-methylators *Candidatus Omnitrophica bacterium* CG1-02-41-171 and *Thermosulfurimonas dismutans*. Relative abundance of *hgcA* amplicons from Tidal Marsh sediment (1064) and East Fork Poplar Creek NH sediments were calculated from OTU clustering at 90%, and the values placed on tree using ‘pplacer’. Phylum-level classifications of HgcA reference sequences are color-coded, and distinct subtrees are labeled, including metagenome-assembled genomes from Rifle Aquifer Field Research Site and marine datasets. ‘Other’ phyla include phyla with low number of HgcA representatives: *Planctomycetes, Nitrospina*, and Candidate phyla *Raymondbacteria*, WOR-1, WOR-3, and *Wallibacteria*. Gray background indicates reference sequences are from metagenome-assembled genomes (MAGs) or environmental single cell genome center sequencing (SCGC) datasets. Red branches indicate that sequences were included in phylogenetic analysis from Podar et al.(Podar et al., 2015a). Black branches are sequences that have since been identified.

Directly sequencing *hgcA* amplicons enables a deeper dive into the diversity of the Hg-methylator community compared to cloning and metagenomic techniques, especially when community composition is complex. We used rarefraction curves to compare the richness of *hgcA* OTUs in NH sediment captured by cloning, high-throughput amplicon, and metagenomic sequencing techniques (Figure S5). The NH3 clone and amplicon *hgcA* sequences from this study were compared to *hgcA* sequences pulled from a NH sediment metagenome from a previous study (Christensen et al., 2019). The *hgcA* sequences from each technique were pooled into one dataset and processed following the OTU-clustering and *hgcA* classification pipeline described above. The clone and metagenome datasets captured only a small subset of the total number of *hgcA* OTUs compared to high-throughput amplicon sequencing. The NH3-uni-R clone library captured 25 OTUs from 33 reads, the NH3-uni-32R captured 42 OTUs from 57 reads, and the NH metagenome captured 63 OTUs from 79 *hgcA* reads (pulled from 6.31 Gbp of metagenomic sequencing). The clone libraries captured the relatively abundant OTUs in the HTS dataset. The NH3-uni-R and NH3-uni-32R clone OTUs contained 25.6% and 33.2% of the total abundance of *hgcA* reads in the dataset, respectively. The metagenome *hgcA* OTUs represented only 1.5% of the total number of reads in the dataset. However, this may be because the metagenomic sequencing was performed on a different sample than NH3 and is not directly comparable. Overall it appears that deeper sequencing would be needed to capture the same diversity of *hgcA* sequences captured by high-throughput amplicon sequencing (Figure S5).

The retention of low abundant and unique sequences may over-estimate *hgcA* diversity in a sample. When singleton sequences were kept in analysis, the amplicon dataset produced using the more degenerate primer-set (NH3-amplicon-uni-R) had a higher richness in OTUs (Figure S5). Inclusion of singleton sequences in rarefraction analysis likely overestimates OTU richness because of spurious OTUs resulting from sequencing error (Huse et al., 2010). Indeed, the sequencing error rate calculated using the mock community amplicons produced with ORNL-HgcAB-uni-R were slightly higher (2.20–2.27%) compared to ORNL-HgcAB-uni-32R (1.94– 2.05%). We recommend removing singletons prior to OTU-clustering and performing quality filtering steps such as trimming poor-quality bases, chimera filtering, and filtering non-*hgcA* reads using the hmm-model to remove spurious OTUs. Even following these guidelines, it is important to note that high-throughput amplicon sequencing may still over-estimate *hgcA* abundance in a sample.

### 3.9 Insights from applying the updated protocol to a previously-studied environment

Use of the new primer set, *hgcAB* reference library and improved PCR protocols expanded the known diversity of methylators in EFPC sediments, a previously well-studied site (Mosher et al., 2012; Christensen et al., 2016; Christensen et al., 2019). Prior studies at EFPC identified the dominant members of the Hg-methylating community including *Deltaproteobacteria*, methanogenic Archaea, and *Firmicutes* based on 16S rDNA sequencing, clone-derived *hgcAB* sequencing, and meta-omic analyses (Mosher et al., 2012; Christensen et al., 2016; Christensen et al., 2019). With the improved primer set and analysis pipeline we have recovered *hgcA* genes from all clades represented in the previous studies as well as novel potential methylators like *Nitrospirae* and *Elusimicrobia*, as discussed above. The high-throughput nature of this updated method allowed for comparisons of the *hgcA* population across the sediment depth profile (Figure 8) showing the relative abundance of *Proteobacteria* decreasing with depth while *Firmicutes* and methanogenic Archaea increase in relative abundance. *Nitrospirae* containing *hgcA* were abundant in these samples, representing up to 20% of the *hgcA* population. The updated analysis pipeline provides a more nuanced understanding of the Hg-methylating community than previously available and suggests that novel potential methylators may represent a larger proportion of the Hg-methylating community in the stream than previously realized.

**Figure 8.**
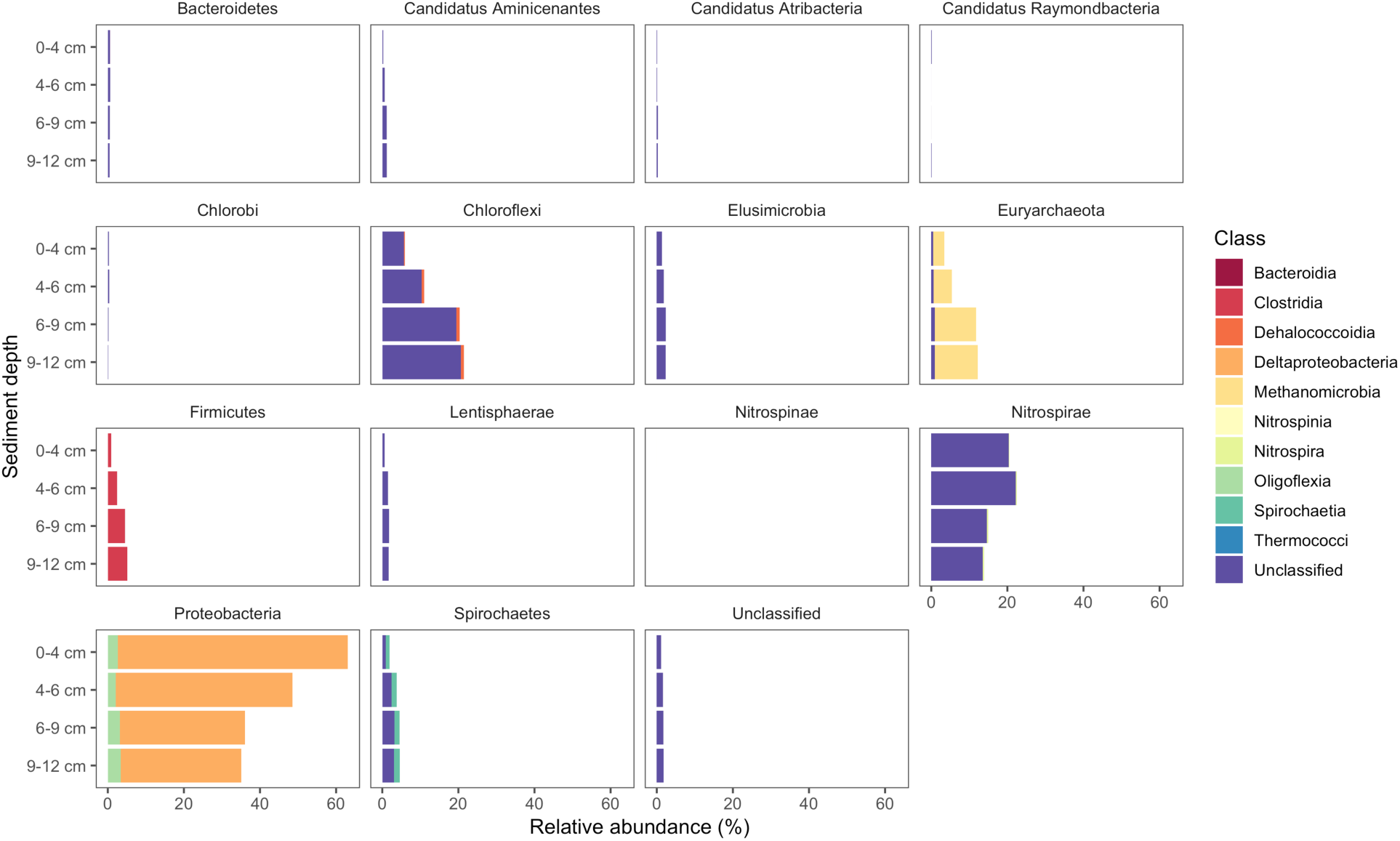
Known and novel methylators in a sediment core from East Fork Poplar Creek (Oak Ridge, TN). Relative abundance of *hgcA* phyla across a 12cm sediment core were identified using the new degenerate primers, direct sequencing technique, and the updated *hgcA* database. Known methylators (*Deltaproteobacteria, Firmicutes-Clostridia*, and *Euryarchaeota*-*Methanomicrobia*) were recovered as well as novel potential methylators.

## 4 Discussion

Broad-range PCR primers can measure the presence and diversity of genes in environmental samples. In this study, we provide updated primers and protocols for amplifying genes encoding Hg-methylation to best capture Hg-methylator diversity. While primer degeneracy is necessary to capture *hgcAB* diversity, a trade-off in specificity and binding efficiencies may occur. This new and improved PCR protocol was modified to enhance amplification efficiencies of *hgcAB* from environmental samples by decreasing the reverse primer degeneracy from 96 to 32. The reverse primer is paired with the published forward broad-range primer which has a degeneracy value of 48 (Christensen et al., 2016). Since the inception of *hgcA* and *hgcAB* primers (Bae et al., 2014; Liu et al., 2014; Schaefer et al., 2014; Christensen et al., 2016), most amplification has been followed by laborious clone library construction and Sanger sequencing. One drawback is the limited number of *hgcAB*^+^ organisms that can be identified (50–100 clones/library) as well as inherent protocol bias. Direct amplicon sequencing is desirable given the increased data-output and lower labor (Bravo et al., 2018a; Liu et al., 2018b). Here we include direct *hgcAB* amplicon sequencing which was validated using a mock community and tidal salt marsh soils spiked with the mock community.

### 4.1 Evaluating HgcA phylogeny

With the growing number of available reference HgcA-encoding sequences, it is apparent that the phylogeny of HgcA sequences is not congruent to species trees. While the majority of HgcA sequences tend to cluster by clade, there are several exceptions (Figure 7). The *Deltaproteobacteria* HgcA are most similar to one another, with sulfate-reducing, syntrophic, and metal-reducing members forming their own subtrees. The same is true for fermentative and sulfate-reducing *Firmicutes*, and *Methanomicrobia*. However, clustering patterns are not as well defined for HgcA representatives from lesser represented taxa, for example *Elusimicrobia, Spirochaetes, Phycisphaerae* and *Lentisphaerae* (Figure 7). Often HgcA sequences extracted from environmental MAGs diverge from pure-culture HgcA from the same phyla. Metagenome-resolved HgcA sequences tend to cluster by environment, with HgcA from marine metagenomes often forming distinct subtrees from those sequenced from freshwater aquifers (Figure 7) (Rinke et al., 2013; Baker et al., 2015; Anantharaman et al., 2016; Bendall et al., 2016; Seitz et al., 2016; Probst et al., 2018; Tully et al., 2018). For example, several *Nitrospirae* HgcA-encoding genomes sequenced from aquifer groundwater and estuarine sediments either form their own distinct subtree or cluster with *Deltaproteobacteria* (Baker et al., 2015; Anantharaman et al., 2016). This may be indicative of the close relationship of some *Nitrospirae* to *Deltaproteobacteria* based on 16S rRNA gene analyses, as well as their cohabitation of sulfate-reducing environments (Teske et al., 1994).

Overall the majority of HgcA sequences branch from a cluster of HgcAB fused proteins related to *Thermococci, Atribacteria* (candidate division OP9), *Aminicenantes* (OP8), and *Chloroflexi* (Figure 7). The pattern of distribution and diversity amongst *hgcAB*+ microbes has been attributed to horizontal gene transfer in environments supporting syntrophic interactions, similar to the Woods Ljungdahl (WL) pathway (Podar et al., 2015a). Due to the discontinuity between *hgcA* and organism phylogeny, taxonomic identification from *hgcA* sequences alone are not conclusive. However, the similarity between clone, amplicon, or metagenomic read and representative *hgcA* sequences can estimate the prevalence and diversity of Hg-methylators through nearest neighbor analyses.

### 4.2 Classifying short-read *hgcA* sequences

Environmental samples from Hg-cycling environments (Table S3) were amplified with the new primers, cloned, trimmed, and aligned to both our 654 nt bp and 201 nt bp *hgcA* reference databases (Table S1). The reference database was compiled of *hgcA* genes from cultured, single cell genomes, and MAGs using taxonomy information from the NCBI database (Federhen, 2012). This provided a direct comparison of processing and classifying translated *hgcA* sequences over different sized portions of the gene. Generally, the predicted HgcA phylogeny from the 201 nt bp sequence agreed with the 654 nt bp classification (Figure 9). The environmental clones were primarily similar to HgcA from *Chloroflexi, Euryarchaeota, Firmicutes, Nitrospirae*, and *Proteobacteria* (Figure 9). There were notable disagreements in classification using different sized portions of the gene.

**Figure 9.**
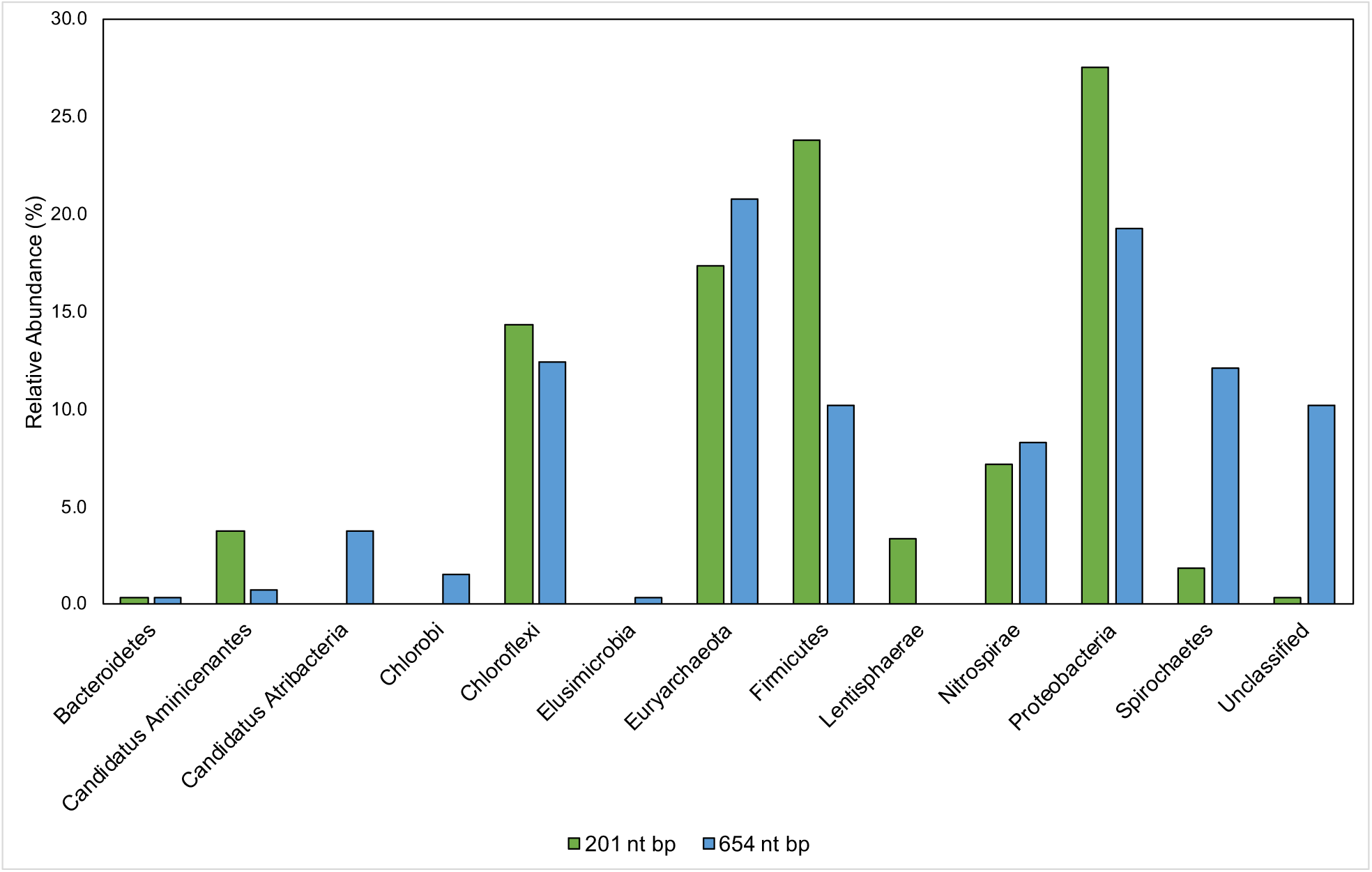
Comparison of taxonomic classifications of environmental *hgcA* clone sequences (n= 265) trimmed to 201 nt bp or 654 nt bp. Classifications were assigned to the translated *hgcA* clone sequences using ‘pplacer’ and ‘guppy classify’ with either the ‘HgcA_201.refpkg’ or ‘HgcA_654_full.refpkg’. Relative abundance of phylum level classifications are reported.

Classifications based on shorter read length may over-predict similarity to reference sequences. This is likely due to a decrease in pairwise identity from 54.3% to 36.1% across the translated 201 nt bp to 654 nt bp region in all included reference HgcA (excluding paralog sequences). Therefore, trimming a sequence from 654 to 201 bp could increase its pairwise identity to reference sequences, inflating confidence values. Approximately 10% of the clone sequences (n = 265) that were assigned taxonomy when only 201 nt bp was included, became unclassified when 654 nt bp was used for analysis. Decreasing the classification cut-off by 30% for the longer reads did not overcome this short-fall.

While the overall structure of the maximum likelihood trees of HgcA in both reference packages are similar, inclusion of a larger portion of the gene helps resolve subtrees that contain mixed phylogeny. For example, over the translated 201 nt bp portion of *hgcA*, phyla *Aminicenantes* and *Atribacteria* are 72% similar. But over the translated 654 nt bp portion of *hgcA*, similarity drops to 33%. Discrepancies in classification often occurred due to differences in resolution between reference trees. Classification of short sequences likely over estimated sequence similarity to *Aminicenantes, Lentisphaerae, Spirochaetes, Proteobacteria, Firmicutes* but under-estimated *Atribacteria, Chlorobi*, and *Elusimicrobia* (Figure 9).

### 4.3 Potential biases in identifying key Hg-methylators

Early *hgcA* primers could be biased towards *Deltaproteobacteria, Methanomicrobia*, and *Firmicutes* (Liu et al., 2014; Schaefer et al., 2014), often identifying *Deltaproteobacteria* and *Methanomicrobia* as dominant with less representation from *Firmicutes, Chloroflexi*, and unclassified clades (Liu et al., 2014; Schaefer et al., 2014; Du et al., 2017; Bravo et al., 2018a). While sulfate-reducing bacteria may be significant Hg-methylators (Compeau and Bartha, 1985; Kerin et al., 2006; Gilmour et al., 2018), it is unclear what role confirmation bias has had in their association with Hg cycling. With high-throughput *hgcAB* sequencing, one bias may be short sequence lengths (i.e. <300 bp), only partially capturing *hgcA* and *hgcB*. While platforms (e.g. PacBio, Nanopore) advance towards longer read capability (>1 kbp), error rates are often two orders of magnitude or greater compared to short reads (Ardui et al., 2018). Hence, reliance on these platforms (Liu et al., 2018b) should be approached with caution. However, the implicit error rate in PacBio sequencing can be corrected at least in part, via circular consensus sequencing, but the amplicon length must be 1/3-1/4 of the maximum read length in order to achieve higher quality reads (Francis et al., 2018). While low throughput, Sanger sequencing provides high quality sequencing of the entire amplicon and can validate the shorter read platforms, as performed herein but is unfortunately not routinely performed, eg. (Liu et al., 2018b).

Use of the new reference database increased diversity in previously published sequencing studies where classification used a smaller database. Biases may also be introduced during classification of translated *hgcA* sequences due to HgcA phylogeny being incongruent to species trees and limited reference sequences. Therefore including new *hgcAB*^+^ organism discoveries are critical (Parks et al., 2013) since new reference sequences may identify ‘unclassified’ *hgcA* sequences. Using an outdated database for *hgcA* classification can underestimate Hg-methylator diversity. A recent study used PacBio *hgcAB* amplicon and Illumina MiSeq metagenomic sequencing from rice paddies without Sanger sequencing validation or using the circular consensus protocol to correct the PacBio data, and indicated that the Hg-methylating community was dominated by *Deltaproteobacteria* and *Methanomicrobia*, with some *Firmicutes* and *Chloroflexi* (Liu et al., 2018b). Additionally, taxonomic classifications were performed with an outdated (65 *hgcAB* sequences) database limited to *Deltaproteobacteria, Methanomicrobia, Firmicutes*, and *Chloroflexi* (Liu et al., 2018b). We applied our 201 nt bp database to the publicly available metagenomic reads, since the amplicon data were not made public, and found that *hgcA* closely related to *Deltaproteobacteria, Methanomicrobia, Chloroflexi*, and *Firmicutes* were present but also present were *Aminicenantes, Atribacteria*, candidate division WOR-1, *Chlorobi, Thermococci, Oligoflexi, Nitrospirae, Spirochaetes, Lentisphaerae, Bacteroidetes* and *Elusimicrobia* and unclassified Bacteria and Euryarchaeota (Figure 10). This represents a 25.4-37.1% increase in diversity. We also found that *hgcA* were dominated by *Deltaproteobacteria* (39-49%) compared to their findings (40%). However, *Methanomicrobia* did not comprise ∼60% of the *hgcA* community, but rather was 5.1-10.5%, similar to *Chloroflexi* (7.2-15.1%), *Firmicutes* (5.4-10.2%) and *Nitrospirae* (3.1-9.1%). While *Geobacter* spp. reportedly comprised ∼34% of the communities with low *hgcAB*^+^ sulfate-reducer abundances, our analyses showed *Geobacter* spp. (2.8-12.9%) and sulfate-reducers were equivalent (3.6-6.1%), suggesting that both Fe(III)- and sulfate-reduction are important and both contribute to Hg-methylation. Going forward it is imperative that such biases are recognized since the implications of incorrect data analysis and interpretation can lead to erroneous conclusions. Should these data be used in predictive models, the models would most likely be incorrect, invalidating the model for less than obvious reasons, and likely leading investigators down the wrong path.

**Figure 10.**
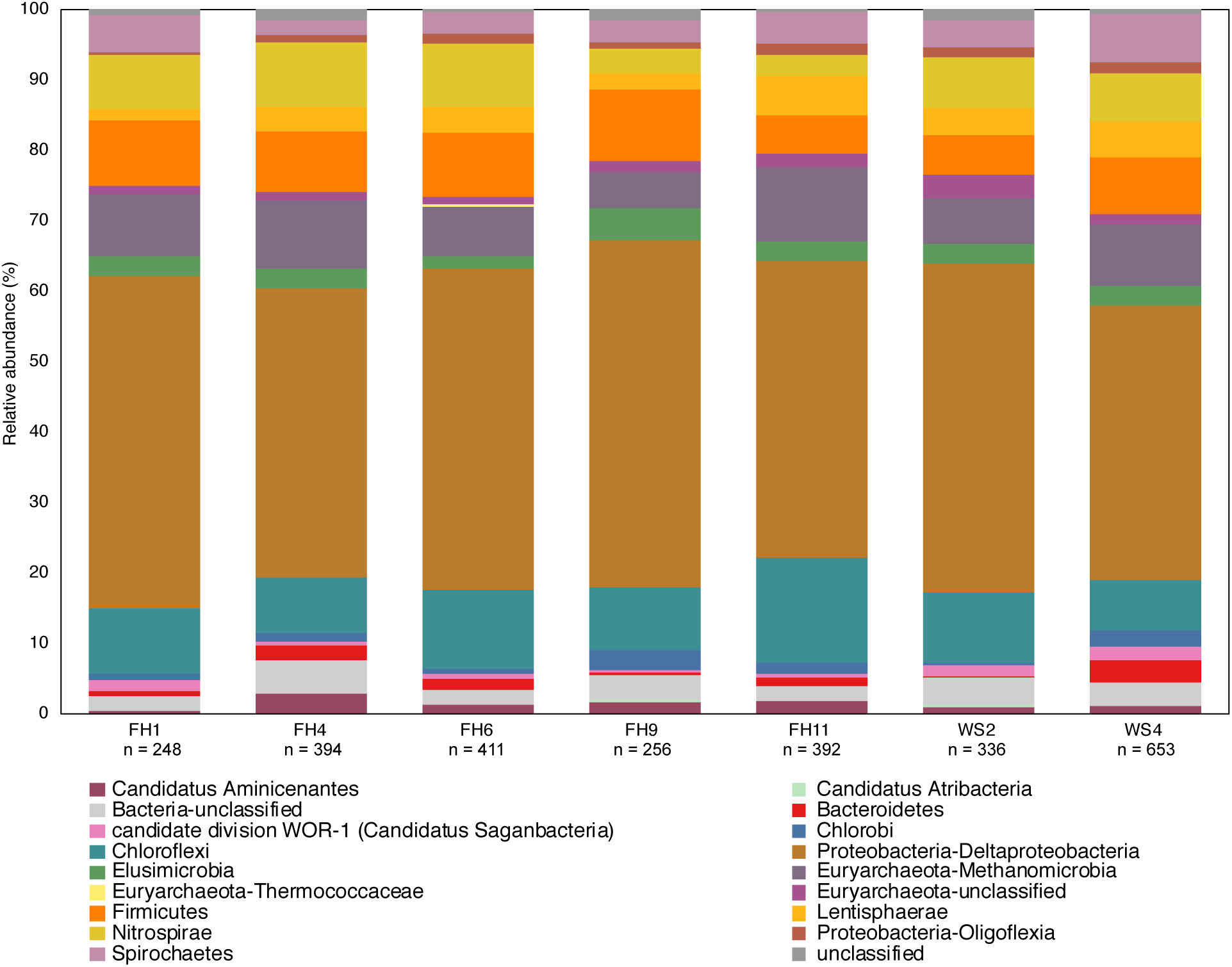
Phylogenetic composition of *hgcA* genes from published rice paddy metagenomes (Genbank PRJNA450451) at the phylum and class level (Liu et al., 2018b). The *hgcA* genes were identified in raw sequence reads using the hmm profile from the in-house reference package (HgcA_201.refpkg) with an inclusion value 1E-7. Taxonomic classifications were made using the in-house reference package with a confidence cut-off of 90%. Relative abundance of *hgcA* genes normalized to total *hgcA* genes (n) that aligned to reference package in each metagenome.

Critically, the revised primer set and protocol outlined herein improves our ability to capture the *hgcAB* diversity in shotgun metagenomic analysis of environmental samples (Figure 7). Both primer-sets captured *hgcA* related to the *Deltaproteobacteria, Firmicutes*, and *Methanomicrobia* as well as clades representing novel potential Hg-methylators including *Nitrospirae, Phycisphaerae, Aminicenantes, Spirochaetes, Chloroflexi*, and *Elusimicrobia*. We conclude that the original reverse primer degeneracy (Christensen et al., 2016) is not necessary. Amplification efficiencies from environmental samples have been improved with the new primers and can now be used directly with high-throughput sequencing to achieve ∼100,000 *hgcAB* amplicons per sample. The ability to capture and directly sequence highly diverse *hgcAB* amplicons from environments will advance our understanding of Hg-methylator distribution across phylogeny and environment. Being able to correctly identify a broad-range of Hg-methylators is critical to understanding the key players in MeHg production and accumulation in aquatic ecosystems. Knowing which organisms are most strongly contributing to Hg methylation in a given environment is crucial for constructing an ecosystem-level biogeochemical model for Hg methylation as well as designing strategies for mitigating MeHg production.

## 6 Author Contributions

Primer design and testing was done by CMG, with assistance from AMW, DE, MP, based on modifications to earlier designs by GC, DE, AW, and MP. High-throughput sequencing method development by DSJ. Environmental DNA extraction, PCR, and cloning method development and testing by AMW, MML, GCC, RLW, and CMG. Model communities and DNA extractions from them were provided by AS and CCG. CMG did the primary writing with assistance from RW, DE, AMW, and CCG

## 8 Acknowledgments

We wish to thank and acknowledge the staff at the University of Minnesota Genomics Center (UMGC) for advice and assistance with high-throughput sequencing of custom *hgcAB* amplicon libraries. We also thank Brandy Toner (University of Minnesota), for peat, hollow and hummock samples from SPRUCE, Helen Hsu-Kim for sediment samples from Sandy Creek, NC, and Baohua Gu for rice paddy soils from Yanwuping, China.

## Figure Legends

**Figure 1**. Alignment showing the highly conserved 20 nt bp region of *hgcB* where the reverse primer binds. Alignment is based on 239 reference *hgcB* sequences, however only the top 50 sequences most divergent from the consensus sequence are shown. Bases that disagree with consensus sequence are highlighted. Reverse complements of each primer (ORNL-HgcAB-uni-R, ORNL-HgcAB-uni-32R) are included for reference. The bar graph at the top of the image shows mean pairwise identity for each base across all sequences – green is 100%, green-brown is between 30 and 100%, and red is <30% conserved in 239 sequences. Y = C/T, R = G/A, V = C/G/A, S = C/G, N = A/T/C/G.

**Figure 2**. Percent occurrence of each 96-degenerate reverse (top; ORNL-HgcAB-uni-R) and 32-degenerate reverse primer (bottom; ORNL-HgcAB-uni-32R) oligonucleotide sequence in the environmental clone *hgcAB* sequences and Miseq amplicons. Primers are listed in Table S2. Matches to each broad-range reverse primer (ORNL-HgcAB-uni-R, ORNL-HgcAB-uni-32R) are shown for each set of clone and amplicon sequences amplified with the ORNL-HgcAB-uni-R (A) and ORNL-HgcAB-uni-32R (B). Oligonucleotide sequences shared between each reverse primer are indicated with asterisk (*). Reverse primer sequences were found in 95 clones (A) and 369 clones (B), and 210, 362 (A) and 233, 934 (B) amplicon reads. Data is shown on a log scale. Dashed line indicates equal distribution of reverse primer sequences across amplicon sequences: A) 1.04%, B) 3.13%. Ordered by decreasing abundance in Miseq amplicon libraries.

**Figure 3**. Principle coordinate analyses (PCoA) showing the variance in phylogenetic distances between mock community (A) and New Horizon (NH) sediment (B) *hgcAB* amplicon libraries. Reverse primer (i.e. ORNL-HgcAB-uni-R, ORNL-HgcAB-uni-32R) is indicated by shape. Weighted UniFrac distances calculated from the taxonomic classifications of clustered (90%) OTUs using the program ‘pplacer’ and the R package ‘phyloseq’.

**Figure 4**. Comparison of expected and actual taxonomic composition of *hgcA* sequences in recovered mock community amplicon libraries. Top, mock community Combo1; bottom, mock community Combo2. Libraries were constructed using the newer less degenerate *hgcAB* reverse primer (left) or the Christensen 2016 reverse primer (right). Comparison of various clustering methods: **Method A**: cluster nucleotide sequences at 90% identity using VSEARCH; **Method B:** cluster peptide sequences at 90% identity, using USEARCH; **Method C:** clustering nucleotide sequences at 90% using cd-hit-est; **Method D:** chimeras removed prior to clustering nucleotide sequences at 90% using cd-hit-est.

**Figure 5**. Relative abundance of cloned and amplicon *hgcAB* sequences by phyla and class. *hgcAB* genes were amplified from New Horizon sediment (NH3) and SPRUCE soil (SPRUCE1) using ORNL-HgcAB-uni-F with both reverse primers (ORNL-HgcAB-uni-R, ORNL-HgcAB-uni-32R). Taxonomic classifications based on phylogenetic placement of representative sequences on the reference tree (‘ORNL_HgcA_201.ref.pkg’) using program Taxtastic with classification cutoff of 90%. OTUs have been merged at the class-level, with relative abundance values calculated from proportion of total clone of amplicon sequences in each library. Figure produced using R packages ‘phyloseq’ and ‘ggplot2’. *hgcA* amplicons were not sampled (n.s.) from SPRUCE soils.

**Figure 6**. Relative abundance profiles of *hgcA* amplicons from mock (A) and environmental (B) NH sediment samples at the phyla level. OTUs were clustered at 90% sequence similarity, and taxonomic classifications were assigned based on phylogenetic placement of representative sequences on the ‘ORNL_HgcA_201.ref.pkg’ reference tree using the program ‘pplacer’ with a classification cut-off of 90%. OTUs have been merged at the phyla level.

**Figure 7**. Molecular phylogenetic analysis of HgcA reference sequences by Maximum Likelihood method. Phylogeny was calculated using the GAMMA model of rate heterogeneity and ML estimate of alpha-parameter in RAxML with 150 bootstrap replicates. The tree was rooted by carbon monoxide dehydrogenases from non-Hg-methylators *Candidatus Omnitrophica bacterium* CG1-02-41-171 and *Thermosulfurimonas dismutans*. Relative abundance of *hgcA* amplicons from Tidal Marsh sediment (1064) and East Fork Poplar Creek NH sediments were calculated from OTU clustering at 90%, and the values placed on tree using ‘pplacer’. Phylum-level classifications of HgcA reference sequences are color-coded, and distinct subtrees are labeled, including metagenome-assembled genomes from Rifle Aquifer Field Research Site and marine datasets. ‘Other’ phyla include phyla with low number of HgcA representatives: *Planctomycetes, Nitrospina*, and Candidate phyla *Raymondbacteria*, WOR-1, WOR-3, and *Wallibacteria*. Gray background indicates reference sequences are from metagenome-assembled genomes (MAGs) or environmental single cell genome center sequencing (SCGC) datasets. Red branches indicate that sequences were included in phylogenetic analysis from Podar et al.(Podar et al., 2015a). Black branches are sequences that have since been identified.

**Figure 8**. Known and novel methylators in a sediment core from East Fork Poplar Creek (Oak Ridge, TN). Relative abundance of *hgcA* phyla across a 12cm sediment core were identified using the new degenerate primers, direct sequencing technique, and the updated *hgcA* database. Known methylators (*Deltaproteobacteria, Firmicutes-Clostridia*, and *Euryarchaeota*-*Methanomicrobia*) were recovered as well as novel potential methylators.

**Figure 9**. Comparison of taxonomic classifications of environmental *hgcA* clone sequences (n= 265) trimmed to 201 nt bp or 654 nt bp. Classifications were assigned to the translated *hgcA* clone sequences using ‘pplacer’ and ‘guppy classify’ with either the ‘HgcA_201.refpkg’ or ‘HgcA_654_full.refpkg’. Relative abundance of phylum level classifications are reported.

**Figure 10**. Phylogenetic composition of *hgcA* genes from published rice paddy metagenomes (Genbank PRJNA450451) at the phylum and class level (Liu et al., 2018b). The *hgcA* genes were identified in raw sequence reads using the hmm profile from the in-house reference package (HgcA_201.refpkg) with an inclusion value 1E-7. Taxonomic classifications were made using the in-house reference package with a confidence cut-off of 90%. Relative abundance of *hgcA* genes normalized to total *hgcA* genes (n) that aligned to reference package in each metagenome.

