## Supplementary Materials for "An improved *hgcAB* primer set and direct high-throughput sequencing expand Hg-methylator diversity in nature"

### Supplementary Material

#### 1 Supplementary Figures and Tables

##### 1.1 Supplementary Figures

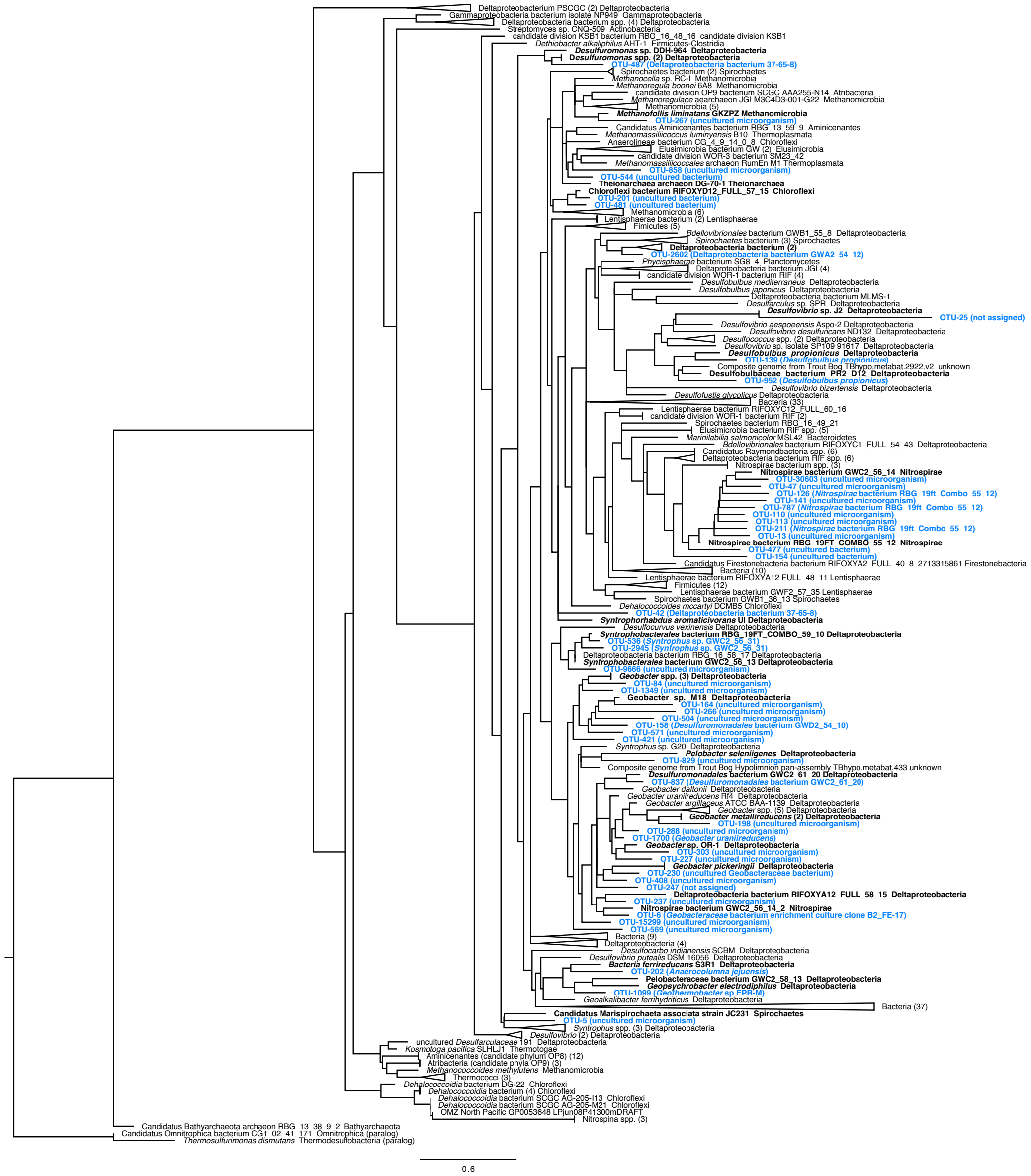

**Supplementary Figure 1**. Maximum likelihood tree of *hgcA* sequences from the updated reference package showing pplacer placements of the top 50 OTUS from New Horizon (NH) *hgcA* amplicon sequences (Matsen et al., 2010). Nearest neighbor shown in bold. NH *hgcA* sequences are shown in blue with classification based on the lowest common ancestor (LCA, 50 % cut-off) of protein BLAST hits to NCBI non-redundant protein database (Altschul et al., 1990); LCA calculated in MEGAN v6 (Huson et al., 2016).

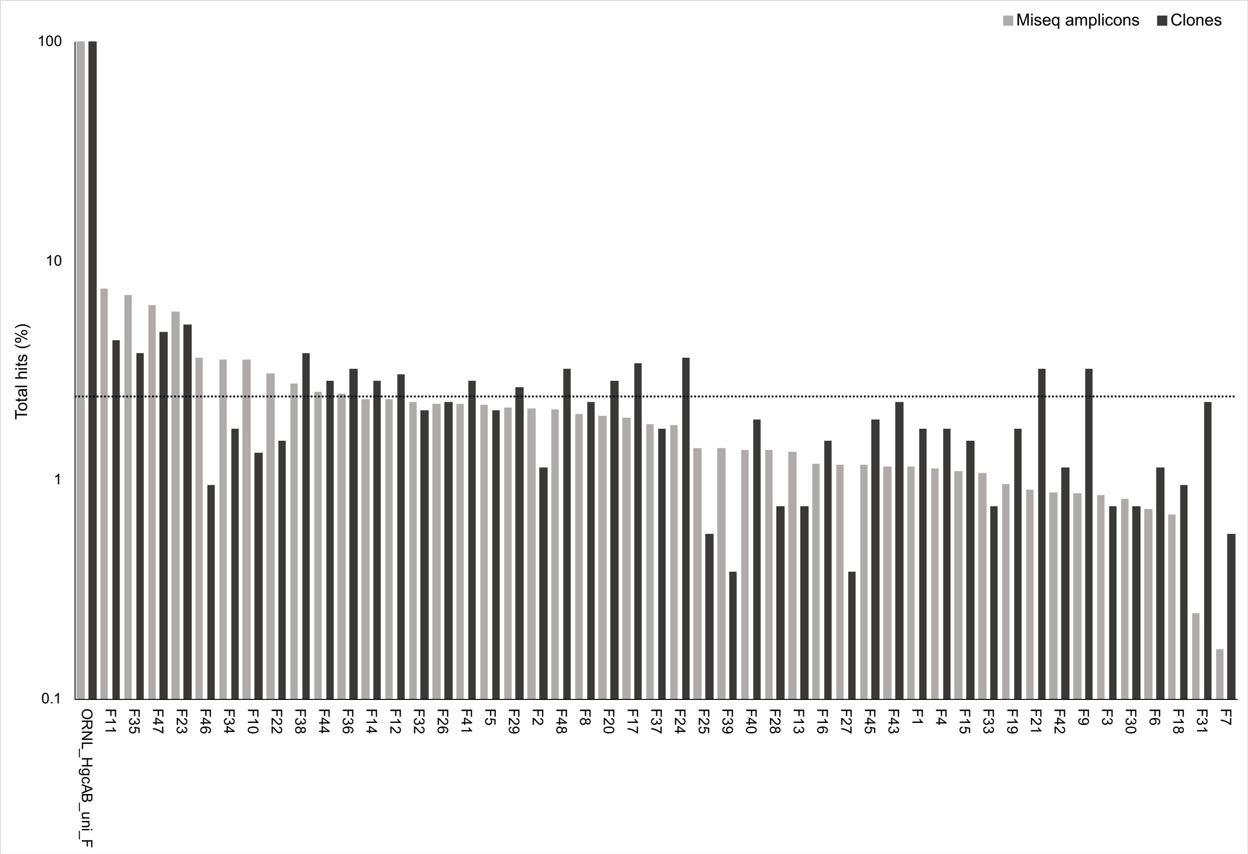


**Supplementary Figure 2.** Percent occurrence of each of the 48-degenerate forward (ORNL-HgcAB-uni-F) oligonucleotide sequence in the environmental clone *hgcAB* sequences (Table S2) and Miseq *hgcAB* amplicons from sites NH. Clones were generated with both the Christensen 2016 primer set (ORNL-HgcAB-uni-F, ORNL-HgcAB-uni-R; n = 119) and the new less degenerate primer set (ORNL-HgcAB-uni-F, ORNL-HgcAB-uni-32R; n = 509). Forward primer binding sites were found in 531 clones of the total 628 clone library. A subset of sequences were pulled from amplicon libraries produced from six separate NH sediment samples using ORNL-HgcAB-uni-F and ORNL-HgcAB-uni-R (n = 87,500) and ORNL-HgcAB-uni-F and ORNL-HgcAB-uni-32R (n = 87,500). Forward primer binding sites were found in 166,984 of the 175,000 amplicon sequences searched. Primers are listed in Table S2. Data is shown on a log scale. Dashed line indicates equal distribution of forward primer sequences across amplicon sequences: 2.08%.

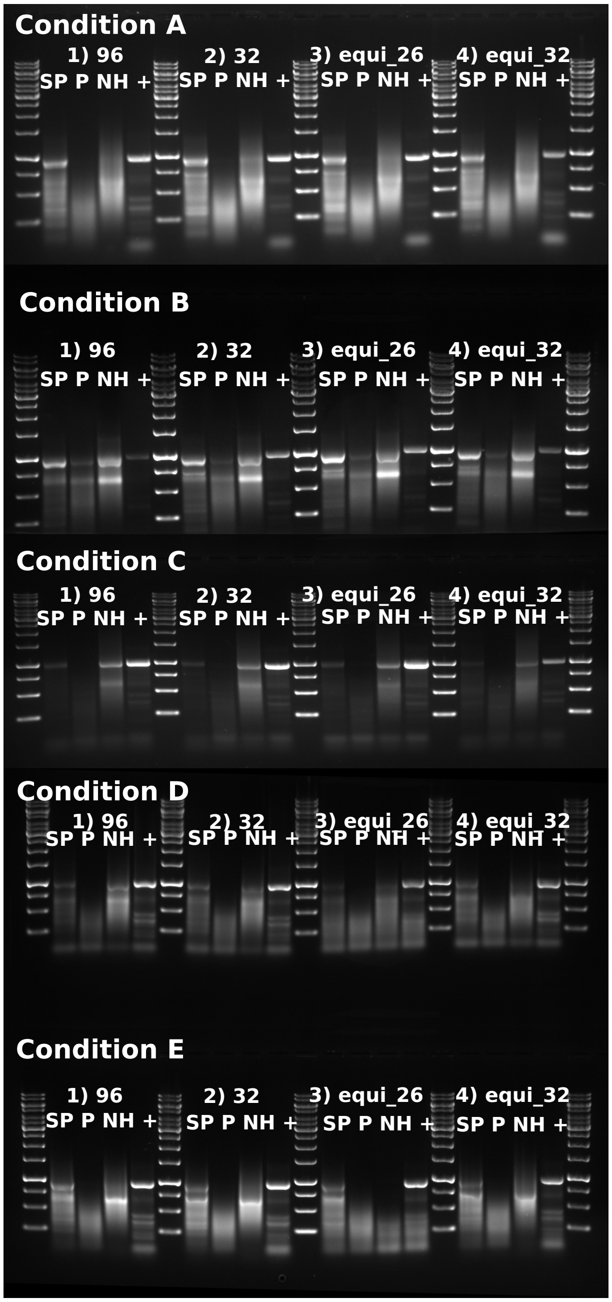


**Supplementary Figure 3.** Gel image showing PCR amplification of *hgcAB* from SPRUCE soil (SP), NH periphyton (P), and NH sediment (NH) using PCR conditions A—E (Table S4) with 96 degenerate primer set (ORNL-HgcAB-uni-F, ORNL-HgcAB-uni-R), 32 degenerate primer set (ORNL-HgcAB-uni-F, ORNL-HgcAB-uni-R32), and equimolar reverse primer mixes of ORNL-HgcAB-uni-R26 and ORNL-HgcAB-uni-R32. *Desulfovibrio desulfuricans* ND132 *hgcAB* amplicon (~980bp) was included as a positive (+) control, compared against a 1 kbp GeneRuler ladder.

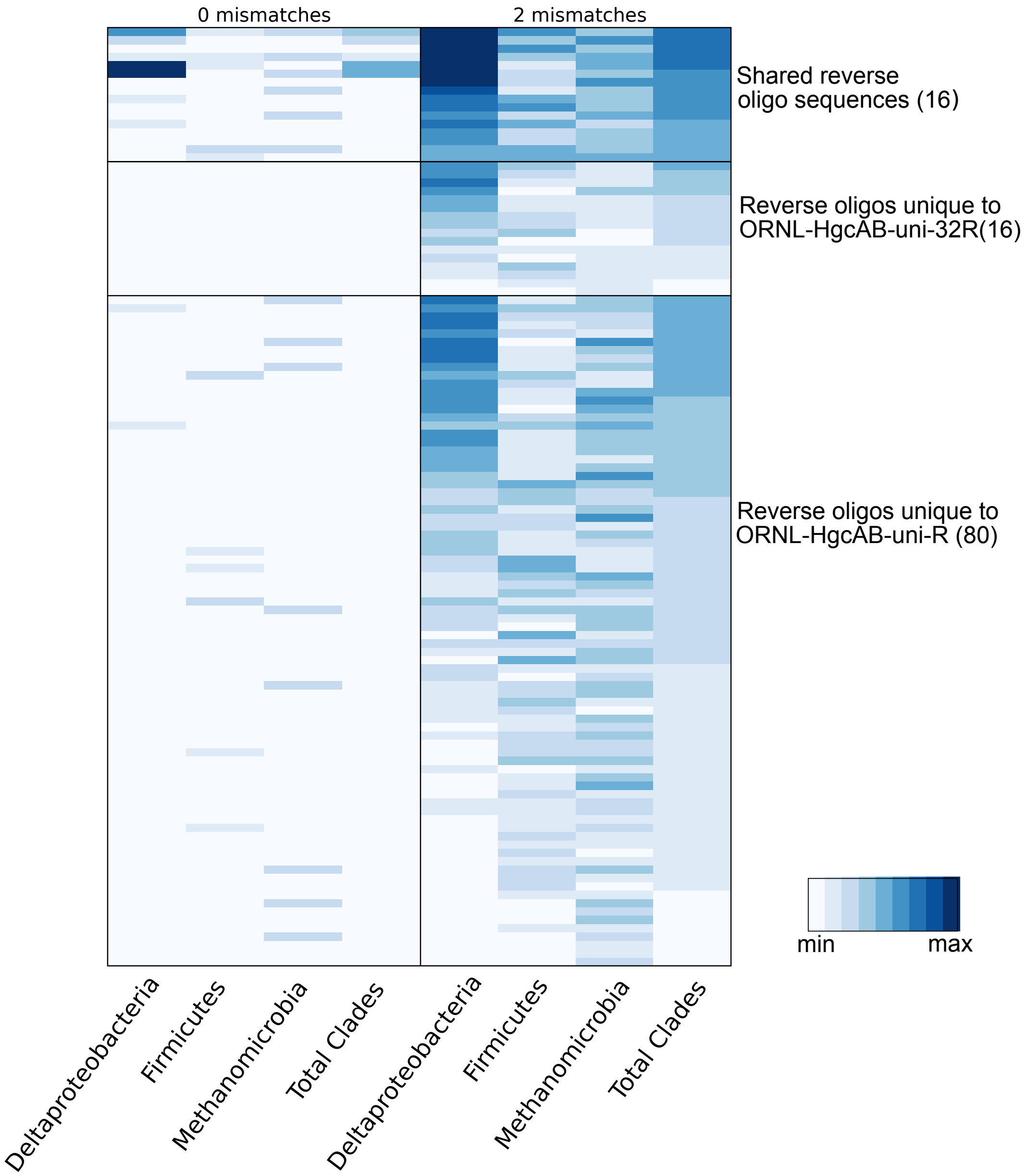


**Supplementary Figure 4.** Heatmap showing the percentage that each possible sequence from the reverse broad-range primer from this study (ORNL-HgcAB-uni-32R) compared to the more degenerate primer from previous study (ORNL-HgcAB-uni-R; Christensen et al. 2016) align with *hgcAB* from reference sequences (listed in Table S1) of the three major clades: *Deltaproteobacteria* (124), *Fimicutes* (32), *Methanomicrobia* (15), and total sequences (239). Alignments allowed for either no mismatches (on the left), and up to 2 mismatches (on the right). The colorbar is scaled based on the minimum and maximum values for the 0-mismatch and 2-mismatch sets separately (0–19% and 1–85%, respectively).

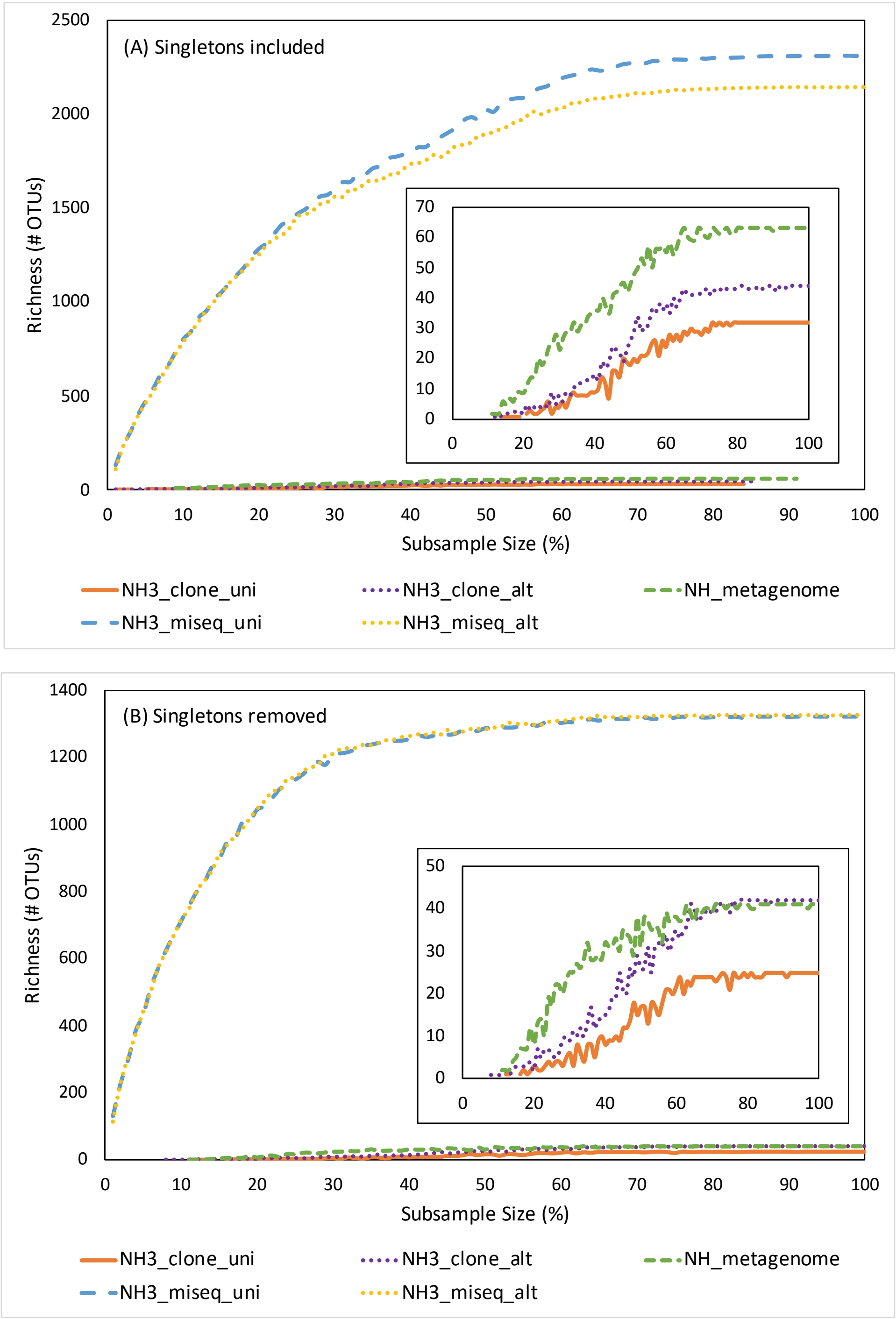


**Supplementary Figure 5.** Rarefraction curves assessing OTU richness in *hgcA* sequences from NH sediment (East Fork Poplar Creek, Oak Ridge, TN) clone, amplicon, and metagenomic datasets. The clone and amplicon sequences were amplified from sample NH3 with the reverse primer from this study ('uni-32R') compared to the more degenerate primer from previous study ('uni-R';(Christensen et al., 2016)). Included are *hgcA* sequences pulled from a NH sediment metagenomic dataset from a previous study (Christensen et al., 2019). OTU richness was calculated using USEARCH with singletons included (A) and discarded (B) prior to OTU clustering (Edgar, 2010).

#### 1.2 Supplementary Tables

**Supplementary Table 1.** List of *hgcAB*+ organisms used in reference package and for *in silico* analyses. Sequences were selected from publicly available genomes from the National Center for Biotechnology Information (http://www.ncbi.nlm.nih.gov), including Hg-methylating organisms, shotgun metagenomics, and metagenome-assemble microbial genomes (MAGS). This list, including *hgcA* and *hgcB* nucleotide sequences, is available online through the DOE Data Explorer (Gionfriddo et al., 2019).
* indicates only an *hgcA* sequence is available for this organism.

*Acetivibrio cellulolyticus* CD2 DSM 1870 (*Firmicutes-Clostridia*)
*Acetonema longum* DSM 6540 (*Firmicutes-Clostridia*)
*Alkaliphilus peptidifermentans* DSM 18978 (*Firmicutes-Clostridia*)
*Anaerolineae* bacterium CG_4_9_14_0_8 (*Chloroflexi*)
*Bacteria ferrireducans* S3R1 (*Deltaproteobacteria*)
*Bacteroides cellulosolvens* DSM 2933 (*Bacteroidetes*)
*Bacteroides* sp. SM1 62 (*Bacteroidetes*)
*Bacteroides* sp. SM23 62 (*Bacteroidetes*)
*Bacteroidetes* bacterium GWA2_32_17 (*Bacteroidetes*)
*Bacteroidetes* bacterium GWF2_35_48 (*Bacteroidetes*)
*Bacteroidetes* bacterium RIFOXYA12_FULL_35_11 (*Bacteroidetes*)
*Bacteroidetes* bacterium RIFOXYC12_FULL_35_7 (*Bacteroidetes*)
*Bdellovibrionales* bacterium GWB1_55_8 (*Deltaproteobacteria*)
*Bdellovibrionales* bacterium RIFOXYB1_FULL_39_21 (*Deltaproteobacteria*)*
*Bdellovibrionales* bacterium RIFOXYC1_FULL_39_130 (*Deltaproteobacteria*)*
*Bdellovibrionales* bacterium RIFOXYC1_FULL_54_43 (*Deltaproteobacteria*)
*Bdellovibrionales* bacterium RIFOXYC12_FULL_39_17 (*Deltaproteobacteria*)*
*Bdellovibrionales* bacterium RIFOXYD12_FULL_39_22 (*Deltaproteobacteria*)*
candidate division KSB1 bacterium RBG_16_48_16 (candidate division KSB1)
candidate division OP8 bacterium SCGC_AAA252-A02 (*Aminicenantes*)*
candidate division OP8 bacterium SCGC_AAA252-F08 (*Aminicenantes*)*
candidate division OP8 bacterium SCGC_AAA252-G05 (*Aminicenantes*)*
candidate division OP8 bacterium SCGC_AAA252-G06 (*Aminicenantes*)*
candidate division OP8 bacterium SCGC_AAA252-J09 (*Aminicenantes*)*
candidate division OP8 bacterium SCGC_AAA252-J21 (*Aminicenantes*)*
candidate division OP8 bacterium SCGC_AAA252-K07 (*Aminicenantes*)*
candidate division OP8 bacterium SCGC_AAA252-O09 (*Aminicenantes*)*
candidate division OP8 bacterium SCGC_AAA252-O19 (*Aminicenantes*)*
candidate division OP8 bacterium SCGC_AAA252-P13 (*Aminicenantes*)*
candidate division OP8 bacterium SCGC_AAA252-P19 (*Aminicenantes*)*
candidate division OP8 bacterium SCGC_AAA255-O15 (*Aminicenantes*)

candidate division OP9 bacterium SCGC AAA252-M02 (*Atribacteria*)*
candidate division OP9 bacterium SCGC AAA255-N14 (*Atribacteria*)
candidate division OP9 bacterium SCGC AB-164-L03 (*Atribacteria*)*
candidate division OP9 bacterium SCGC AB-164-P05 (*Atribacteria*)*
candidate division WOR-1 bacterium RIFOXYA2_FULL_41_14 (candidate division WOR-1)
candidate division WOR-1 bacterium RIFOXYA2_FULL_46_56 (candidate division WOR-1)
candidate division WOR-1 bacterium RIFOXYA12_FULL_43_27 (candidate division WOR-1)
candidate division WOR-1 bacterium RIFOXYB2_FULL_42_35 (candidate division WOR-1)
candidate division WOR-1 bacterium RIFOXYB2_FULL_46_45 (candidate division WOR-1)
candidate division WOR-1 bacterium RIFOXYC2_FULL_46_14 (candidate division WOR-1)
candidate division WOR-3 bacterium SM23_42 (candidate division WOR-3)
Candidatus *Aminicenantes* bacterium RBG_13_59_9 (*Aminicenantes*)
Candidatus *Bathyarchaeota* archaeon RBG_13_38_9_2 (*Bathyarchaeota*)*
Candidatus *Desulfuromonas soudanensis* WTL (*Deltaproteobacteria*)
Candidatus *Firestonebacteria* bacterium RIFOXYA2_FULL_40_8 (*Firestonebacteria*)
Candidatus *Marispirochaeta associata* strain JC231 (*Spirochaetes*)
Candidatus *Methanoregula boonei* 6A8 (*Methanomicrobia*)
Candidatus *Methanosphaerula palustris* E1-9c (*Methanomicrobia*)
Candidatus *Raymondbacteria* bacterium RIFOXYA12_full_50_37 (*Raymondbacteria*)
Candidatus *Raymondbacteria* bacterium RIFOXYA2_FULL_49_16 (*Raymondbacteria*)
Candidatus *Raymondbacteria* bacterium RIFOXYB12_full_50_8 (*Raymondbacteria*)
Candidatus *Raymondbacteria* bacterium RIFOXYB2_FULL_49_35 (*Raymondbacteria*)
Candidatus *Raymondbacteria* bacterium RifOxyC12_full_50_8 (*Raymondbacteria*)
Candidatus *Raymondbacteria* bacterium RIFOXYD12_FULL_49_13 (*Raymondbacteria*)
Candidatus *Wallbacteria* bacterium GWC2_49_35 (*Wallbacteria*)*
*Chloroflexi* bacterium RIFOXYD12_FULL_57_15 (*Chloroflexi*)
*Clostridium cellobioparum* ATCC 15832 (*Firmicutes-Clostridia*)
*Clostridium cellulosi* CS-4-4 (*Firmicutes-Clostridia*)*
clostridium Ga0073690 (*Firmicutes-Clostridia*)
*Clostridium jejuense* DSM 15929 (Firmicutes-Clostridia)
*Clostridium litorale* W6 DSM 5388 (Firmicutes-Clostridia)
*Clostridium* sp. 3 Draft 3 (*Firmicutes-Clostridia*)
*Clostridium termitidis* CT1112 DSM 5398 (*Firmicutes-Clostridia*)
*Clostridium tunisiense* TJ (*Firmicutes-Clostridia*)
*Clostridium xylanovorans* DSM 12503 (*Firmicutes-Clostridia*)
Composite genome from Trout Bog Hypolimnion pan-assembly TBhypo.metabat.433 (unknown)
Composite genome from Trout Bog Hypolimnion TBhypo.metabat.3815 (unknown)
Composite genome from Trout Bog TBhypo.metabat.2922.v2 (unknown)
Composite genome from Trout Bog TBhypo.metabat.5247 (unknown)
*Dehalobacter restrictus* DSM 9455 (*Firmicutes-Clostridia*)

*Dehalobacter* sp. 11DCA (*Firmicutes-Clostridia*)
*Dehalobacter* sp. CF (*Firmicutes-Clostridia*)
*Dehalobacter* sp. UNSWDHB (*Firmicutes-Clostridia*)
*Dehalococcoides mccartyi* DCMB5 (*Chloroflexi*)
*Dehalococcoidia* bacterium DG 22 (*Chloroflexi*)*
*Dehalococcoidia* bacterium SCGC_AG-205-B13 (*Chloroflexi*)*
*Dehalococcoidia* bacterium SCGC_AG-205-I02 (*Chloroflexi*)*
*Dehalococcoidia* bacterium SCGC_AG-205-I13 (*Chloroflexi*)*
*Dehalococcoidia* bacterium SCGC_AG-205-K13 (*Chloroflexi*)*
*Dehalococcoidia* bacterium SCGC_AG-205-M10 (*Chloroflexi*)*
*Dehalococcoidia* bacterium SCGC_AG-205-M21 (*Chloroflexi*)*
delta proteobacterium MLMS-1 (*Deltaproteobacteria*)
Delta proteobacterium NaphS2 (*Deltaproteobacteria*)
delta proteobacterium PSCGC 5419 (*Deltaproteobacteria*)
delta proteobacterium PSCGC 5451 (*Deltaproteobacteria*)
*Deltaproteobacteria* bacterium GWA2_55_10 (*Deltaproteobacteria*)
*Deltaproteobacteria* bacterium isolate ARS66 (*Deltaproteobacteria*)*
*Deltaproteobacteria* bacterium isolate NP36 76525 (*Deltaproteobacteria*)*
*Deltaproteobacteria* bacterium isolate NP36 (*Deltaproteobacteria*)*
*Deltaproteobacteria* bacterium isolate SP3084 (*Deltaproteobacteria*)*
*Deltaproteobacteria* bacterium JGI A06048-F13 (*Deltaproteobacteria*)
*Deltaproteobacteria* bacterium JGI E06040-H20 (*Deltaproteobacteria*)
*Deltaproteobacteria* bacterium RBG_16_44_11 (*Deltaproteobacteria*)
*Deltaproteobacteria* bacterium RBG_16_58_17 (*Deltaproteobacteria*)
*Deltaproteobacteria* bacterium RBG_19FT_COMBO_43_11 (*Deltaproteobacteria*)
*Deltaproteobacteria* bacterium RIFCSPHIGHO2_02_FULL_42_44 (*Deltaproteobacteria*)
*Deltaproteobacteria* bacterium RIFCSPHIGHO2_02_FULL_43_33 (*Deltaproteobacteria*)
*Deltaproteobacteria* bacterium RIFCSPLOWO2_01_FULL_42_9 (*Deltaproteobacteria*)*
*Deltaproteobacteria* bacterium RIFCSPLOWO2_02_FULL_42_39 (*Deltaproteobacteria*)
*Deltaproteobacteria* bacterium RIFCSPLOWO2_02_FULL_55_12 (*Deltaproteobacteria*)
*Deltaproteobacteria* bacterium RIFCSPLOWO2_12_FULL_43_16 (*Deltaproteobacteria*)*
*Deltaproteobacteria* bacterium RIFOXYA2_FULL_42_10 (*Deltaproteobacteria*)
*Deltaproteobacteria* bacterium RIFOXYA12_FULL_58_15 (*Deltaproteobacteria*)
*Deltaproteobacteria* bacterium RIFOXYD12_FULL_50_9 (*Deltaproteobacteria*)
*Deltaproteobacteria* bacterium RIFOXYD12_FULL_53_23 (*Deltaproteobacteria*)
*Deltaproteobacteria* bacterium RIFOXYD12_FULL_56_24 (*Deltaproteobacteria*)
*Deltaproteobacteria* bacterium RIFOXYD12_FULL_57_12 (*Deltaproteobacteria*)
*Deltaproteobacteria* bacterium SG8 13 (*Deltaproteobacteria*)
*Deltaproteobacteria* bacterium SM23 61 (*Deltaproteobacteria*)
Deltaproteobacterium sp. OalgD1a (*Deltaproteobacteria*)

Deltaproteobacterium sp. OalgD1b (*Deltaproteobacteria*)
Deltaproteobacterium sp. OalgD3 (*Deltaproteobacteria*)
Deltaproteobacterium sp. OalgD4 (*Deltaproteobacteria*)
*Desulfacinum hydrothermale* DSM 13146 (*Deltaproteobacteria*)
*Desulfacinum infernum* DSM 9756 (*Deltaproteobacteria*)
*Desulfarculus* sp. SPR (*Deltaproteobacteria*)
*Desulfitobacterium dehalogenans* JWIU-DC1 ATTC 51507 (*Firmicutes-Clostridia*)
*Desulfitobacterium dichloroeliminans* LMG P21439 (*Firmicutes-Clostridia*)
*Desulfitobacterium metallireducens* 853-15A DSM 15288 (*Firmicutes-Clostridia*)
*Desulfitobacterium* sp. PCE1 DSM 10344 (*Firmicutes-Clostridia*)
*Desulfobacter* sp. isolate ARS36 (*Deltaproteobacteria*)
*Desulfobacterales* bacterium RIFOXYA12_FULL_46_15 (*Deltaproteobacteria*)
*Desulfobacterales* bacterium SG8_35_2 (*Deltaproteobacteria*)
*Desulfobacterium* vacuolatum DSM 3385 (*Firmicutes-Clostridia*)
*Desulfobacula phenolica* DSM 3384 (*Deltaproteobacteria*)
*Desulfobacula* sp. TS (*Deltaproteobacteria*)
*Desulfobulbaceae* bacterium PR2 D12 (*Deltaproteobacteria*)
*Desulfobulbus japonicus* DSM 18378 (*Deltaproteobacteria*)
*Desulfobulbus mediterraneus* DSM 13871 (*Deltaproteobacteria*)
*Desulfobulbus propionicus* DSM 2032 (*Deltaproteobacteria*)
*Desulfobulbus* sp. Tol-SR (*Deltaproteobacteria*)
*Desulfocarbo indianensis* SCBM (*Deltaproteobacteria*)
*Desulfococcus biacutus* KMRActS (*Deltaproteobacteria*)
*Desulfococcus multivorans* DSM 2059 (*Deltaproteobacteria*)
*Desulfocurvus vexinensis* DSM 17965 (*Deltaproteobacteria*)
*Desulfofustis glycolicus* DSM 9705 (*Deltaproteobacteria*)
*Desulfoluna spongiiphila* AA1 (*Deltaproteobacteria*)
*Desulfomicrobium apsheronum* DSM 5918 (*Deltaproteobacteria*)
*Desulfomicrobium baculatum* DSM 4028 (*Deltaproteobacteria*)
*Desulfomicrobium escambiense* DSM 10707 (*Deltaproteobacteria*)
*Desulfomicrobium norvegicum* DSM 1741 (*Deltaproteobacteria*)
*Desulfomonile* sp. Ga0081660 (*Deltaproteobacteria*)*
*Desulfomonile tiedjei* DCB-1 DSM 6799 (*Deltaproteobacteria*)
*Desulfonatronospira thiodismutans* ASO3-1 (*Deltaproteobacteria*)
*Desulfonatronovibrio hydrogenovorans* DSM 9292 (*Deltaproteobacteria*)
*Desulfonatronum lacustre* Z-7951 DSM 10312 (*Deltaproteobacteria*)
*Desulfonatronum thioautotrophicum* ASO4-1 (*Deltaproteobacteria*)
*Desulfonatronum thiodismutans* MLF-1 (*Deltaproteobacteria*)
*Desulfonatronum thiosulfatophilum ASO4-2 (Deltaproteobacteria)*
*Desulfonatronum zhilinae* Al915-01 (*Deltaproteobacteria*)

*Desulfopila aestuarii* DSM 18488 (*Deltaproteobacteria*)
*Desulfosarcina cetonica* JCM 12296 (*Deltaproteobacteria*)
*Desulfospira joergensenii* DSM 10085 (*Deltaproteobacteria*)
*Desulfosporosinus acididurans* M1 (*Firmicutes-Clostridia*)
*Desulfosporosinus acidophilus* SJ4 DSM 22704 (*Firmicutes-Clostridia*)
*Desulfosporosinus lacus* DSM 15449 (*Firmicutes-Clostridia*)
*Desulfosporosinus orientis* Singapore I DSM 765 (*Firmicutes-Clostridia*)
*Desulfosporosinus* sp. I2 (*Firmicutes-Clostridia*)
*Desulfosporosinus* sp. OT (*Firmicutes-Clostridia*)
*Desulfosporosinus* sp. Tol-M Ga0063340 (*Firmicutes-Clostridia*)
*Desulfosporosinus youngiae* JWYJL-B18 DSM 17734 (*Firmicutes-Clostridia*)
*Desulfotignum balticum* DSM 7044 (*Deltaproteobacteria*)
*Desulfotignum phosphitoxidans* FiPS-3 (*Deltaproteobacteria*)
*Desulfovibrio aespoeensis* Aspo-2 chromosome (*Deltaproteobacteria*)
*Desulfovibrio africanus* DSM 2603 2527068623 (*Deltaproteobacteria*)
*Desulfovibrio africanus* PCS 2520045431 (*Deltaproteobacteria*)
*Desulfovibrio africanus* Walvis Bay (*Deltaproteobacteria*)
*Desulfovibrio alkalitolerans* DSM 16529 (*Deltaproteobacteria*)
*Desulfovibrio bizertensis* DSM 18034 (*Deltaproteobacteria*)
*Desulfovibrio desulfuricans* ND132 (*Deltaproteobacteria*)
*Desulfovibrio halophilus* DSM 5663 (*Deltaproteobacteria*)
*Desulfovibrio inopinatus* DSM 10711 (*Deltaproteobacteria*)
*Desulfovibrio longus* DSM 6739 (*Deltaproteobacteria*)
*Desulfovibrio oxyclinae* DSM 11498 (*Deltaproteobacteria*)
*Desulfovibrio putealis* DSM 16056 (*Deltaproteobacteria*)
*Desulfovibrio* sp. isolate SP109 91617 (*Deltaproteobacteria*)
*Desulfovibrio* sp. J2 (*Deltaproteobacteria*)
*Desulfovibrio* sp. L21-Syr-AB (*Deltaproteobacteria*)
*Desulfovibrio* sp. X2 (*Deltaproteobacteria*)
*Desulfuromonadales* bacterium GWC2_61_20 (*Deltaproteobacteria*)
*Desulfuromonas* sp. DDH964 (*Deltaproteobacteria*)
*Desulfuromonas* sp. WTL (*Deltaproteobacteria*)
*Dethiobacter alkaliphilus* AHT 1 (*Firmicutes-Clostridia*)
*Elusimicrobia* bacterium GWA2_64_40 (*Elusimicrobia*)
*Elusimicrobia* bacterium GWA2_69_24 (*Elusimicrobia*)
*Elusimicrobia* bacterium RIFOXYA2_FULL_40_6 (*Elusimicrobia*)
*Elusimicrobia* bacterium RIFOXYA2_FULL_47_53 (*Elusimicrobia*)
*Elusimicrobia* bacterium RIFOXYA12_FULL_49_49 (*Elusimicrobia*)
*Elusimicrobia* bacterium RIFOXYB1_FULL_48_9 (*Elusimicrobia*)
*Elusimicrobia* bacterium RIFOXYB2_FULL_46_23 (*Elusimicrobia*)

*Elusimicrobia* bacterium RIFOXYB2_FULL_48_7 (*Elusimicrobia*)*
*Elusimicrobia* bacterium RIFOXYB12_FULL_50_12 (*Elusimicrobia*)
*Ethanoligenens harbinense* YUAN-3 chromosome (*Firmicutes-Clostridia*)
*Gammaproteobacteria* bacterium isolate NP949 (*Gammaproteobacteria*)*
*Geoalkalibacter ferrihydriticus* DSM 17813 (*Deltaproteobacteria*)
*Geobacter anodireducens* SD-1 (*Deltaproteobacteria*)
*Geobacter argillaceus* ATCC BAA-1139 (*Deltaproteobacteria*)
*Geobacter bemidjiensis* Bem (*Deltaproteobacteria*)
*Geobacter bremensis* R1 (*Deltaproteobacteria*)
*Geobacter daltonii* (*Deltaproteobacteria*)
*Geobacter metallireducens* GS-15 (*Deltaproteobacteria*)
*Geobacter metallireducens* RCH3 (*Deltaproteobacteria*)
*Geobacter pickeringii* G13 DSM 17153 (*Deltaproteobacteria*)
*Geobacter soli* GSS01 (*Deltaproteobacteria*)
*Geobacter* sp. M18 (*Deltaproteobacteria*)
*Geobacter* sp. M21 (*Deltaproteobacteria*)
*Geobacter* sp. OR-1 (Deltaproteobacteria)
*Geobacter sulfurreducens* AM-1 (*Deltaproteobacteria*)
*Geobacter sulfurreducens* KN400 (*Deltaproteobacteria*)
*Geobacter sulfurreducens* PCA (*Deltaproteobacteria*)
*Geobacter uraniumreducens* Rf4 (*Deltaproteobacteria*)
*Geopsychrobacter electrodiphilus* DSM 16401 (*Deltaproteobacteria*)
*Ignavibacteria* bacterium RIFOXYA2 FULL 35 10 (*Chlorobi*)*
*Kosmotoga pacifica* SLHLJ1 (*Thermotogae*)
*Lentisphaerae* bacterium GWF2_57_35 (*Lentisphaerae*)
*Lentisphaerae* bacterium RIFOXYA12_64_32 (*Lentisphaerae*)
*Lentisphaerae* bacterium RIFOXYA12_FULL_48_11 (*Lentisphaerae*)
*Lentisphaerae* bacterium RIFOXYB12_FULL_65_16 (*Lentisphaerae*)
*Lentisphaerae* bacterium RIFOXYC12_FULL_60_16 (*Lentisphaerae*)
*Marinilabilia salmonicolor* MSL42 (*Bacteroidetes*)
*Methanocella paludicola* SANAE (*Methanomicrobia*)
*Methanocella* sp. RC-I (*Methanomicrobia*)
*Methanococcoides methylutens* DSM 2657 (*Methanomicrobia*)
*Methanocorpusculum bavaricum* DSM 4179 (*Methanomicrobia*)
*Methanofollis liminatans* GKZPZ DSM 4140 (*Methanomicrobia*)
*Methanolobus profundi* Mob M (*Methanomicrobia*)
*Methanolobus psychrophilus* R15 (*Methanomicrobia*)
*Methanolobus tindarius* DSM 2278 (*Methanomicrobia*)
*Methanolobus vulcani* PL 12M (*Methanomicrobia*)
*Methanomassiliicoccales* archaeon RumEn M1 Ga0117923 (*Thermoplasmata*)

*Methanomassiliicoccus luminyensis* B10 (*Thermoplasmata*)
*Methanomethylovorans hollandica* DSM 15978 (*Methanomicrobia*)
*Methanoregula formicicum* SMSP (*Methanomicrobia*)
*Methanoregulaceae* archaeon JGI M3C4D3-001-G22 (*Methanomicrobia*)
*Methanospirillum hungatei* JF-1 (*Methanomicrobia*)
*Natronincola peptidivorans* DSM 18979 (*Firmicutes-Clostridia*)
*Nitrospina* AB-629-B06 (*Nitrospina*)*
*Nitrospina* SCGC AAA288-L16 (*Nitrospina*)*
*Nitrospira* bacterium SG8_3_2 (*Nitrospirae*)
*Nitrospira* bacterium SG8_3 (*Nitrospirae*)
*Nitrospira* bacterium SG8_35_1 (*Nitrospirae*)
*Nitrospirae* bacterium GWC2_56_14_2 (*Nitrospirae*)
*Nitrospirae* bacterium GWC2_56_14 (*Nitrospirae*)
*Nitrospirae* bacterium GWF2_44_13 (*Nitrospirae*)
*Nitrospirae* bacterium RBG_19FT_COMBO_55_12 (*Nitrospirae*)
*Nitrospirae* bacterium RIFOXYA2_FULL_44_9 (Nitrospirae)
*Nitrospirae* bacterium RIFOXYB2_FULL_43_5_partialB (*Nitrospirae*)
OMZ North Pacific GP0053648 LPjun08P16500mDRAFT (*Nitrospina*)*
OMZ North Pacific GP0053648 LPjun08P41300mDRAFT (*Nitrospina*)*
*Pelobacter seleniigenes* DSM 18267 (*Deltaproteobacteria*)
*Pelobacteraceae* bacterium GWC2 58 13 (*Deltaproteobacteria*)
*Phycisphaerae* bacterium SG8_4 (*Planctomycetes*)
*Pyrococcus furiosus* COM1 DSM 3638 (*Thermococci*)
*Smithella* sp. F21 (*Deltaproteobacteria*)
*Spirochaeta* sp. JC202 (*Spirochaetes*)
*Spirochaetes* bacterium GWB1_27_13 (*Spirochaetes*)
*Spirochaetes* bacterium GWB1_36_13 (*Spirochaetes*)
*Spirochaetes* bacterium GWB1_60_80 (*Spirochaetes*)
*Spirochaetes* bacterium GWB1_66_5 (*Spirochaetes*)
*Spirochaetes* bacterium GWE2_31_10 (*Spirochaetes*)
*Spirochaetes* bacterium GWF1_31_7 (*Spirochaetes*)
*Spirochaetes* bacterium GWF1_41_5 (*Spirochaetes*)
*Spirochaetes* bacterium GWF1_49_6 (*Spirochaetes*)
*Spirochaetes* bacterium GWF1_51_8 (*Spirochaetes*)
*Spirochaetes* bacterium GWF1_60_12 (*Spirochaetes*)
*Spirochaetes* bacterium RBG_16_49_21 (*Spirochaetes*)
*Spirochaetes* bacterium RBG_16_67_19 (*Spirochaetes*)
*Spirochaetes* bacterium RIFOXYB1_FULL_32_8 (*Spirochaetes*)
*Spirochaetes* bacterium RIFOXYC1_FULL_54_7 (*Spirochaetes*)
*Streptomyces* sp. CNQ-509 (*Actinobacteria*)

*Syntrophobacterales* bacterium GWC2_56_13 (*Deltaproteobacteria*)
*Syntrophobacterales* bacterium RBG_19FT_COMBO_59_10 (*Deltaproteobacteria*)*
*Syntrophobotulus glycolicus* DSM_8271 (*Firmicutes-Clostridia*)
*Syntrophorhabdus aromaticivorans* UI (*Deltaproteobacteria*)
*Syntrophus aciditrophicus* SB (*Deltaproteobacteria*)
*Syntrophus gentianae* DSM 8423 (*Deltaproteobacteria*)
*Syntrophus* sp. G20 Ga0063310 (*Deltaproteobacteria*)
*Theionarchaea archaeon* DG-70-1 (*Theionarchaea*)
*Thermococcus* sp. EP1 (*Thermococci*)
*Treponema* sp. GWA1_62_8 (*Spirochaetes*)
*Treponema* sp. GWB1_62_6 (*Spirochaetes*)
uncultured *Desulfarculaceae* 191 (*Deltaproteobacteria*)*

**Supplementary Table 2.** All forward and reverse primer oligonucleotide sequences from this study, including GC content (%), annealing temperature (Tm), whether the sequence has an exact match to reference library (Table S1) or amplicon library (Table S4) *hgcA* and *hgcB* sequences, and if the sequence is shared between reverse primers (shown in bold).

| **Sequence Name** | **Sequence 5' to 3'** | **% GC** | **Tm (°C)** | **Reference (Y/N)** | **Amplicon (Y/N)** | **Same As** |
|---|---|---|---|---|---|---|
| ORNL-HgcAB-uni-F | AAYGTCTGGTGYGCNGCVGG | 68.8 | 60.9 - 70.7 | Y | Y | |
| ORNL-HgcAB-uni-32R | CAGGCNCCGCAYTCSATRCA | 64.7 | 60.2 - 68.5 | Y | Y | |
| ORNL-HgcAB-uni-R | CABGCNCCRCAYTCCATRCA | 60 | 56.1 - 68.2 | Y | Y | |
| **ORNL-HgcAB-uni-F1** | AACGTCTGGTGCGCAGCCGG | 70 | 68.4 | N | Y | |
| **ORNL-HgcAB-uni-F2** | AACGTCTGGTGCGCAGCGGG | 70 | 68.4 | Y | Y | |
| **ORNL-HgcAB-uni-F3** | AACGTCTGGTGCGCAGCAGG | 65 | 66 | N | Y | |
| **ORNL-HgcAB-uni-F4** | AACGTCTGGTGCGCCGCCGG | 75 | 70.7 | Y | Y | |
| **ORNL-HgcAB-uni-F5** | AACGTCTGGTGCGCCGCGGG | 75 | 70.7 | Y | Y | |
| **ORNL-HgcAB-uni-F6** | AACGTCTGGTGCGCCGCAGG | 70 | 68.4 | Y | Y | |
| **ORNL-HgcAB-uni-F7** | AACGTCTGGTGCGCTGCCGG | 70 | 68.4 | Y | Y | |
| **ORNL-HgcAB-uni-F8** | AACGTCTGGTGCGCTGCGGG | 70 | 68.4 | Y | Y | |
| **ORNL-HgcAB-uni-F9** | AACGTCTGGTGCGCTGCAGG | 65 | 66 | N | Y | |
| **ORNL-HgcAB-uni-F10** | AACGTCTGGTGCGCGGCCGG | 75 | 70.7 | Y | Y | |
| **ORNL-HgcAB-uni-F11** | AACGTCTGGTGCGCGGCGGG | 75 | 70.7 | Y | Y | |
| **ORNL-HgcAB-uni-F12** | AACGTCTGGTGCGCGGCAGG | 70 | 68.4 | Y | Y | |
| **ORNL-HgcAB-uni-F13** | AACGTCTGGTGTGCAGCCGG | 65 | 65.7 | Y | Y | |
| **ORNL-HgcAB-uni-F14** | AACGTCTGGTGTGCAGCGGG | 65 | 65.7 | N | Y | |
| **ORNL-HgcAB-uni-F15** | AACGTCTGGTGTGCAGCAGG | 60 | 63.3 | N | Y | |
| **ORNL-HgcAB-uni-F16** | AACGTCTGGTGTGCCGCCGG | 70 | 68.1 | N | Y | |
| **ORNL-HgcAB-uni-F17** | AACGTCTGGTGTGCCGCGGG | 70 | 68.1 | N | Y | |
| **ORNL-HgcAB-uni-F18** | AACGTCTGGTGTGCCGCAGG | 65 | 65.7 | Y | Y | |
| **ORNL-HgcAB-uni-F19** | AACGTCTGGTGTGCTGCCGG | 65 | 65.7 | Y | Y | |
| **ORNL-HgcAB-uni-F20** | AACGTCTGGTGTGCTGCGGG | 65 | 65.7 | N | Y | |
| **ORNL-HgcAB-uni-F21** | AACGTCTGGTGTGCTGCAGG | 60 | 63.3 | N | Y | |
| **ORNL-HgcAB-uni-F22** | AACGTCTGGTGTGCGGCCGG | 70 | 68.1 | Y | Y | |
| **ORNL-HgcAB-uni-F23** | AACGTCTGGTGTGCGGCGGG | 70 | 68.1 | Y | Y | |
| **ORNL-HgcAB-uni-F24** | AACGTCTGGTGTGCGGCAGG | 65 | 65.7 | N | Y | |
| **ORNL-HgcAB-uni-F25** | AATGTCTGGTGCGCAGCCGG | 65 | 66.1 | Y | Y | |
| **ORNL-HgcAB-uni-F26** | AATGTCTGGTGCGCAGCGGG | 65 | 66.1 | Y | Y | |
| **ORNL-HgcAB-uni-F27** | AATGTCTGGTGCGCAGCAGG | 60 | 63.7 | Y | Y | |
| **ORNL-HgcAB-uni-F28** | AATGTCTGGTGCGCCGCCGG | 70 | 68.5 | Y | Y | |
| **ORNL-HgcAB-uni-F29** | AATGTCTGGTGCGCCGCGGG | 70 | 68.5 | Y | Y | |

| | | | | | | |
|---|---|---|---|---|---|---|
| **ORNL-HgcAB-uni-F30** | AATGTCTGGTGCGCCGCAGG | 65 | 66.1 | Y | Y | |
| **ORNL-HgcAB-uni-F31** | AATGTCTGGTGCGCTGCCGG | 65 | 66.1 | Y | Y | |
| **ORNL-HgcAB-uni-F32** | AATGTCTGGTGCGCTGCGGG | 65 | 66.1 | Y | Y | |
| **ORNL-HgcAB-uni-F33** | AATGTCTGGTGCGCTGCAGG | 60 | 63.7 | Y | Y | |
| **ORNL-HgcAB-uni-F34** | AATGTCTGGTGCGCGGCCGG | 70 | 68.5 | Y | Y | |
| **ORNL-HgcAB-uni-F35** | AATGTCTGGTGCGCGGCGGG | 70 | 68.5 | Y | Y | |
| **ORNL-HgcAB-uni-F36** | AATGTCTGGTGCGCGGCAGG | 65 | 66.1 | Y | Y | |
| **ORNL-HgcAB-uni-F37** | AATGTCTGGTGTGCAGCCGG | 60 | 63.4 | Y | Y | |
| **ORNL-HgcAB-uni-F38** | AATGTCTGGTGTGCAGCGGG | 60 | 63.4 | Y | Y | |
| **ORNL-HgcAB-uni-F39** | AATGTCTGGTGTGCAGCAGG | 55 | 60.9 | Y | Y | |
| **ORNL-HgcAB-uni-F40** | AATGTCTGGTGTGCCGCCGG | 65 | 65.8 | Y | Y | |
| **ORNL-HgcAB-uni-F41** | AATGTCTGGTGTGCCGCGGG | 65 | 65.8 | Y | Y | |
| **ORNL-HgcAB-uni-F42** | AATGTCTGGTGTGCCGCAGG | 60 | 63.4 | Y | Y | |
| **ORNL-HgcAB-uni-F43** | AATGTCTGGTGTGCTGCCGG | 60 | 63.4 | Y | Y | |
| **ORNL-HgcAB-uni-F44** | AATGTCTGGTGTGCTGCGGG | 60 | 63.4 | Y | Y | |
| **ORNL-HgcAB-uni-F45** | AATGTCTGGTGTGCTGCAGG | 55 | 60.9 | N | Y | |
| **ORNL-HgcAB-uni-F46** | AATGTCTGGTGTGCGGCCGG | 65 | 65.8 | Y | Y | |
| **ORNL-HgcAB-uni-F47** | AATGTCTGGTGTGCGGCGGG | 65 | 65.8 | Y | Y | |
| **ORNL-HgcAB-uni-F48** | AATGTCTGGTGTGCGGCAGG | 60 | 63.4 | Y | Y | |
| **ORNL-HgcAB-uni-32R-1** | CAGGCACCGCATTCCATACA | 55 | 60.7 | Y | Y | ORNL-HgcAB-uni-R-56 |
| **ORNL-HgcAB-uni-32R-2** | CAGGCACCGCATTCCATGCA | 60 | 64.1 | Y | Y | ORNL-HgcAB-uni-R-8 |
| ORNL-HgcAB-uni-32R-3 | CAGGCACCGCATTCGATACA | 55 | 60.8 | N | Y | |
| ORNL-HgcAB-uni-32R-4 | CAGGCACCGCATTCGATGCA | 60 | 64.1 | Y | Y | |
| **ORNL-HgcAB-uni-32R-5** | CAGGCACCGCACTCCATACA | 60 | 62.5 | Y | Y | ORNL-HgcAB-uni-R-80 |
| **ORNL-HgcAB-uni-32R-6** | CAGGCACCGCACTCCATGCA | 65 | 65.8 | N | Y | ORNL-hgcAB-uni-R-32 |
| ORNL-HgcAB-uni-32R-7 | CAGGCACCGCACTCGATACA | 60 | 62.6 | N | Y | |
| ORNL-HgcAB-uni-32R-8 | CAGGCACCGCACTCGATGCA | 65 | 65.8 | N | Y | |
| **ORNL-HgcAB-uni-32R-9** | CAGGCTCCGCATTCCATACA | 55 | 60.2 | Y | Y | ORNL-HgcAB-uni-R-59 |
| **ORNL-HgcAB-uni-32R-10** | CAGGCTCCGCATTCCATGCA | 60 | 63.5 | Y | Y | ORNL-HgcAB-uni-R-11 |
| ORNL-HgcAB-uni-32R-11 | CAGGCTCCGCATTCGATACA | 55 | 60.2 | N | Y | |
| ORNL-HgcAB-uni-32R-12 | CAGGCTCCGCATTCGATGCA | 60 | 63.5 | N | Y | |
| **ORNL-HgcAB-uni-32R-13** | CAGGCTCCGCACTCCATACA | 60 | 62 | N | Y | ORNL-HgcAB-uni-R-83 |
| **ORNL-HgcAB-uni-32R-14** | CAGGCTCCGCACTCCATGCA | 65 | 65.3 | Y | Y | ORNL-HgcAB-uni-R-35 |
| ORNL-HgcAB-uni-32R-15 | CAGGCTCCGCACTCGATACA | 60 | 62 | N | Y | |
| ORNL-HgcAB-uni-32R-16 | CAGGCTCCGCACTCGATGCA | 65 | 65.3 | N | Y | |
| **ORNL-HgcAB-uni-32R-17** | CAGGCCCCGCATTCCATACA | 60 | 62.6 | Y | Y | ORNL-HgcAB-uni-R-53 |
| **ORNL-HgcAB-uni-32R-18** | CAGGCCCCGCATTCCATGCA | 65 | 66 | Y | Y | ORNL-HgcAB-uni-R-5 |
| ORNL-HgcAB-uni-32R-19 | CAGGCCCCGCATTCGATACA | 60 | 62.7 | N | Y | |
| ORNL-HgcAB-uni-32R-20 | CAGGCCCCGCATTCGATGCA | 65 | 65.9 | Y | Y | |

| | | | | | | |
|---|---|---|---|---|---|---|
| **ORNL-HgcAB-uni-32R-21** | CAGGCCCCGCACTCCATACA | 65 | 64.4 | Y | Y | ORNL-HgcAB-uni-R-77 |
| **ORNL-HgcAB-uni-32R-22** | CAGGCCCCGCACTCCATGCA | 70 | 67.7 | Y | Y | ORNL-HgcAB-uni-R-29 |
| ORNL-HgcAB-uni-32R-23 | CAGGCCCCGCACTCGATACA | 65 | 64.4 | N | Y | |
| ORNL-HgcAB-uni-32R-24 | CAGGCCCCGCACTCGATGCA | 70 | 67.7 | N | Y | |
| **ORNL-HgcAB-uni-32R-25** | CAGGCGCCGCATTCCATACA | 60 | 63.5 | Y | Y | ORNL-HgcAB-uni-R-50 |
| **ORNL-HgcAB-uni-32R-26** | CAGGCGCCGCATTCCATGCA | 65 | 66.8 | Y | Y | ORNL-HgcAB-uni-R-2 |
| ORNL-HgcAB-uni-32R-27 | CAGGCGCCGCATTCGATACA | 60 | 63.6 | Y | Y | |
| ORNL-HgcAB-uni-32R-28 | CAGGCGCCGCATTCGATGCA | 65 | 66.8 | Y | Y | |
| **ORNL-HgcAB-uni-32R-29** | CAGGCGCCGCACTCCATACA | 65 | 65.3 | Y | Y | ORNL-HgcAB-uni-R-74 |
| **ORNL-HgcAB-uni-32R-30** | CAGGCGCCGCACTCCATGCA | 70 | 68.5 | Y | Y | ORNL-HgcAB-uni-R-26 |
| ORNL-HgcAB-uni-32R-31 | CAGGCGCCGCACTCGATACA | 65 | 65.3 | N | Y | |
| ORNL-HgcAB-uni-32R-32 | CAGGCGCCGCACTCGATGCA | 70 | 68.5 | N | Y | |
| ORNL-HgcAB-uni-R-1 | CATGCTCCGCACTCCATGCA | 60 | 63.5 | N | Y | |
| **ORNL-HgcAB-uni-R-2** | CAGGCTCCGCACTCCATGCA | 65 | 65.3 | N | Y | ORNL-HgcAB-uni-32R-26 |
| ORNL-HgcAB-uni-R-3 | CACGCTCCGCACTCCATGCA | 65 | 65.8 | N | Y | |
| ORNL-HgcAB-uni-R-4 | CATGCGCCGCACTCCATGCA | 65 | 66.8 | Y | Y | |
| **ORNL-HgcAB-uni-R-5** | CAGGCGCCGCACTCCATGCA | 70 | 68.5 | Y | Y | ORNL-HgcAB-uni-32R-18 |
| ORNL-HgcAB-uni-R-6 | CACGCGCCGCACTCCATGCA | 70 | 69 | Y | Y | |
| ORNL-HgcAB-uni-R-7 | CATGCACCGCACTCCATGCA | 60 | 64 | N | Y | |
| **ORNL-HgcAB-uni-R-8** | CAGGCACCGCACTCCATGCA | 65 | 65.8 | N | Y | ORNL-HgcAB-uni-32R-26 |
| ORNL-HgcAB-uni-R-9 | CACGCACCGCACTCCATGCA | 65 | 66.3 | N | Y | |
| ORNL-HgcAB-uni-R-10 | CATGCCCCGCACTCCATGCA | 65 | 65.9 | Y | Y | |
| **ORNL-HgcAB-uni-R-11** | CAGGCCCCGCACTCCATGCA | 70 | 67.7 | Y | Y | ORNL-HgcAB-uni-32R-10 |
| ORNL-HgcAB-uni-R-12 | CACGCCCCGCACTCCATGCA | 70 | 68.2 | Y | Y | |
| ORNL-HgcAB-uni-R-13 | CATGCTCCACACTCCATGCA | 55 | 60.7 | N | Y | |
| ORNL-HgcAB-uni-R-14 | CAGGCTCCACACTCCATGCA | 60 | 62.5 | Y | Y | |
| ORNL-HgcAB-uni-R-15 | CACGCTCCACACTCCATGCA | 60 | 63.1 | N | Y | |
| ORNL-HgcAB-uni-R-16 | CATGCGCCACACTCCATGCA | 60 | 64 | N | Y | |
| ORNL-HgcAB-uni-R-17 | CAGGCGCCACACTCCATGCA | 65 | 65.8 | N | Y | |
| ORNL-HgcAB-uni-R-18 | CACGCGCCACACTCCATGCA | 65 | 66.3 | N | Y | |
| ORNL-HgcAB-uni-R-19 | CATGCACCACACTCCATGCA | 55 | 61.2 | Y | Y | |
| ORNL-HgcAB-uni-R-20 | CAGGCACCACACTCCATGCA | 60 | 63 | N | Y | |
| ORNL-HgcAB-uni-R-21 | CACGCACCACACTCCATGCA | 60 | 63.6 | N | Y | |
| ORNL-HgcAB-uni-R-22 | CATGCCCCACACTCCATGCA | 60 | 63.1 | N | Y | |
| ORNL-HgcAB-uni-R-23 | CAGGCCCCACACTCCATGCA | 65 | 64.9 | Y | Y | |
| ORNL-HgcAB-uni-R-24 | CACGCCCCACACTCCATGCA | 65 | 65.5 | N | Y | |
| ORNL-HgcAB-uni-R-25 | CATGCTCCGCATTCCATGCA | 55 | 61.7 | Y | Y | |
| **ORNL-HgcAB-uni-R-26** | CAGGCTCCGCATTCCATGCA | 60 | 63.5 | Y | Y | ORNL-HgcAB-uni-32R-30 |
| ORNL-HgcAB-uni-R-27 | CACGCTCCGCATTCCATGCA | 60 | 64.1 | N | Y | |

| | | | | | | |
|---|---|---|---|---|---|---|
| ORNL-HgcAB-uni-R-28 | CATGCGCCGCATTCCATGCA | 60 | 65 | Y | Y | |
| **ORNL-HgcAB-uni-R-29** | CAGGCGCCGCATTCCATGCA | 65 | 66.8 | Y | Y | ORNL-HgcAB-uni-32R-22 |
| ORNL-HgcAB-uni-R-30 | CACGCGCCGCATTCCATGCA | 65 | 67.3 | N | Y | |
| ORNL-HgcAB-uni-R-31 | CATGCACCGCATTCCATGCA | 55 | 62.3 | Y | Y | |
| **ORNL-HgcAB-uni-R-32** | CAGGCACCGCATTCCATGCA | 60 | 64.1 | Y | Y | ORNL-HgcAB-uni-32R-6 |
| ORNL-HgcAB-uni-R-33 | CACGCACCGCATTCCATGCA | 60 | 64.6 | Y | Y | |
| ORNL-HgcAB-uni-R-34 | CATGCCCCGCATTCCATGCA | 60 | 64.2 | Y | Y | |
| **ORNL-HgcAB-uni-R-35** | CAGGCCCCGCATTCCATGCA | 65 | 66 | Y | Y | ORNL-HgcAB-uni-32R-14 |
| ORNL-HgcAB-uni-R-36 | CACGCCCCGCATTCCATGCA | 65 | 66.5 | N | Y | |
| ORNL-HgcAB-uni-R-37 | CATGCTCCACATTCCATGCA | 50 | 58.9 | Y | Y | |
| ORNL-HgcAB-uni-R-38 | CAGGCTCCACATTCCATGCA | 55 | 60.7 | N | Y | |
| ORNL-HgcAB-uni-R-39 | CACGCTCCACATTCCATGCA | 55 | 61.3 | N | Y | |
| ORNL-HgcAB-uni-R-40 | CATGCGCCACATTCCATGCA | 55 | 62.3 | N | Y | |
| ORNL-HgcAB-uni-R-41 | CAGGCGCCACATTCCATGCA | 60 | 64.1 | Y | Y | |
| ORNL-HgcAB-uni-R-42 | CACGCGCCACATTCCATGCA | 60 | 64.6 | Y | Y | |
| ORNL-HgcAB-uni-R-43 | CATGCACCACATTCCATGCA | 50 | 59.5 | N | Y | |
| ORNL-HgcAB-uni-R-44 | CAGGCACCACATTCCATGCA | 55 | 61.3 | N | Y | |
| ORNL-HgcAB-uni-R-45 | CACGCACCACATTCCATGCA | 55 | 61.9 | N | Y | |
| ORNL-HgcAB-uni-R-46 | CATGCCCCACATTCCATGCA | 55 | 61.3 | N | Y | |
| ORNL-HgcAB-uni-R-47 | CAGGCCCCACATTCCATGCA | 60 | 63.1 | N | Y | |
| ORNL-HgcAB-uni-R-48 | CACGCCCCACATTCCATGCA | 60 | 63.7 | N | Y | |
| ORNL-HgcAB-uni-R-49 | CATGCTCCGCACTCCATACA | 55 | 60.2 | N | Y | |
| **ORNL-HgcAB-uni-R-50** | CAGGCTCCGCACTCCATACA | 60 | 62 | N | Y | ORNL-HgcAB-uni-32R-25 |
| ORNL-HgcAB-uni-R-51 | CACGCTCCGCACTCCATACA | 60 | 62.6 | N | Y | |
| ORNL-HgcAB-uni-R-52 | CATGCGCCGCACTCCATACA | 60 | 63.5 | N | Y | |
| **ORNL-HgcAB-uni-R-53** | CAGGCGCCGCACTCCATACA | 65 | 65.3 | Y | Y | ORNL-HgcAB-uni-32R-17 |
| ORNL-HgcAB-uni-R-54 | CACGCGCCGCACTCCATACA | 65 | 65.8 | Y | Y | |
| ORNL-HgcAB-uni-R-55 | CATGCACCGCACTCCATACA | 55 | 60.7 | N | Y | |
| **ORNL-HgcAB-uni-R-56** | CAGGCACCGCACTCCATACA | 60 | 62.5 | Y | Y | ORNL-HgcAB-uni-32R-1 |
| ORNL-HgcAB-uni-R-57 | CACGCACCGCACTCCATACA | 60 | 63.1 | Y | Y | |
| ORNL-HgcAB-uni-R-58 | CATGCCCCGCACTCCATACA | 60 | 62.6 | N | Y | |
| **ORNL-HgcAB-uni-R-59** | CAGGCCCCGCACTCCATACA | 65 | 64.4 | Y | Y | ORNL-HgcAB-uni-32R-9 |
| ORNL-HgcAB-uni-R-60 | CACGCCCCGCACTCCATACA | 65 | 65 | N | Y | |
| ORNL-HgcAB-uni-R-61 | CATGCTCCACACTCCATACA | 50 | 57.3 | Y | Y | |
| ORNL-HgcAB-uni-R-62 | CAGGCTCCACACTCCATACA | 55 | 59.1 | Y | Y | |
| ORNL-HgcAB-uni-R-63 | CACGCTCCACACTCCATACA | 55 | 59.8 | N | Y | |
| ORNL-HgcAB-uni-R-64 | CATGCGCCACACTCCATACA | 55 | 60.7 | N | Y | |
| ORNL-HgcAB-uni-R-65 | CAGGCGCCACACTCCATACA | 60 | 62.5 | N | Y | |
| ORNL-HgcAB-uni-R-66 | CACGCGCCACACTCCATACA | 60 | 63.1 | N | Y | |

| | | | | | | |
|---|---|---|---|---|---|---|
| ORNL-HgcAB-uni-R-67 | CATGCACCACACTCCATACA | 50 | 57.9 | Y | Y | |
| ORNL-HgcAB-uni-R-68 | CAGGCACCACACTCCATACA | 55 | 59.7 | N | Y | |
| ORNL-HgcAB-uni-R-69 | CACGCACCACACTCCATACA | 55 | 60.3 | N | Y | |
| ORNL-HgcAB-uni-R-70 | CATGCCCCACACTCCATACA | 55 | 59.7 | Y | Y | |
| ORNL-HgcAB-uni-R-71 | CAGGCCCCACACTCCATACA | 60 | 61.6 | Y | Y | |
| ORNL-HgcAB-uni-R-72 | CACGCCCCACACTCCATACA | 60 | 62.2 | N | Y | |
| ORNL-HgcAB-uni-R-73 | CATGCTCCGCATTCCATACA | 50 | 58.4 | Y | Y | |
| **ORNL-HgcAB-uni-R-74** | CAGGCTCCGCATTCCATACA | 55 | 60.2 | Y | Y | ORNL-HgcAB-uni-32R-29 |
| ORNL-HgcAB-uni-R-75 | CACGCTCCGCATTCCATACA | 55 | 60.8 | N | Y | |
| ORNL-HgcAB-uni-R-76 | CATGCGCCGCATTCCATACA | 55 | 61.8 | N | Y | |
| **ORNL-HgcAB-uni-R-77** | CAGGCGCCGCATTCCATACA | 60 | 63.5 | Y | Y | ORNL-HgcAB-uni-32R-21 |
| ORNL-HgcAB-uni-R-78 | CACGCGCCGCATTCCATACA | 60 | 64.1 | Y | Y | |
| ORNL-HgcAB-uni-R-79 | CATGCACCGCATTCCATACA | 50 | 59 | Y | Y | |
| **ORNL-HgcAB-uni-R-80** | CAGGCACCGCATTCCATACA | 55 | 60.7 | Y | Y | ORNL-HgcAB-uni-32R-5 |
| ORNL-HgcAB-uni-R-81 | CACGCACCGCATTCCATACA | 55 | 61.4 | Y | Y | |
| ORNL-HgcAB-uni-R-82 | CATGCCCCGCATTCCATACA | 55 | 60.8 | N | Y | |
| **ORNL-HgcAB-uni-R-83** | CAGGCCCCGCATTCCATACA | 60 | 62.6 | Y | Y | ORNL-HgcAB-uni-32R-13 |
| ORNL-HgcAB-uni-R-84 | CACGCCCCGCATTCCATACA | 60 | 63.2 | Y | Y | |
| ORNL-HgcAB-uni-R-85 | CATGCTCCACATTCCATACA | 45 | 55.5 | Y | Y | |
| ORNL-HgcAB-uni-R-86 | CAGGCTCCACATTCCATACA | 50 | 57.3 | Y | Y | |
| ORNL-HgcAB-uni-R-87 | CACGCTCCACATTCCATACA | 50 | 58 | N | Y | |
| ORNL-HgcAB-uni-R-88 | CATGCGCCACATTCCATACA | 50 | 59 | Y | Y | |
| ORNL-HgcAB-uni-R-89 | CAGGCGCCACATTCCATACA | 55 | 60.7 | Y | Y | |
| ORNL-HgcAB-uni-R-90 | CACGCGCCACATTCCATACA | 55 | 61.4 | N | Y | |
| ORNL-HgcAB-uni-R-91 | CATGCACCACATTCCATACA | 45 | 56.1 | Y | Y | |
| ORNL-HgcAB-uni-R-92 | CAGGCACCACATTCCATACA | 50 | 57.9 | N | Y | |
| ORNL-HgcAB-uni-R-93 | CACGCACCACATTCCATACA | 50 | 58.6 | N | Y | |
| ORNL-HgcAB-uni-R-94 | CATGCCCCACATTCCATACA | 50 | 57.9 | Y | Y | |
| ORNL-HgcAB-uni-R-95 | CAGGCCCCACATTCCATACA | 55 | 59.7 | N | Y | |
| ORNL-HgcAB-uni-R-96 | CACGCCCCACATTCCATACA | 55 | 60.4 | N | Y | |

**Supplementary Table 3.** Environmental samples used for *hgcAB* clone libraries. Unless otherwise designated, *hgcAB* clone sequences were amplified with primer set ORNL-HgcAB-uni-F/ORNL-HgcAB-uni-32R. The number of clones are those that passed quality filtering criteria and were classified as *hgcAB*. The environmental clone hgcA sequences from this study are publicly available under the NCBI GenBank accession numbers MT122211 - MT122744.

| Clone Sample IDs | Sample Site | Environment Type | Sample collection (depth, cm) | Sample collection (date) | Location | Clones (#) |
|---|---|---|---|---|---|---|
| Spruce and Peatland Responses Under Changing Environments (SPRUCE) (Iversen et al., 2014) | | Boreal spruce and peatland sediments | From surface (0 cm) to 255 cm depth | | USDA Forest service Marcell Experiment Forest (Grand Rapids, MN) | 232 |
| 4 | 2012-10-T-Hol | Treed Hollow | -30 | 2012-08-14 | S1-Bog, Plot #10 | 2 |
| 5 | 2012-10-T-Hol | Treed Hollow | -40 | 2012-08-14 | S1-Bog, Plot #10 | 5 |
| 6 | 2012-10-T-Hol | Treed Hollow | -50 | 2012-08-14 | S1-Bog, Plot #10 | 5 |
| 7 | 2012-10-T-Hol | Treed Hollow | -60 | 2012-08-14 | S1-Bog, Plot #10 | 5 |
| 8 | 2012-10-T-Hol | Treed Hollow | -70 | 2012-08-14 | S1-Bog, Plot #10 | 3 |
| 9 | 2012-10-T-Hol | Treed Hollow | -80 | 2012-08-14 | S1-Bog, Plot #10 | 5 |
| 10 | 2012-10-T-Hol | Treed Hollow | -90 | 2012-08-14 | S1-Bog, Plot #10 | 2 |
| 11 | 2012-10-T-Hol | Treed Hollow | -100 | 2012-08-14 | S1-Bog, Plot #10 | 5 |
| 12 | 2012-10-T-Hol | Treed Hollow | -125 | 2012-08-14 | S1-Bog, Plot #10 | 3 |
| 13 | 2012-10-T-Hol | Treed Hollow | -150 | 2012-08-14 | S1-Bog, Plot #10 | 4 |
| 15 | 2012-10-T-Hol | Treed Hollow | -200 | 2012-08-14 | S1-Bog, Plot #10 | 5 |
| 16 | 2012-10-T-Hol | Treed Hollow | -225 | 2012-08-14 | S1-Bog, Plot #10 | 7 |
| 17 | 2012-10-T-Hol | Treed Hollow | -255 | 2012-08-14 | S1-Bog, Plot #10 | 5 |
| 20 | 2012-10-T-Hum | Treed Hummock | +10 | 2012-08-14 | S1-Bog, Plot #10 | 3 |
| 21 | 2012-10-T-Hum | Treed Hummock | 0 | 2012-08-14 | S1-Bog, Plot #10 | 2 |
| 22 | 2012-10-T-Hum | Treed Hummock | -10 | 2012-08-14 | S1-Bog, Plot #10 | 5 |
| 23 | 2012-10-T-Hum | Treed Hummock | -20 | 2012-08-14 | S1-Bog, Plot #10 | 4 |
| 24 | 2012-10-T-Hum | Treed Hummock | -30 | 2012-08-14 | S1-Bog, Plot #10 | 4 |
| 25 | 2012-10-T-Hum | Treed Hummock | -40 | 2012-08-14 | S1-Bog, Plot #10 | 5 |
| 26 | 2012-10-T-Hum | Treed Hummock | -50 | 2012-08-14 | S1-Bog, Plot #10 | 4 |
| 27 | 2012-10-T-Hum | Treed Hummock | -60 | 2012-08-14 | S1-Bog, Plot #10 | 5 |
| 28 | 2012-10-T-Hum | Treed Hummock | -70 | 2012-08-14 | S1-Bog, Plot #10 | 6 |
| 29 | 2012-10-T-Hum | Treed Hummock | -80 | 2012-08-14 | S1-Bog, Plot #10 | 5 |
| 30 | 2012-10-T-Hum | Treed Hummock | -90 | 2012-08-14 | S1-Bog, Plot #10 | 5 |
| 31 | 2012-10-T-Hum | Treed Hummock | -100 | 2012-08-14 | S1-Bog, Plot #10 | 4 |
| 32 | 2012-10-T-Hum | Treed Hummock | -125 | 2012-08-14 | S1-Bog, Plot #10 | 4 |
| 33 | 2012-10-T-Hum | Treed Hummock | -150 | 2012-08-14 | S1-Bog, Plot #10 | 4 |
| 34 | 2012-10-T-Hum | Treed Hummock | -175 | 2012-08-14 | S1-Bog, Plot #10 | 5 |
| 39 | 2012-6-T-Hol | Treed Hollow | -40 | 2012-08-14 | S1-Bog, Plot #6 | 2 |
| 40 | 2012-6-T-Hol | Treed Hollow | -50 | 2012-08-14 | S1-Bog, Plot #6 | 4 |
| 41 | 2012-6-T-Hol | Treed Hollow | -60 | 2012-08-14 | S1-Bog, Plot #6 | 4 |
| 42 | 2012-6-T-Hol | Treed Hollow | -70 | 2012-08-14 | S1-Bog, Plot #6 | 5 |
| 43 | 2012-6-T-Hol | Treed Hollow | -80 | 2012-08-14 | S1-Bog, Plot #6 | 4 |
| 44, 44H, 44L | 2012-6-T-Hol | Treed Hollow | -90 | 2012-08-14 | S1-Bog, Plot #6 | 10 |
| 45H | 2012-6-T-Hol | Treed Hollow | -100 | 2012-08-14 | S1-Bog, Plot #6 | 4 |
| 46, 46-4, 46-7 | 2012-6-T-Hol | Treed Hollow | -125 | 2012-08-14 | S1-Bog, Plot #6 | 5 |

| | | | | | | |
|---|---|---|---|---|---|---|
| 47, 47-2, 47H, 47L | 2012-6-T-Hol | Treed Hollow | -150 | 2012-08-14 | S1-Bog, Plot #6 | 12 |
| 48-5, 48-17 | 2012-6-T-Hol | Treed Hollow | -175 | 2012-08-14 | S1-Bog, Plot #6 | 6 |
| 49 | 2012-6-T-Hol | Treed Hollow | -200 | 2012-08-14 | S1-Bog, Plot #6 | 5 |
| 53, 53-17 | 2012-6-T-Hum | Treed Hummock | -10 | 2012-08-14 | S1-Bog, Plot #6 | 5 |
| 54, 54H | 2012-6-T-Hum | Treed Hummock | -30 | 2012-08-14 | S1-Bog, Plot #6 | 5 |
| 55 | 2012-6-T-Hum | Treed Hummock | -40 | 2012-08-14 | S1-Bog, Plot #6 | 6 |
| 56, 56H | 2012-6-T-Hum | Treed Hummock | -50 | 2012-08-14 | S1-Bog, Plot #6 | 3 |
| 57,57H | 2012-6-T-Hum | Treed Hummock | -60 | 2012-08-14 | S1-Bog, Plot #6 | 4 |
| 58, 58H | 2012-6-T-Hum | Treed Hummock | -70 | 2012-08-14 | S1-Bog, Plot #6 | 6 |
| 59,59H | 2012-6-T-Hum | Treed Hummock | -80 | 2012-08-14 | S1-Bog, Plot #6 | 4 |
| 60 | 2012-6-T-Hum | Treed Hummock | -90 | 2012-08-14 | S1-Bog, Plot #6 | 5 |
| 61 | 2012-6-T-Hum | Treed Hummock | -100 | 2012-08-14 | S1-Bog, Plot #6 | 5 |
| 62 | 2012-6-T-Hum | Treed Hummock | -125 | 2012-08-14 | S1-Bog Plot #6 | 3 |
| 63 | 2012-6-T-Hum | Treed Hummock | -150 | 2012-08-14 | S1-Bog, Plot #6 | 4 |
| SPRUCE1 (OP1+, NP1) | Equimolar mixture of gDNA from SPRUCE samples 16, 34, 40, and 57 | | | | | 58 (OP1+), 65 (NP1) |
| Sandy Creek (BRL) (Ndu et al., 2018) | Freshwater stream sediment | Surface grab sample | 2017-03-03 | Durham, NC | | 19 |
| New Horizon sediment (HGC5, OP3+, NP3) | Freshwater sediments | Surface grab sample (0 cm) | 2017-09-21 (HGC5); 2018-01-19 (NP3, OP3) | East Fork Poplar Creek, Oak Ridge, TN | | 3 (HGC 5), 42 (OP3+), 61 (NP3) |
| New Horizon periphyton (HGC2, Y117) (Olsen et al., 2016) | Freshwater periphyton biomass | Collected from rocks in stream | 2016-08-23 (Y117) And 2017-09-21(HGC2) | East Fork Poplar Creek, Oak Ridge, TN | | 1 (HGC 2), 14 (Y117) |
| Mesohaline tidal salt marsh sediment (GC) (Mitchell and Gilmour, 2008) | Marsh sediment | Top 15 cm | 2015-09-30 | Rhode River, Edgewater, MD | | 12 |
| Yanwuping Rice Paddy (D118) (Vishnivetskaya et al., 2018) | Rice paddy soils | 5 cm depth | 2011-07 | Guizhou, China | | 38 |

+*hgcAB* clone sequences from primer set ORNL-HgcAB-uni-F/ORNL-HgcAB-uni-R

**Supplementary Table 4.** Miseq *hgcAB* amplicon sequencing libraries for environmental, mock, and spiked communities. The environmental and mock community *hgcAB* amplicon raw sequence files are publicly available at the NCBI SRA accession (PRJNA608965).

| Sample type | Sample environment | Sample ID | Amplicon library | Replicates + (#) | Reverse primer | PCR cycles | Amplicons (#) |
|---|---|---|---|---|---|---|---|
| Environmental | Tidal salt marsh sediment (1064) | 1064 | 1064_Alt35 | 1 | ORNL-HgcAB-uni-32R | x35 | 58374 |
| | | | 1064_Uni35 | 1 | ORNL-HgcAB-uni-R | x35 | 41218 |
| | New Horizon (NH) surface sediment | NH1 | NH1_Alt35 | 1 | ORNL-HgcAB-uni-32R | x35 | 43828 |
| | | | NH1_Uni35 | 1 | ORNL-HgcAB-uni-R | x35 | 62533 |
| | | NH2 | NH2_Alt35 | 1 | ORNL-HgcAB-uni-32R | x35 | 63434 |
| | | | NH2_Uni35 | 1 | ORNL-HgcAB-uni-R | x35 | 69702 |
| | | NH3 | NH3_Alt35 | 1 | ORNL-HgcAB-uni-32R | x35 | 96455 |
| | | | NH3_Uni35 | 1 | ORNL-HgcAB-uni-R | x35 | 61379 |
| | | NH4 | NH4_Alt30 | 1 | ORNL-HgcAB-uni-32R | x30 | 58452 |
| | | | NH4_Uni30 | 1 | ORNL-HgcAB-uni-R | x30 | 46964 |
| | | NH5 | NH5_Alt30 | 1 | ORNL-HgcAB-uni-32R | x30 | 80990 |
| | | | NH5_Uni30 | 1 | ORNL-HgcAB-uni-R | x30 | 44168 |
| | | NH6 | NH6_Alt30 | 1 | ORNL-HgcAB-uni-32R | x30 | 24519 |
| | | | NH6_Uni30 | 1 | ORNL-HgcAB-uni-R | x30 | 20762 |
| | | NH-H | NH-H-Alt30 | 4 | ORNL-HgcAB-uni-32R | X30 | 68046 |
| | | NH-I | NH-I-Alt30 | 1 | ORNL-HgcAB-uni-32R | X30 | 22275 |
| | | NH-J | NH-J-Alt30 | 1 | ORNL-HgcAB-uni-32R | X30 | 40817 |
| | | NH-K | NH-K-Alt30 | 4 | ORNL-HgcAB-uni-32R | X30 | 102968 |
| Environmental + spike | 1064 + mock community 1 spike | 1064_combo 1 | m1064_1combo_Alt30 | 4 | ORNL-HgcAB-uni-32R | x30 | 103803 |
| | | | m1064_1combo_Alt35 | 2 | ORNL-HgcAB-uni-32R | x35 | 93182 |
| | | | m1064_1combo_Uni30 | 5 | ORNL-HgcAB-uni-R | x30 | 231095 |
| | | | m1064_1combo_Uni35 | 2 | ORNL-HgcAB-uni-R | x35 | 141747 |
| | 1064 + mock community 2 spike | 1064_combo 2 | m1064_2combo_Alt30 | 5 | ORNL-HgcAB-uni-32R | x30 | 133057 |
| | | | m1064_2combo_Alt35 | 6 | ORNL-HgcAB-uni-32R | x35 | 656352 |
| | | | m1064_2combo_Uni30 | 5 | ORNL-HgcAB-uni-R | x30 | 88372 |
| | | | m1064_2combo_Uni35p 1 | 5 | ORNL-HgcAB-uni-R | x35 | 290878 |
| Mock community | Mock community 1 | combo1 | m1combo_alt30 | 5 | ORNL-HgcAB-uni-32R | x30 | 284603 |
| | | | m1combo_uni30 | 6 | ORNL-HgcAB-uni-R | x30 | 235704 |
| | Mock community 2 | combo2 | m2combo_Alt30p1 | 5 | ORNL-HgcAB-uni-32R | x30 | 546866 |
| | | | m2combo_Uni30p1n1 | 6 | ORNL-HgcAB-uni-R | x30 | 322774 |

+ total number of biological and sequencing replicates included in analyses

**Supplementary Table 5.** Occurrence of reverse oligo sequences in reference *hgcAB* sequences (n = 239) and percent occurrence in environmental *hgcAB* clones (n = 369), and amplicons (n = 233934) amplified by ORNL-HgcAB-uni-F and ORNL-HgcAB-uni-32R.

| ORNL-HgcAB-uni-32R oligo | Primer Sequence | Reference *hgcB* database (n=239) | | | % in environmental clones (n = 369) | % in environmental amplicons (n = 233934) |
|---|---|---|---|---|---|---|
| | | 0-mismatch | 1-mismatch | 2-mismatches | | |
| 1 | CAGGCACCGCATTCCATACA | 3 | 18 | 82 | 0.5 | 0.6 |
| 2 | CAGGCACCGCATTCCATGCA | 4 | 46 | 126 | 3.5 | 2.4 |
| 3 | CAGGCACCGCATTCGATACA | 0 | 7 | 35 | 0.8 | 1.2 |
| 4 | CAGGCACCGCATTCGATGCA | 2 | 12 | 58 | 4.6 | 3.3 |
| 5 | CAGGCACCGCACTCCATACA | 1 | 13 | 84 | 0.0 | 0.9 |
| 6 | CAGGCACCGCACTCCATGCA | 0 | 51 | 118 | 0.5 | 2.5 |
| 7 | CAGGCACCGCACTCGATACA | 0 | 1 | 21 | 0.0 | 1.3 |
| 8 | CAGGCACCGCACTCGATGCA | 0 | 4 | 60 | 1.4 | 4.2 |
| 9 | CAGGCTCCGCATTCCATACA | 1 | 20 | 82 | 0.0 | 0.6 |
| 10 | CAGGCTCCGCATTCCATGCA | 6 | 41 | 126 | 4.3 | 2.8 |
| 11 | CAGGCTCCGCATTCGATACA | 0 | 5 | 32 | 0.5 | 1.2 |
| 12 | CAGGCTCCGCATTCGATGCA | 0 | 11 | 55 | 7.6 | 4.3 |
| 13 | CAGGCTCCGCACTCCATACA | 0 | 11 | 89 | 0.5 | 0.8 |
| 14 | CAGGCTCCGCACTCCATGCA | 1 | 54 | 109 | 3.3 | 2.8 |
| 15 | CAGGCTCCGCACTCGATACA | 0 | 0 | 17 | 0.5 | 1.3 |
| 16 | CAGGCTCCGCACTCGATGCA | 0 | 1 | 61 | 3.3 | 5.0 |
| 17 | CAGGCCCCGCATTCCATACA | 4 | 27 | 108 | 0.0 | 0.7 |
| 18 | CAGGCCCCGCATTCCATGCA | 13 | 71 | 129 | 4.6 | 2.5 |
| 19 | CAGGCCCCGCATTCGATACA | 0 | 7 | 40 | 0.8 | 1.2 |
| 20 | CAGGCCCCGCATTCGATGCA | 2 | 19 | 79 | 5.1 | 3.9 |
| 21 | CAGGCCCCGCACTCCATACA | 4 | 39 | 102 | 0.0 | 0.9 |
| 22 | CAGGCCCCGCACTCCATGCA | 22 | 69 | 123 | 2.4 | 2.9 |
| 23 | CAGGCCCCGCACTCGATACA | 0 | 4 | 44 | 2.2 | 1.5 |
| 24 | CAGGCCCCGCACTCGATGCA | 0 | 24 | 77 | 3.5 | 5.3 |
| 25 | CAGGCGCCGCATTCCATACA | 1 | 33 | 108 | 2.7 | 1.1 |
| 26 | CAGGCGCCGCATTCCATGCA | 14 | 70 | 133 | 10.3 | 5.8 |
| 27 | CAGGCGCCGCATTCGATACA | 1 | 4 | 45 | 4.3 | 2.5 |
| 28 | CAGGCGCCGCATTCGATGCA | 1 | 21 | 84 | 10.6 | 10.0 |
| 29 | CAGGCGCCGCACTCCATACA | 3 | 35 | 99 | 0.5 | 1.5 |
| 30 | CAGGCGCCGCACTCCATGCA | 22 | 66 | 120 | 10.0 | 6.9 |
| 31 | CAGGCGCCGCACTCGATACA | 0 | 6 | 38 | 3.3 | 3.1 |
| 32 | CAGGCGCCGCACTCGATGCA | 0 | 23 | 79 | 7.3 | 15.1 |
| | % of sequences recovered | 56.3% | 96.9% | 100% | 84.4% | 100% |

**Supplementary Table 6**. Theoretical binding efficiencies of reverse primers

| | In silico binding to *hgcAB* reference database (n = 239) | | | | | | | | In silico binding to Ferredoxin database (n = 14,161) | | | |
|---|---|---|---|---|---|---|---|---|---|---|---|---|
| | 0 mismatches | | | | 2 mismatches allowed | | | | 0 mismatches | | 2 mismatches | |
| Primer Version | D*. | F*. | M*. | All* | D*. | F*. | M*. | All* | *hgcB* | Non-*hgcB* | *hgcB* | Non-*hgcB* |
| ORNL-HgcAB-uni-32R | 0.71 | 0.26 | 0.38 | 0.50 | 0.98 | 0.94 | 0.88 | 0.98 | 0.61 | 0 | 1.00 | 0.04 |
| ORNL-HgcAB-uni-R | 0.88 | 0.45 | 0.88 | 0.75 | 0.98 | 1.00 | 0.88 | 0.98 | 0.90 | 0 | 1.00 | 0.03 |

*Theoretical binding efficiencies (%) of each reverse primer to a reference database of 239 *hgcB* nucleotide sequences, including matches to *Deltaproteobacteria* (D), *Firmicutes* (F), *Methanomicrobia* (M), and to a 4Fe-4S ferredoxin reference database that contained 88 *hgcB* sequences and 14,1073 non-*hgcB* sequences.

**Supplementary Table 7.** All PCR conditions tested for optimal primer to template ratio for amplification of *hgcAB* from environmental samples.

| Condition | Template concentration (final) | Primer concentration (final) | Reaction Volume | Primer Set |
|---|---|---|---|---|
| A[+] | 0.5 ng/µl | 1.0 µM | 20 µl | 1. ORNL-HgcAB-uni-F & ORNL-HgcAB-uni-R<br>2. ORNL-HgcAB-uni-F & ORNL-HgcAB-32-R<br>3. ORNL-HgcAB-uni-F & Equimolar 26<br>4. ORNL-HgcAB-uni-F & Equimolar 32 |
| B* | 0.5 ng/µl | 0.5 µM | 20 µl | 1. ORNL-HgcAB-uni-F & ORNL-HgcAB-uni-R<br>2. ORNL-HgcAB-uni-F & ORNL-HgcAB-32-R<br>3. ORNL-HgcAB-uni-F & Equimolar 26<br>4. ORNL-HgcAB-uni-F & Equimolar 32 |
| C | 0.002 ng/µl | 0.5 µM | 50 µl | 1. ORNL-HgcAB-uni-F & ORNL-HgcAB-uni-R<br>2. ORNL-HgcAB-uni-F & ORNL-HgcAB-32-R<br>3. ORNL-HgcAB-uni-F & Equimolar 26<br>4. ORNL-HgcAB-uni-F & Equimolar 32 |
| D | 0.002 ng/µl | 1.0 µM | 50 µl | 1. ORNL-HgcAB-uni-F & ORNL-HgcAB-uni-R<br>2. ORNL-HgcAB-uni-F & ORNL-HgcAB-32-R<br>3. ORNL-HgcAB-uni-F & Equimolar 26<br>4. ORNL-HgcAB-uni-F & Equimolar 32 |
| E | 0.02 ng/µl | 1.0 µM | 50 µl | 1. ORNL-HgcAB-uni-F & ORNL-HgcAB-uni-R<br>2. ORNL-HgcAB-uni-F & ORNL-HgcAB-32-R<br>3. ORNL-HgcAB-uni-F & Equimolar 26<br>4. ORNL-HgcAB-uni-F & Equimolar 32 |

[+]Condition from Christensen et al. (2016)

*Condition used to amplify *hgcAB* for cloning experiment
